# Validation of Enhancer Regions in Primary Human Neural Progenitor Cells using Capture STARR-seq

**DOI:** 10.1101/2024.03.14.585066

**Authors:** Sophia C. Gaynor-Gillett, Lijun Cheng, Manman Shi, Jason Liu, Gaoyuan Wang, Megan Spector, Mary Flaherty, Martha Wall, Ahyeon Hwang, Mengting Gu, Zhanlin Chen, Yuhang Chen, PsychENCODE Consortium, Jennifer R. Moran, Jing Zhang, Donghoon Lee, Mark Gerstein, Daniel Geschwind, Kevin P. White

## Abstract

Genome-wide association studies (GWAS) and expression analyses implicate noncoding regulatory regions as harboring risk factors for psychiatric disease, but functional characterization of these regions remains limited. We performed capture STARR-sequencing of over 78,000 candidate regions to identify active enhancers in primary human neural progenitor cells (phNPCs). We selected candidate regions by integrating data from NPCs, prefrontal cortex, developmental timepoints, and GWAS. Over 8,000 regions demonstrated enhancer activity in the phNPCs, and we linked these regions to over 2,200 predicted target genes. These genes are involved in neuronal and psychiatric disease-associated pathways, including dopaminergic synapse, axon guidance, and schizophrenia. We functionally validated a subset of these enhancers using mutation STARR-sequencing and CRISPR deletions, demonstrating the effects of genetic variation on enhancer activity and enhancer deletion on gene expression. Overall, we identified thousands of highly active enhancers and functionally validated a subset of these enhancers, improving our understanding of regulatory networks underlying brain function and disease.

## Introduction

Psychiatric disorders are among the most common illnesses in the United States, with over 20% of adults being afflicted (*1*). Additionally, 20-25% of children in the United States currently have, or have had in their lifetime, a serious mental illness (*2*). The causes of mental illness are complex and include both genetic and environmental components such as stressful life events, brain damage, or childhood neglect (*3*). To better understand the genetic contributors to psychiatric disease, hundreds of genome-wide association studies (GWAS) have been conducted (*4*) to identify common genetic variants associated with disease. Over a thousand psychiatric disease-associated variants have been identified through GWAS, and the majority of these implicated variants occur in noncoding parts of the genome (*4, 5*). Additionally, studies of genome-wide gene expression have identified hundreds of candidate genes that are differentially regulated in certain brain regions between case and control cohorts (*6–9*). Many of these expression level changes may have genetic or epigenetic roots in noncoding regulatory element activity (*10–13*).

A major goal of the PsychENCODE Consortium is to identify and characterize noncoding regulatory elements in the human brain, both normal and diseased, in an effort to understand the role of these noncoding regions. Since its first publication (*14*), the PsychENCODE Consortium has compiled an extensive online resource of data from postmortem human brain samples, including developing and adult brains as well as normal and diseased brains (https://psychencode.synapse.org/). Through a multi-omics approach, PsychENCODE has endeavored to develop a regulatory map of noncoding elements in the human brain. Following on this effort, human cell line models have been utilized to functionally validate noncoding regulatory elements identified through these large-scale omics approaches. Towards that end, the current study utilized primary human neural progenitor cells (phNPCs) obtained directly from fetal brain. These phNPCs have been shown to closely recapitulate early stages of *in vivo* human fetal brain development, even more so than human induced pluripotent stem cell (hiPSC)-derived NPCs or human embryonic stem cell (hESC)-derived NPCs (*15, 16*). These primary cells are a valuable model system for investigating the potential role of noncoding regions in brain development and function.

In the current study, we leveraged several PsychENCODE datasets, including data from the prefrontal cortex (PFC), NPC lines, and developmental timepoints to generate a list of putative enhancer regions to investigate using the phNPC model. Many of the noncoding variants implicated in psychiatric disease have been associated with enhancer regions (*17–19*), which are a specific type of noncoding regulatory element activating gene expression. To investigate these enhancers, we performed capture self-transcribing active regulatory region sequencing (CapSTARR-seq). The CapSTARR-seq method is a plasmid-based, *in vitro* approach for large-scale validation of enhancer regions (*20*). Our lab has previously applied the STARR-seq approach on a genome-wide scale to a number of cell lines (*21*), including the SH-SY5Y neuroblastoma cell line (https://doi.org/doi:10.17989%2FENCSR983SZZ). With a primary cell line like phNPCs, however, the cell counts necessary for whole-genome STARR-seq, usually in the hundreds of millions, are not feasible. As an alternative approach, CapSTARR-seq begins with a hybridization-based capture method to isolate candidate enhancer regions of interest from sheared genomic DNA (*22*). By targeting a more limited number of regions, CapSTARR-seq requires fewer cells and can be used in cell lines that are difficult to expand to very large numbers, like phNPCs.

In this study, we identified over 8,000 regions that demonstrated enhancer activity in the phNPCs. Notably, we found that these active enhancer regions were enriched for binding site motifs of transcription factors with high expression in the phNPCs and genetic variants implicated in psychiatric disease. Additionally, the target genes for these enhancer regions are strongly enriched for neuronal pathways. For a subset of these enhancer regions, we used CRISPR-based deletions to demonstrate the effect of the enhancers on gene expression and a mutation STARR-seq approach (MutSTARR-seq) to show that introducing genetic variation in these enhancer regions affects their enhancer activity.

## Results

### Data quality

We used two separate panels of CapSTARR-seq to test the enhancer activity of over 78,000 putative enhancer regions in phNPCs (**Fig. 1A-C**). For the first panel, we targeted 22,400 regions based on ATAC-seq, DNase-seq, Hi-C, and ChIP-seq data from the PFC (**Fig. 1A**; Materials and Methods). The second panel targeted 56,215 regions based on data from NPCs, PFC, developmental timepoints, and psychiatric disease-associated GWAS (**Fig. 1B**; Materials and Methods). Initial quality control revealed that our sequencing data was of high quality (Table S1), with over 94% of reads from each panel and replicate aligning with the genome. The rates of polymerase chain reaction (PCR) duplication ranged from 11.83% to 49.30% across all sequencing data, with the highest levels of duplication coming from the Panel 2 output data. The higher level of duplicates in Panel 2, particularly of barcoded duplicates, represents high levels of enhancer activity and indicates that our panel was well-designed. We also calculated “on-target” and “off-target” read percentages based on whether the reads fell within our initial target regions. Over 89% of our sequencing reads qualified as “on-target” (Table S1).

**Figure 1.**
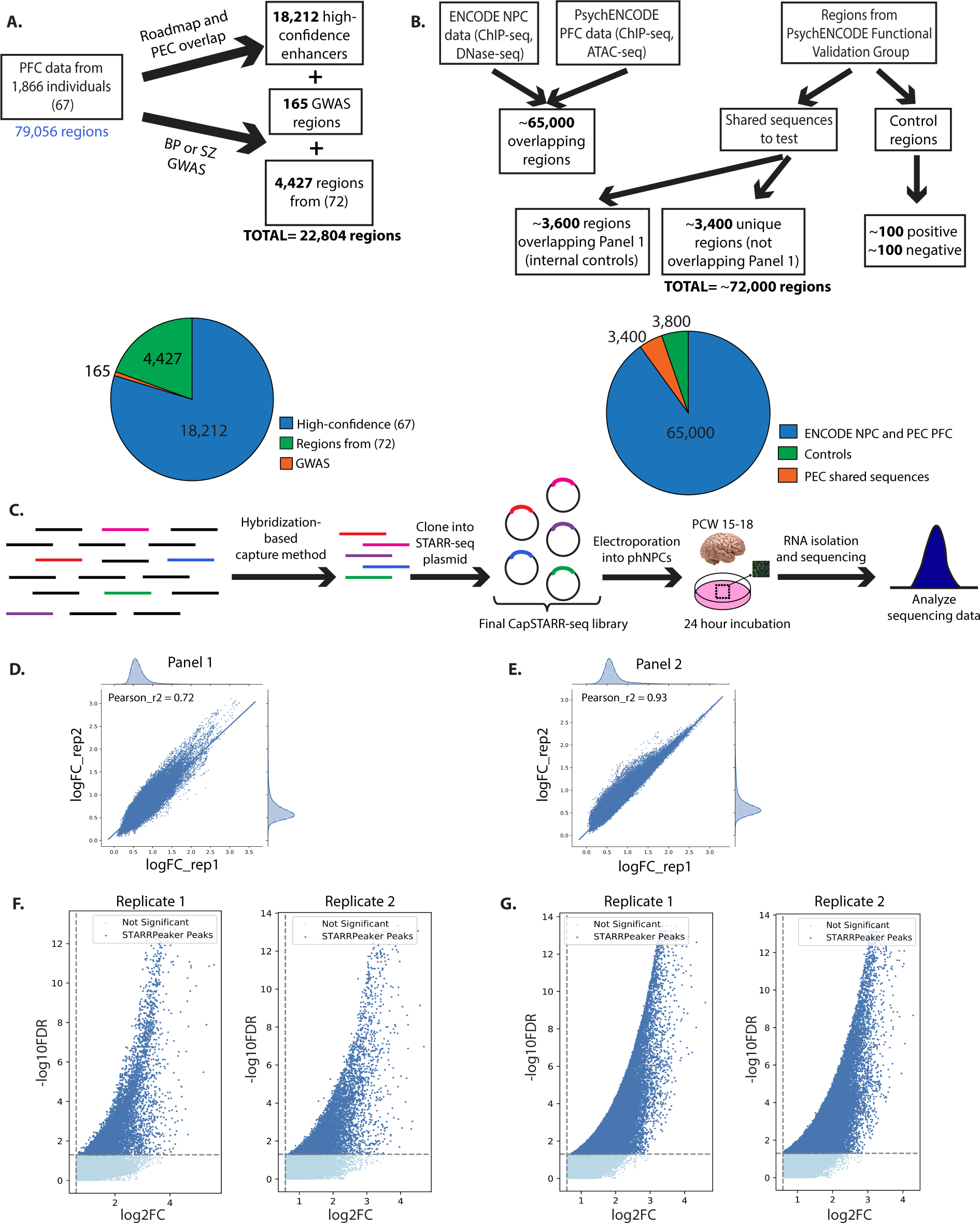
Panel design and quality control results. (**A**) and (**B**) show the process for candidate enhancer selection for Panel 1 (**A**) and Panel 2 (**B**). See Materials and Methods for a complete description of candidate selection. (**C**) demonstrates the experimental workflow. Sheared genomic DNA was hybridized to probes specific for the candidate enhancer regions. These regions were then cloned into the STARR-seq plasmid and transfected into phNPCs. (**D**) and (**E**) show the fold change correlation between the two technical replicates for Panel 1 (**D**) and Panel 2 (**E**). Pearson r^2^ values are included on the graphs. (**F**) and (**G**) are volcano plots representing the tested enhancer regions. Regions that had significant peaks as determined by STARRPeaker are in dark blue while non-significant regions are in light blue. Abbreviations: PFC = prefrontal cortex; PEC = PsychENCODE Consortium; BP = bipolar disorder; SZ = schizophrenia; GWAS = genome-wide association study; NPC = neural progenitor cell; FDR = false discovery rate; FC = fold change.

### Peak calling

We used STARRPeaker (*23*) to call peaks from our dataset representing active enhancer regions. From the first panel, we identified 1,137 and 1,142 regions from replicates 1 and 2 respectively that demonstrated active enhancer activity (**Table 1**, Table S2). Of these regions, 914 overlapped, representing an r^2^ value of 0.72 between replicates (**Figures 1D, 1F**). For the second panel, we identified 6,202 and 6,484 regions from replicates 1 and 2 respectively that demonstrated active enhancer activity (**Table 1**). For Panel 2, 5,698 regions overlapped between replicates 1 and 2, producing an r^2^ value of 0.93 between replicates (**Figures 1E, 1G**). Across both panels, we identified 8,148 regions with evidence of enhancer activity in at least one replicate. A total of 6,612 regions (∼8.4%) demonstrated strong evidence of enhancer activity in the phNPCs based on their replication in two separate experiments. This percentage of strongly enriched regions is similar to the percentage seen in previous CapSTARR-seq experiments in non-neuronal cell lines (∼6%; (*22*)).

**Table 1.** Active enhancer regions across each CapSTARR-seq panel. Active enhancer regions are designated as regions with a STARRPeaker q-value ≤ 0.05. The overlap value indicates how many regions were designated active enhancers across both technical replicates of the panel. The Pearson r^2^ value (also depicted in **Figures 1D** and **1E**) demonstrates the correlation between replicates.

| | Replicate 1 | Replicate 2 | Overlap | Pearson $r^2$ Value |
| --- | --- | --- | --- | --- |
| **Panel 1** | 1,137 | 1,142 | 914 | 0.72 |
| **Panel 2** | 6,202 | 6,484 | 5,698 | 0.93 |

### Enrichment of transcription factor binding site motifs

To determine whether our active enhancer regions were enriched for transcription factor binding sites (TFBS), we searched for enrichment of binding motifs for several common transcription factors (TFs; see Materials and Methods) focusing on the high confidence enhancers that overlapped between the panel replicates (**Table 2**). We compared enrichment in our active enhancer regions with control regions that showed no enhancer activity in our CapSTARR-seq assay. In Panel 1, the top enriched motifs were JunB/bZip (p = 1×10^-61^), TP53 (p = 1×10^-29^), MITF (p = 1×10^-21^), and SOX10 (p = 1×10^-17^). In Panel 2, the top enriched motifs were YY1 (p = 1×10^-609^), ELK1/ETS (p = 1×10^-490^), THAP11 (p = 1×10^-232^), SREBF2 (p = 1×10^-200^), and ZNF143 (p = 1×10^-139^).

**Table 2.**
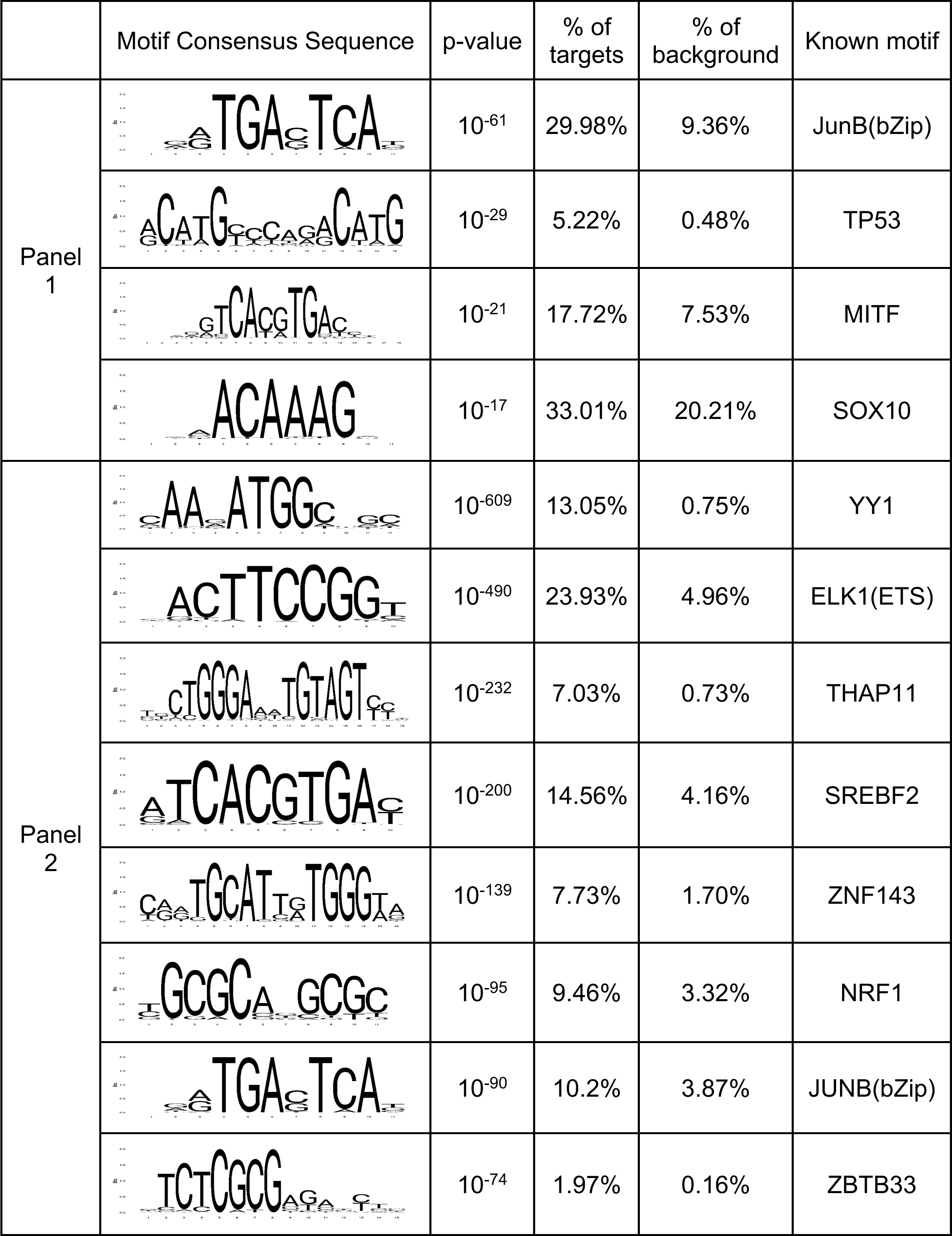
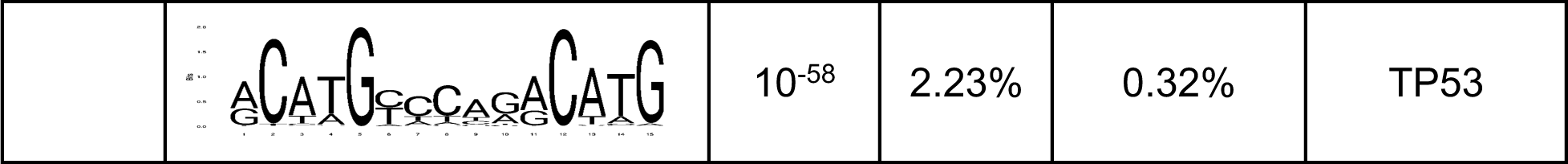
Transcription factor binding site motifs enriched in active enhancers. The “% of targets” column indicates the percentage of active enhancers from our CapSTARR-seq assay that contain each motif. The “% of background” indicates the percentage of regions from our CapSTARR-seq assay that did not show enhancer activity and contain each motif (see Materials and Methods). The known motif column indicates the transcription factors known to bind to the given motif sequence.

|  | Motif Consensus Sequence | p-value | % of targets | % of background | Known motif |
| --- | --- | --- | --- | --- | --- |
| Panel 1 | | $10^{-61}$ | 29.98% | 9.36% | JunB(bZip) |
| | | $10^{-29}$ | 5.22% | 0.48% | TP53 |
| | | $10^{-21}$ | 17.72% | 7.53% | MITF |
| | | $10^{-17}$ | 33.01% | 20.21% | SOX10 |
| Panel 2 | | $10^{-609}$ | 13.05% | 0.75% | YY1 |
| | | $10^{-490}$ | 23.93% | 4.96% | ELK1(ETS) |
| | | $10^{-232}$ | 7.03% | 0.73% | THAP11 |
| | | $10^{-200}$ | 14.56% | 4.16% | SREBF2 |
| | | $10^{-139}$ | 7.73% | 1.70% | ZNF143 |
| | | $10^{-95}$ | 9.46% | 3.32% | NRF1 |
| | | $10^{-90}$ | 10.2% | 3.87% | JUNB(bZip) |
| | | $10^{-74}$ | 1.97% | 0.16% | ZBTB33 |
| | | $10^{-58}$ | 2.23% | 0.32% | TP53 |

Several of these TFs have previous implications in pathways related to brain development and psychiatric disease, including glial development (SOX10; (*24, 25*)), drug addiction (JunB; (*26*)), and schizophrenia (SOX10, SREBF2; (*27, 28*)).

To further validate that the enriched TFBS motifs correlate with TFs that are highly expressed in phNPCs, we analyzed their expression in previously published microarray data from the phNPCs (*15*), taking into account similar motifs matching the enriched sequences. Motifs matching the same target sequence are referred to as a motif family. We compared microarray expression levels of the TFs from the four high-confidence motif families in Panel 1 and the nine in Panel 2 (**Table 2**) with 100 randomly selected TFs from Lambert et al. (see Materials and Methods; (*29*)). In Panel 1, the motif-enriched TFs showed a trend towards higher expression than the randomly selected TFs, but this did not reach significance (p = 0.211; **Figure 2**, Tables S3-S5).

**Figure 2.**
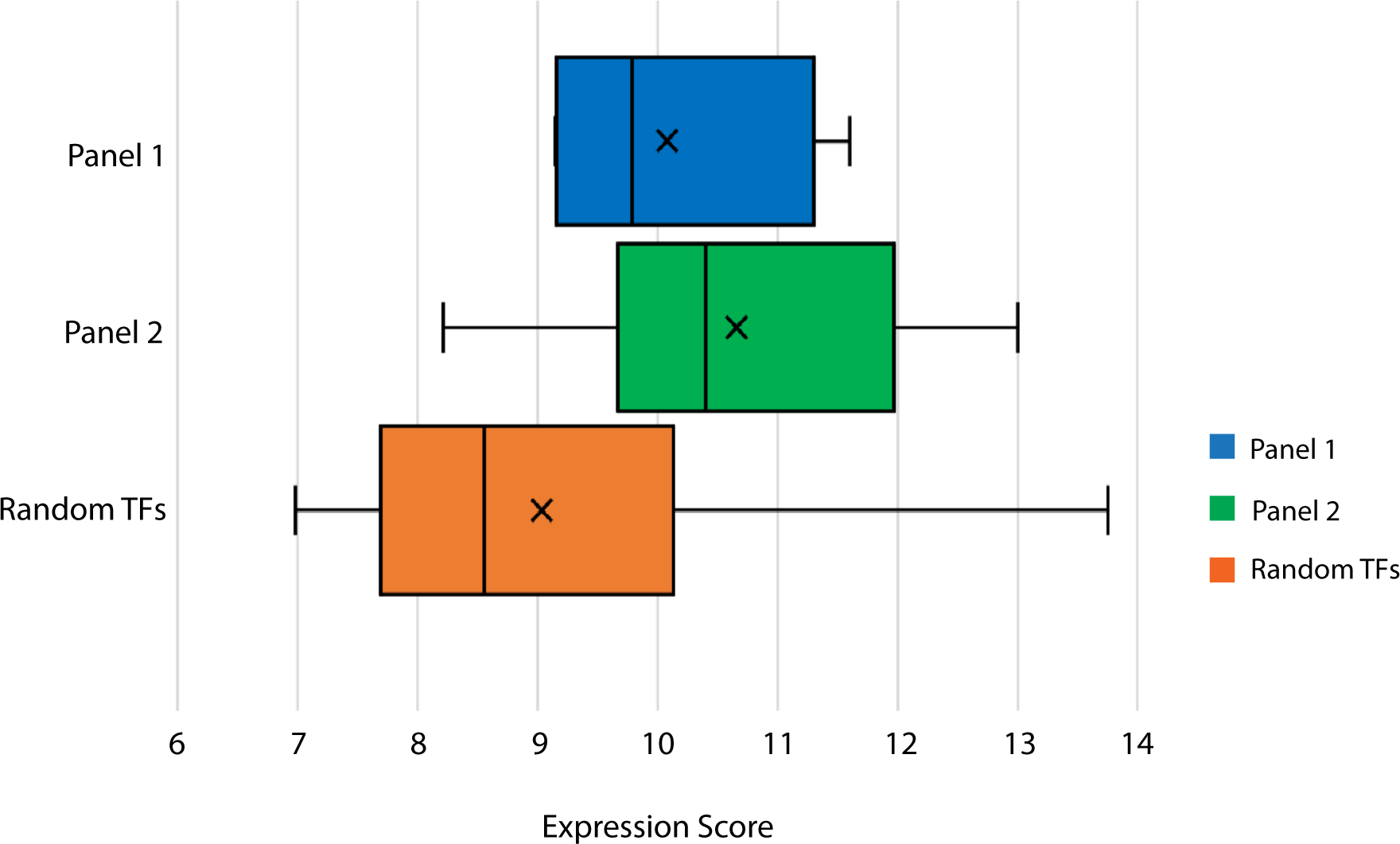
Expression of TFs with enriched binding site motifs. Expression score is displayed on the X-axis. A higher expression score indicates a more highly expressed gene. For each plot, the solid line represents the median expression score while the “X” represents the mean expression score. Expression scores were compared using a two-tailed t-test. The expression of motif-enriched TFs from Panel 1 did not differ significantly from the expression of random TFs (p = 0.211). The expression of motif-enriched TFs from Panel 2 was significantly higher than the expression of random TFs (p = 0.0052).

The same comparison with Panel 2 showed significantly higher expression than the randomly selected TFs (p = 0.0052; **Figure 2**, Tables S3-S5). These results demonstrate that the TFs associated with enriched binding site motifs from our active enhancer regions have higher expression in phNPCs than randomly selected TFs.

### Gene-enhancer linkage

From the set of 8,148 putative enhancers, we first used adult brain data (*30*) and identified 2,288 unique predicted target genes regulated by 427 TFs (Table S6; Materials and Methods). We used the following pathway analysis tools to identify biological pathways enriched within our gene set: DAVID (https://david.ncifcrf.gov/; (*31*)), Enrichr (https://maayanlab.cloud/Enrichr/; (*32*)), and STRING (https://string-db.org/; (*33*)). These databases identified highly overlapping pathways, which is expected given their redundancy. Many of the most highly enriched pathways were related to neuronal processes, including generation of neurons, synapse organization, axonogenesis, axon guidance, glutamatergic synapse, dopaminergic synapse, GABAergic synapse, and synaptic vesicle cycle (**Tables 3-4**, **Figure 3A-C**). These databases also examine disease-related pathways, and many of the most highly enriched diseases were brain-related. Enriched disease pathways included nervous system disease, neurodegenerative disease, and Alzheimer’s disease (**Tables 3-4**, **Figure 3D**). To ensure that these pathways were not enriched as an artifact of our candidate enhancer selection method, we ran the same set of pathway analyses on randomly selected subsets of genes from our entire candidate list (see Materials and Methods). We found that many of the identified pathways were significantly more enriched for our CapSTARR-seq gene set than for randomly selected gene subsets (Table S7).

**Figure 3.**
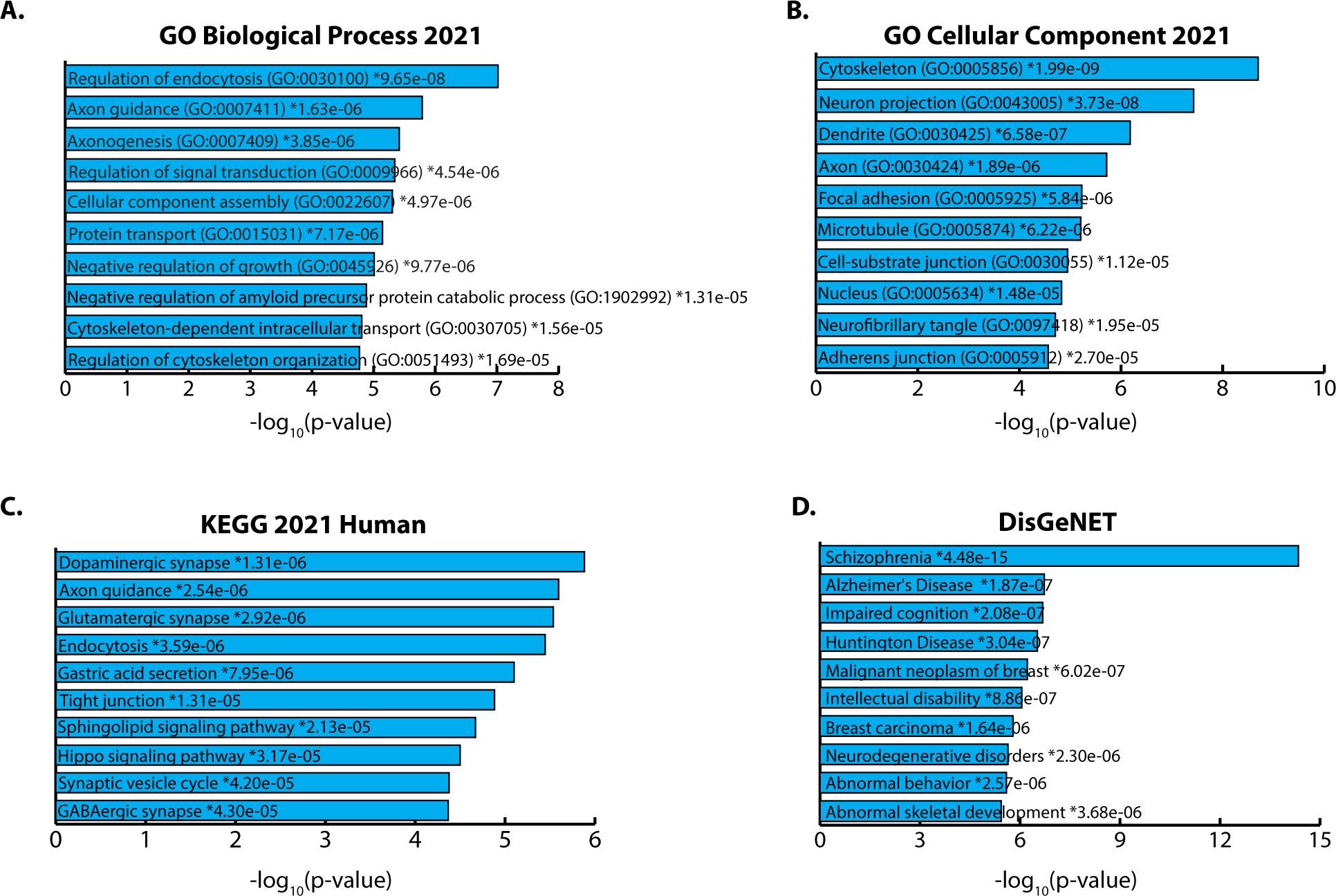
Pathway analysis results from Enrichr. (**A**) and (**B**) depict results from the gene ontology (GO) knowledgebase - biological process (**A**) and cellular component (**B**). (**C**) depicts results from the KEGG database. (**D**) depicts results from DisGeNET examining gene-disease associations. The p-values for each category are included on the bars for each category. The asterisks (*) indicate that the adjusted p-value for that category is also significant (<0.05). Enrichr calculates p-values using the Fisher exact test and adjusted p-values using the Benjamini-Hochberg method.

**Table 3.** DAVID pathway analysis of predicted target genes. Table includes the top 10 results for the GOTERM_BP_DIRECT and KEGG_PATHWAY categories and all results with a p-value < 0.05 for the GAD_DISEASE_CLASS category. The “Count” column indicates the number of genes from our set present in that specific pathway. DAVID calculates p-values using the Fisher exact test and false discovery rates (FDR) using the Benjamini-Hochberg method.

| Category | Term | Count | P-Value | Benjamini FDR |
| --- | --- | --- | --- | --- |
| GOTERM_BP_DIRECT | Regulation of catalytic activity | 73 | 1.30E-06 | 3.50E-03 |
| GOTERM_BP_DIRECT | Signal transduction | 192 | 1.80E-06 | 3.50E-03 |
| GOTERM_BP_DIRECT | Nervous system development | 78 | 2.20E-06 | 3.50E-03 |
| GOTERM_BP_DIRECT | Intracellular signal transduction | 83 | 2.40E-06 | 3.50E-03 |
| GOTERM_BP_DIRECT | Axonogenesis | 26 | 6.30E-06 | 7.30E-03 |
| GOTERM_BP_DIRECT | Positive regulation of endocytosis | 12 | 1.00E-05 | 9.70E-03 |
| GOTERM_BP_DIRECT | Axon guidance | 41 | 1.40E-05 | 1.10E-02 |
| GOTERM_BP_DIRECT | Regulation of small GTPase mediated signal transduction | 30 | 1.50E-05 | 1.10E-02 |
| GOTERM_BP_DIRECT | Synapse organization | 19 | 2.20E-05 | 1.40E-02 |
| GOTERM_BP_DIRECT | Cytoskeleton-dependent intracellular transport | 10 | 7.30E-05 | 4.20E-02 |
| KEGG_PATHWAY | Axon guidance | 43 | 1.50E-05 | 1.80E-03 |
| KEGG_PATHWAY | Glutamatergic synapse | 31 | 1.80E-05 | 1.80E-03 |
| KEGG_PATHWAY | Endocytosis | 54 | 1.90E-05 | 1.80E-03 |
| KEGG_PATHWAY | Dopaminergic synapse | 34 | 2.20E-05 | 1.80E-03 |
| KEGG_PATHWAY | Gastric acid secretion | 23 | 4.90E-05 | 3.20E-03 |
| KEGG_PATHWAY | Tight junction | 39 | 7.00E-05 | 3.30E-03 |
| KEGG_PATHWAY | Hippo signaling pathway | 37 | 7.00E-05 | 3.30E-03 |
| KEGG_PATHWAY | Sphingolipid signaling pathway | 30 | 1.10E-04 | 4.60E-03 |
| KEGG_PATHWAY | GABAergic synapse | 24 | 2.20E-04 | 6.40E-03 |
| KEGG_PATHWAY | Synaptic vesicle cycle | 22 | 2.20E-04 | 6.40E-03 |
| GAD_DISEASE_CLASS | Chemdependency | 525 | 5.90E-07 | 1.10E-05 |
| GAD_DISEASE_CLASS | Psych | 298 | 1.40E-05 | 1.30E-04 |
| GAD_DISEASE_CLASS | Pharmacogenomic | 391 | 5.00E-05 | 3.00E-04 |
| GAD_DISEASE_CLASS | Neurological | 387 | 1.10E-02 | 5.10E-02 |
| GAD_DISEASE_CLASS | Cardiovascular | 552 | 2.50E-02 | 9.10E-02 |
| GAD_DISEASE_CLASS | Developmental | 204 | 4.00E-02 | 1.20E-01 |

**Table 4.** STRING pathway analysis of predicted target genes. Table includes the top 10 results for the Gene Ontology (GO) Biological Process, KEGG Pathways, and Reactome Pathways categories and all results for the DISEASES category. The “# Genes” column indicates the number of genes from our set present in that specific pathway. STRING calculates false discovery rates (FDR) using the Benjamini-Hochberg method.

| Category | Description | # Genes | FDR Value |
| --- | --- | --- | --- |
| Diseases | Central nervous system disease | 191 | 1.70E-03 |
| Diseases | Neurodegenerative disease | 94 | 3.40E-03 |
| Diseases | Nervous system disease | 318 | 1.57E-02 |
| GO Biological Process | Biological regulation | 1630 | 2.55E-15 |
| GO Biological Process | Regulation of biological process | 1546 | 2.16E-14 |
| GO Biological Process | Nervous system development | 425 | 8.76E-14 |
| GO Biological Process | Regulation of cellular process | 1477 | 2.09E-13 |
| GO Biological Process | Neurogenesis | 309 | 4.29E-11 |
| GO Biological Process | Regulation of signaling | 567 | 5.56E-11 |
| GO Biological Process | Regulation of cell communication | 561 | 6.55E-11 |
| GO Biological Process | Cellular component organization | 801 | 4.98E-10 |
| GO Biological Process | Generation of neurons | 287 | 5.36E-10 |
| GO Biological Process | Cellular component organization or biogenesis | 823 | 5.57E-10 |
| KEGG Pathways | Endocytosis | 54 | 9.50E-03 |
| KEGG Pathways | Axon guidance | 42 | 9.50E-03 |
| KEGG Pathways | Glutamatergic synapse | 31 | 9.50E-03 |
| KEGG Pathways | Sphingolipid signaling pathway | 30 | 1.57E-02 |
| KEGG Pathways | Hippo signaling pathway | 36 | 1.57E-02 |
| KEGG Pathways | Tight junction | 37 | 1.57E-02 |
| KEGG Pathways | Synaptic vesicle cycle | 22 | 1.57E-02 |
| KEGG Pathways | GABAergic synapse | 24 | 1.57E-02 |
| KEGG Pathways | Dopaminergic synapse | 32 | 1.57E-02 |
| KEGG Pathways | Pathogenic E. coli infection | 42 | 1.57E-02 |
| Reactome Pathways | Neuronal system | 88 | 7.60E-04 |
| Reactome Pathways | Transmission across chemical synapses | 61 | 5.70E-03 |
| Reactome Pathways | Hemostasis | 110 | 1.34E-02 |
| Reactome Pathways | Neurotransmitter receptors and postsynaptic signal transmission | 47 | 1.68E-02 |
| Reactome Pathways | Membrane trafficking | 111 | 1.68E-02 |
| Reactome Pathways | Vesicle-mediated transport | 116 | 1.68E-02 |
| Reactome Pathways | Nervous system development | 104 | 1.68E-02 |
| Reactome Pathways | Signaling by receptor tyrosine kinases | 92 | 1.84E-02 |
| Reactome Pathways | Integration of energy metabolism | 29 | 3.40E-02 |
| Reactome Pathways | Transport of small molecules | 121 | 3.40E-02 |

Because these predicted genes were based on data from the adult brain, we also explored how our putative enhancers intersected with data from the fetal brain, which might be more relevant to regulatory relationships in phNPCs. We examined the overlap between our enhancer regions and fetal expression quantitative trait loci (eQTL) from Wen et al. (*34*). We found that, of our 8,148 putative enhancers, 2,284 (28%) overlapped a fetal eQTL while only 404 (5%) overlapped an adult eQTL (Table S8; (*30*)). Of the 404 that overlapped an adult eQTL, 300 also overlapped a fetal eQTL (Table S8). Of these 300 enhancers, 135 had the same predicted target gene and the same direction of effect on that target gene (Table S9). Pathway analyses of this set of overlapping genes largely implicated immune-related pathways (Table S10) using DAVID. Additionally, Enrichr identified autism spectrum disorder and schizophrenia as significantly enriched pathways for this gene set based on enrichment of single nucleotide polymorphisms (SNPs) from GWAS datasets (Fig. S1).

### MutSTARR-seq

To investigate the allelic effect of eQTLs on enhancer activity, we overlaid our CapSTARRseq results with eQTL data from 150 individuals with genotype and single nucleus RNA-seq (snRNA-seq) data intersecting the putative enhancer regions from adult to see if some enhancer regions were also active in adult brain (*30*). We selected 47 eQTLs to interrogate through MutSTARR-seq (Table S11; see Materials and Methods). MutSTARR-seq employs the same techniques as STARR-seq but utilizes synthetic gene fragments that are generated with and without the eQTL variant, which permits determination of the allelic effect of the eQTL variant on predicted enhancer activity. Through this analysis, we identified 4 variants that significantly altered the enhancer activity with the presence of the alternate allele (**Figure 4**, Table S12). For each of these regions, enhancer activity was increased from the alternate allele. However, the only region that survived correction for multiple testing was chr17:45,894,107-45,894,607 (adjusted p-value = 0.005).

**Figure 4.**
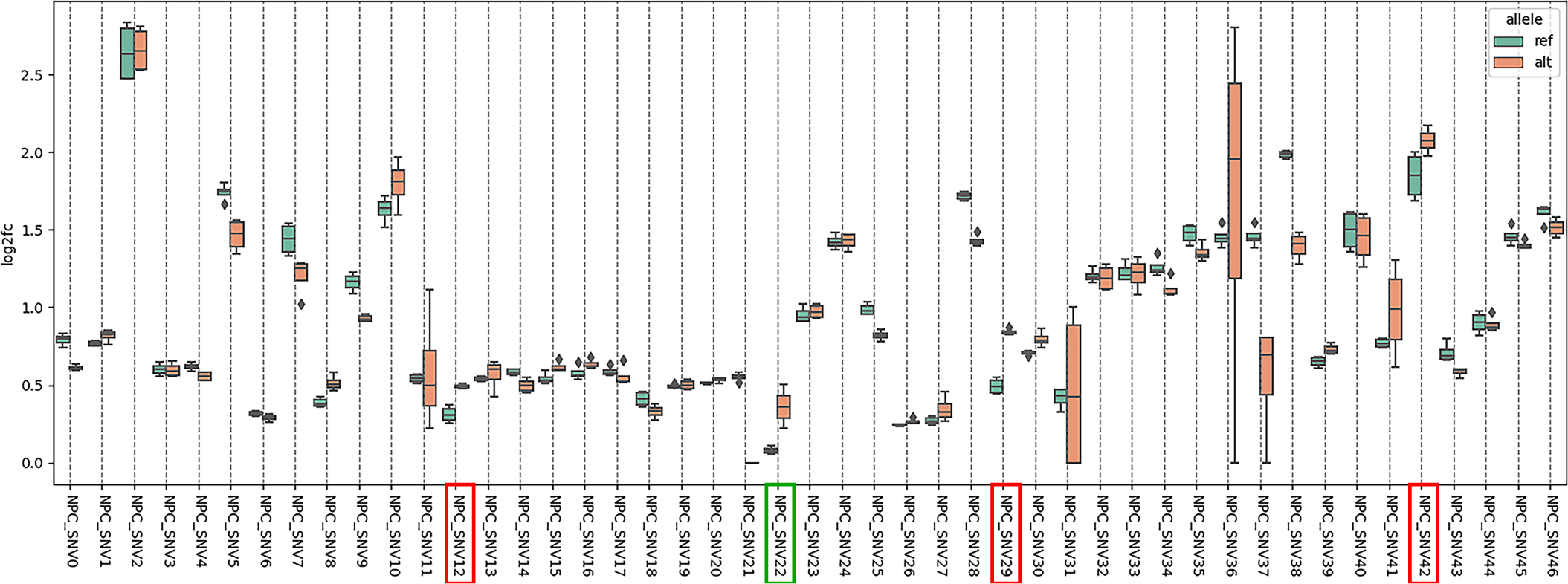
MutSTARR-seq results comparing enhancer activity (log2fc) between the reference allele (ref) and alternate allele (alt). Box plots represent the distribution of activity across four technical replicates. Enhancer activity is defined by log2 fold change, which represents the normalized output/input ratio in log2 space. Variants that had a nominally significant effect on enhancer activity (p < 0.05 before multiple testing correction) are boxed in red. P-values were calculated using a Chi-squared test. The variant that survived correction for multiple testing (p-value adjusted = 0.005) is boxed in green. Correction for multiple testing was done using Bonferroni and Benjamini-Hochberg methods. Abbreviations: log2fc = log2 fold change, NPC = neural progenitor cell, SNV = single nucleotide variant, ref = reference allele, alt = alternate allele.

To assess the validity of this finding, we compared it with the eQTL results used to generate our candidate list. Because our MutSTARR-seq results showed increased enhancer activity with the alternate allele, we expected to see increased expression of the target genes in individuals with the alternate allele. The predicted target genes for our significant region were *AC126544.2*, *ARL17A*, *ARL17B*, *CR936218.1*, *CRHR1*, *FAM215B*, *KANSL1-AS1, KANSL1 LRRC37A, LRRC37A2, MAPT-AS1,* and *MAPT*. Notably, for almost all genes and cell types examined, the target gene showed increased expression in individuals with the alternate allele (Table S13). The only genes that had decreased expression with the alternate allele were *ARL17A* and *MAPT-AS1.* This implicated enhancer region on chromosome 17 and its associated target genes fall within the 17q21.31 locus, a region of extremely high linkage disequilibrium (LD; (*35*)). This region has been extensively linked to a number of neurodegenerative diseases, including Alzheimer’s disease (*36*), Parkinson’s disease (*35, 37, 38*), frontotemporal dementia (*39*) and progressive supranuclear palsy (PSP; (*40, 41*)).

### Target gene expression level change after enhancer knockout (KO)

We chose for further functional analysis four active enhancers showing high enrichment in the CapSTARR-seq experiment and genetically associated with neuropsychiatric disorders. Ribonucleoprotein (RNP)-mediated CRISPR/Cas9 genome editing was used to delete candidate enhancers in phNPCs. We validated the editing by Sanger sequencing, which showed that cleavage occurred at the expected upstream and downstream CRISPR cut sites of all candidate enhancers tested (Fig. S2).

Densitometry analysis of genotyping PCR products showed genome editing KO efficiency ranging from roughly 19.69% to 44.97% for the enhancers tested (**Figure 5A-D**, Table S14). We then examined the relative expression level change of four nearby target genes: Neuronal guanine nucleotide exchange factor (*NGEF*), RAR related orphan receptor B (*RORB*), Pleckstrin homology domain containing O1 (*PLEKHO1*), and Target of Myb1 like 2 membrane trafficking protein (*TOM1L2*). Relative expression, measured by Taqman real-time quantitative PCR (qPCR) assay, showed that expression of all four target genes was diminished after enhancer KO: *NGEF* relative expression level was decreased to 0.45 (standard deviation (SD) = ± 0.01), *RORB* decreased to 0.56 (SD = ± 0.2), *PLEKHO1* decreased to 0.16 (SD = ± 0.02), *TOM1L2* decreased to 0.42 (SD = ± 0.01; **Figure 5A-D**, right).

**Figure 5.**
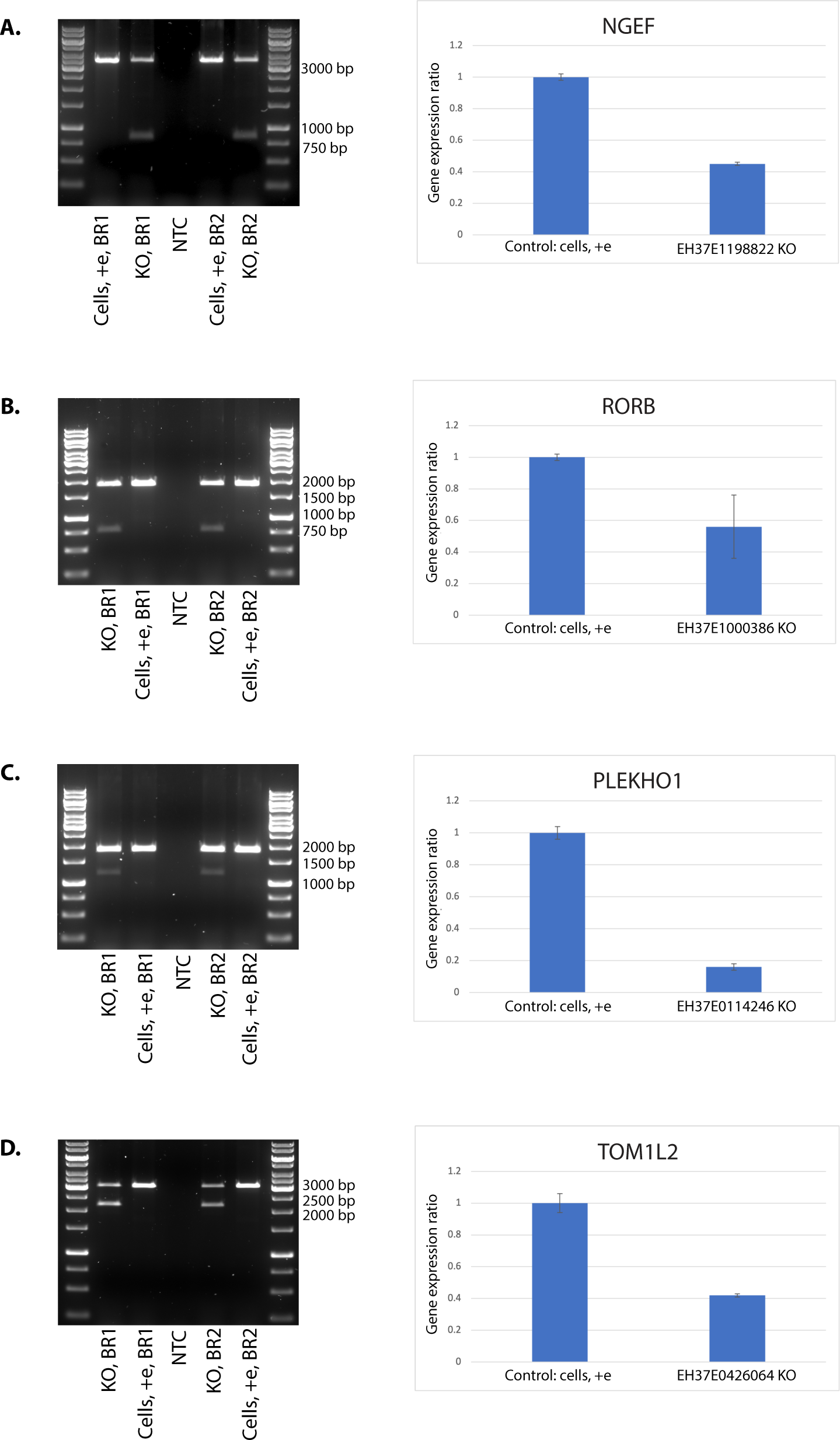
CRISPR/Cas9 enhancer knockout (KO). (**A-D**), left: DNA agarose gel image of the genotyping PCR results after KO of candidate enhancers in phNPCs through ribonucleoprotein (RNP)-mediated CRISPR/Cas9 genome editing. Control cells undergoing the same electroporation without any RNPs showed a clear strong WT band. For enhancer KO samples, besides the higher WT band, there is an additional clearly visible lower band in both BR1 and BR2 samples. The sizes of these lower bands are the same as the expected size of genome edited bands after enhancer KO. **(A-D)**, right: TaqMan qPCR Probe assay showed diminished expression level of the target gene after enhancer KO. C*_T_* values from triplicates were used to calculate the expression of the target gene relative to control cells using the Pfaffl method. Averages of the BR1 and BR2 and standard deviations are shown as error bars. (**A**) Left: EH37E1198822 KO genotyping PCR result (WT band: 3375 bp, genome edited band: 850 bp). Right: relative expression level change of the target gene *NGEF* after enhancer KO. (**B**) Left: EH37E1000386 genotyping PCR result (WT band: 1938 bp, genome edited band: 809 bp). Right: relative expression level change of the target gene *RORB* after enhancer KO. (**C**) Left: EH37E0114246 genotyping PCR result (WT band: 1849 bp, genome edited band: 1205 bp). Right: relative expression level change of the target gene *PLEKHO1* after enhancer KO. (**D**) Left: EH37E0426064 genotyping PCR result (WT band: 3069 bp, genome edited band: 2203 bp). Right: relative expression level change of the target gene *TOM1L2* after enhancer KO. Abbreviations: Cells, +e = phNPCs cells without RNPs underwent the same electroporation served as control; NTC = PCR non template control; BR = biological replicate.

## Discussion

Moving from computational predictions to functional evidence of gene regulatory activity remains an important and challenging area of modern genomics. Because there is substantial evidence for tissue-specific gene regulation and evidence that genetic risk for brain disorders resides within brain-enriched regulatory regions, maps of regulatory elements in brain-relevant cell types are of substantial value. In this study, we utilized a large-scale capture STARR-seq approach to validate putative enhancer regions and characterize the noncoding genomic landscape of phNPCs. We selected over 78,000 candidate enhancer regions based on data from the PFC, NPCs, and GWAS and identified 8,148 regions with enhancer activity in phNPCs. Of these regions, 6,612 were replicated in two separate experiments, demonstrating strong evidence of enhancer activity.

This study utilized the STARR-seq approach in a primary human neuronal cell line, suggesting this method may be applicable to additional human neuronal model systems including hiPSC- and hESC-derived neurons and neuronal organoids. The phNPC line is a particularly valuable model, because it very closely recapitulates the expression patterns and network architecture of the developing human fetal brain. Stein et al. (2014) found that the phNPC line had higher transcriptomic correlation with the *in vivo* human developing brain than the SH-SY5Y neuroblastoma cell line, hiPSC-derived NPCs, and hESC-derived NPCs (*16*). Our percentage of enriched regions (∼8.4%) was much higher than that seen in whole-genome STARR-seq experiments (∼0.06%; (*21*)) and similar to that seen in previous CapSTARR-seq studies in non-neuronal cell lines (*22*). Vanhille et al. (*22*) were the first to describe the CapSTARR-seq approach, and they used it to interrogate predicted regulatory regions in a mouse T-cell line. They tested over 7,000 regions and found ∼6% of their regions were classified as strong enhancers, a percentage very similar to what we observed in the phNPCs. Regions that were not identified as active enhancers in our CapSTARR-seq assay may represent other types of regulatory elements, including promoters, silencers, or insulators. As enhancers are highly dependent on cell type and developmental timepoint, these regions also may be active enhancer regions in a different cell type of the PFC or at a different stage of development.

We showed active enhancer regions were enriched for TFBS (**Table 2**), which is an established feature of enhancer genomic sequences (*42, 43*). Active enhancer regions from Panel 1 were enriched for binding sites for four different TF motifs, while Panel 2 was enriched for binding sites for nine different TF motifs. We hypothesize that this discrepancy is due to differences in size and candidate selection methods between the two panels. Panel 1 (22,400 regions) was substantially smaller than Panel 2 (56,215 regions). Additionally, Panel 1 was selected based on data from the PFC, while Panel 2 utilized data from PFC, NPCs, and the developing human brain. By comparing the TFBS present in our active enhancer regions with microarray data from the phNPC line (*15*), we found that the TFs that are enriched for binding sites within our active enhancer regions are also more highly expressed in the phNPCs than randomly selected TFs (**Figure 2**). This suggests that these specific TFs may play an important role in gene regulation within the phNPCs, specifically at the active enhancer regions identified through CapSTARR-seq. These data thus provide a resource for future interrogation of these TFs for their role in neurogenesis and development.

To further support the potential role of these TFs in neuronal gene regulation, several of the most enriched TFs have previous implications in brain development and disease. Expression of *JunB*, which was implicated in both panels of our CapSTARR-seq experiment, has been associated with cocaine use and addiction (*26*). SOX10 is involved in neural crest peripheral glial development (*24, 25*), and differential methylation of *SOX10* has been identified in brain tissue from individuals with schizophrenia (*27*). Polymorphisms in *SREBF2* also have been associated with schizophrenia (*28*). Other implicated TFs, including TP53 and YY1, are better known for their roles in cancer, but they have been implicated in neurodevelopment and psychiatric disease as well (*44–48*). The association of these TFs with neuronal development and disease provides further validation for the role of our active enhancer regions in important neuronal pathways.

To complete our understanding of enhancers, we need to link STARR-seq active regions with genes that these enhancers regulate. However, assigning enhancers to target genes remains a challenging task. To overcome this challenge, we integrated open chromatin data from ATAC-seq and expression profiles from RNA-seq to identify predicted target genes for our putative enhancers (*30*). We identified 2,288 unique predicted target genes that are regulated through 427 TFs. Following a pathway analysis, we found that our predicted target genes were heavily enriched for neuronally-associated pathways such as synapse organization, axon guidance, glutamatergic synapse, GABAergic synapse, and dopaminergic synapse. Disease-specific databases showed that these target genes were enriched for brain-related diseases, including Alzheimer’s disease and nervous system disease. We also saw high levels of overlap between our predicted enhancer regions and fetal eQTLs (28%; (*34*)), particularly when compared with adult eQTLs (5%; (*30*)), providing additional support for the phNPC line being a valuable proxy for the developing fetal human brain (*16*). A subset of our enhancer regions overlapped both adult and fetal eQTL regions, and the predicted target genes for this subset were enriched for GWAS variants associated with autism spectrum disorder and schizophrenia (Fig. S1). The enrichment of these target genes for neuronal pathways further supports the functionality of our putative enhancer regions in regulation of human brain development. Additionally, the implication of these target genes in brain-related diseases provides evidence that these putative enhancers may be involved in disease pathogenesis in both the fetal brain and the adult brain.

We also utilized eQTL data to assess whether eQTL variants affect the activity of our putative enhancers. We investigated 47 eQTL variants through MutSTARR-seq. We identified one variant (2.1%) that significantly affected enhancer activity, leading to an increase in activity observed through the STARR-seq approach. This proportion is in line with other studies investigating the effects of single nucleotide variants on regulatory activity through massively parallel reporter assays (*41, 49, 50*) and with large-scale eQTL analyses from the Genotype-Tissue Expression (GTEx) project demonstrating that approximately 1% of single nucleotide variants significantly affects gene expression (*51*). The enhancer containing our significant variant is located on chromosome 17 and is predicted to regulate 12 different genes: *AC126544.2*, *ARL17A*, *ARL17B*, *CR936218.1*, *CRHR1*, *FAM215B*, *KANSL1-AS1, KANSL1, LRRC37A, LRRC37A2, MAPT-AS1,* and *MAPT.* This region of chromosome 17 falls within the 17q21.31 region, which is a 1.5Mb inversion region (*35*). This region displays extremely high LD, explaining why one single eQTL is predicted to regulate expression of 12 different genes. Two major haplotypes, H1 and H2, exist in this region, with H1 being the more prevalent haplotype in individuals of European ancestry (∼80%). The H1 haplotype has been associated with a number of neurodegenerative diseases, including Alzheimer’s disease (*36*), Parkinson’s disease (*35, 37, 38*), and PSP (*40, 41*).

Our implicated enhancer in the 17q21.31 region showed increased enhancer activity with the eQTL variant (alternate allele), so it was important to determine if these target genes also showed increased expression in individuals with the alternate allele. Only two genes, *ARL17A* and *MAPT-AS1*, had decreased expression in the alternate allele case. The remaining target genes had increased expression, suggesting that increased enhancer activity correlates with increased expression. Interestingly, expression of a number of these target genes has been previously associated with psychiatric disease. *CRHR1* encodes a corticotropin-releasing hormone (CRH) receptor, a protein involved in hypothalamic-pituitary-adrenal axis-mediated response to stress (*52*). Increased expression of *CRHR1* has been implicated in anxiety and major depressive disorder, and CRHR1 antagonists have exhibited antidepressant effects (*53*). The *MAPT* gene encodes the tau protein, which is associated with neurodegenerative disorders including Alzheimer’s disease and, most prominently, frontotemporal dementia (*39*) and PSP (*40, 41*). Increased expression of *MAPT* has been observed in individuals with Alzheimer’s disease (*54*). One of the genes that showed decreased expression in the eQTL analysis, *MAPT-AS1*, encodes an anti-sense transcript of *MAPT* that negatively regulates translation of the tau protein (*55*). Decreased expression of *MAPT-AS1* has also been associated with neurodegenerative disorders (*55*). The correlation of our MutSTARR-seq enhancer findings and the eQTL expression data further supports the validity of our findings. Further, the extensive implication of these target genes with psychiatric and neurodegenerative disorders provides insight into how variation within these enhancer regions may contribute to disease development and progression.

To further validate the functional activity of candidate enhancers, we employed a dual RNP-mediated CRISPR/Cas9 deletion strategy to knock out enhancer regions in phNPCs and measure the expression level change of the predicted target gene. Our DNA genotyping and Sanger sequencing validation confirmed that the pair of upstream and downstream RNPs can work simultaneously to induce double strand breaks precisely at the expected CRISPR target sequences. Repair by non-homologous end-joining results in enhancer deletions ranging in size from 644 bp to 2,525 bp (Table S15), with KO editing efficiency 20% to 45% (Table S14). We observed a substantial decrease in expression level of these four target genes: 0.45 for *NGEF*, 0.56 for *RORB*, 0.16 for *PLEKHO1* and 0.42 for *TOM1L2*. This result demonstrates that these active enhancers do up-regulate transcription of the target gene tested. This is consistent with previous results of high enrichment of these enhancers in the CapSTARR-seq experiment revealing strong enhancer activity, which may partially explain the mechanism of the target gene expression knockdown phenotype observed.

Notably, *NGEF*, acting as a neuronal guanine nucleotide exchange factor, is highly expressed in the brain specifically in the caudate nucleus and plays a role in axon guidance regulating ephrin-induced growth cone collapse and dendritic spine morphogenesis (*56, 57*). GWAS have shown association between *NGEF* and schizophrenia (*58*) or bipolar disorder (*58, 59*). *RORB*, a clock gene involved in neurogenesis, stress response, and modulation of circadian rhythms, has been found to have positive associations with the pediatric bipolar phenotype in the case-control sample (*60*). A GWAS study identified *PLEKHO1*, a gene that plays a role in the regulation of the actin cytoskeleton, as a significant bipolar disorder risk locus (*61*).

*TOM1L2*, a gene encoding a protein putatively involved in intracellular protein transport, showed evidence of being causal in Alzheimer’s disease brain tissue. *TOM1L2* has a proportionately higher level of connectivity with known Alzheimer’s disease genes than other genes in the 17p LD block, which indicated its role as a possible Alzheimer’s disease susceptibility gene (*62*). Recently, two studies using human single cell RNA seq and human brain protein quantitative trait locus (pQTL) data showed *TOM1L2* was enriched in astrocytes, and higher levels of brain *TOM1L2* were associated with greater risk of Alzheimer’s disease (*63, 64*). Collectively, these target genes are important candidates for further functional investigation in the search for the molecular basis of psychiatric disorders.

While our study identified thousands of active enhancer regions in phNPCs, this approach also has a few inherent limitations. STARR-seq, by design, is a plasmid-based, ectopic approach (*20*). This design prevents us from investigating the activity of putative enhancers in their endogenous genomic context. That recognized, we did select our candidate enhancer regions based on endogenous functional genomic data (e.g., ATAC-seq, ChIP-seq) from the human brain. We also note that another very similar plasmid-based reporter assay also shows very high correspondence to endogenous gene regulatory predictions and experimental validation (*41*), suggesting that there is relatively good correspondence between these out of context assays and native genomic loci. Another limitation of our approach is that our Panel 1 design utilized data exclusively from the PFC. While this data indicates open chromatin and predicted active enhancer regions in the adult brain, it does not directly represent active enhancer regions in NPCs. We addressed this limitation in our Panel 2 design, which incorporated data from PFC, NPCs, and the developing human brain. As a result, we increased our rate of active enhancers from ∼4% in Panel 1 to ∼10% in Panel 2. We also validated several of our active enhancers through MutSTARR-seq and CRISPR-based approaches, but we could not conduct these validation experiments on the same scale as our initial STARR-seq screen. Additionally, elucidating the function of enhancers through the one enhancer-one target gene pair strategy utilized in our CRISPR experiment is limited by the fact that one enhancer can in principle act on multiple genes, or one gene can be regulated by multiple enhancers. Further experimental validations should be undertaken to achieve a comprehensive matching of enhancers and putative target genes, such as single cell RNA-seq (scRNA-seq) to reveal the whole transcriptome change after candidate enhancer KO, the establishment of the candidate enhancer KO mouse model, or pooled CRISPR interference (*41*).

Future experiments should also aim to identify the specific TFs involved in gene regulation at these enhancers, potentially by utilizing our TFBS motif analysis to knock down specific TFs in the phNPCs and examine effects on gene expression. We are also currently interrogating additional variants through the MutSTARR-seq approach to improve our understanding of the effects of these psychiatric disease-associated variants on enhancer activity. Finally, while the phNPCs recapitulate many features of embryonic and fetal corticogenesis and development, they do not mature past the mid-fetal stage (*16*). The open chromatin landscape of the phNPCs (*65*) also differs substantially from even closely related model systems like hESC-derived NPCs (https://www.encodeproject.org/experiments/ENCSR278FVO/; Fig. S3, Table S16), emphasizing the importance of cell type in enhancer studies. Similar studies should be conducted in models that better recapitulate later stages of brain development and the postnatal brain, such as brain organoids (*66*).

In this study, we identified over 8,000 regions with enhancer activity in a primary human neuronal progenitor line. We demonstrated that these enhancer regions were enriched for binding sites of TFs with high levels of expression in the phNPCs. Further, about 30% of these regions overlap with fetal or adult brain eQTLs, which provides a high-confidence group of brain enhancers. We also identified over 2,200 predicted target genes for these enhancer regions and showed that these target genes are implicated in a number of neuronal pathways and brain-related diseases. Finally, we performed functional validation on a subset of these enhancer regions through MutSTARR-seq and CRISPR-based approaches, demonstrating that variation in these regions affects enhancer activity and deletion of these regions affects gene expression. This study provides a comprehensive dataset of active enhancer regions in phNPCs and provides insight into how these enhancer regions may be involved in brain development and function.

## Materials and Methods

### phNPC Cell Line Generation and Maintenance

The phNPC line was obtained from Dr. Daniel Geschwind’s lab at UCLA. The creation of this line is described in detail in Konopka et al. (*15*) and Stein et al. (*16*). Briefly, the phNPC line was generated using a neurosphere isolation method from human fetal brains at 15-18 weeks post-conception. The specific line used for this study was named “3C” and was derived from a female fetus of Mexican descent.

Following isolation, the cells were established into a monolayer cell culture and grown in 10cm dishes coated with 5µg/mL poly-ornithine and 5µg/mL fibronectin. The base media for culturing proliferating phNPCs included the following components: neurobasal A medium (Invitrogen), antibiotic-antimycotic (1X; Gibco), BIT 9500 serum substitute (10%; StemCell Technologies), GlutaMAX (1X; Invitrogen), and heparin (1µg/mL). The following growth factors were freshly added to the base media to create the final proliferation media: EGF (20pg/µL; PeproTech), FGF (20pg/µL; PeproTech), PDGF (20pg/µL; PeproTech), and LIF (2ng/mL; MilliporeSigma). While culturing, half of the media was replaced every other day until the cells were about ∼80% confluent.

Once the phNPCs were ∼80% confluent, they were passaged using 0.25% trypsin-EDTA (Gibco) to a concentration of approximately 1-1.5 million cells per 10cm dish. All experiments were done using cells at low passage number (passage < 20) to ensure cellular integrity.

### CapSTARR-seq Panel Design

#### Panel 1

The first panel for CapSTARR-seq was created from a subset of enhancers identified in Wang et al. (*67*). Candidate enhancer regions were chosen using a matched filter process as outlined in Sethi et al. (*68*). Briefly, PFC samples from the ENCODE (*69*), Roadmap Epigenomics (*70*), and PsychENCODE projects were analyzed in order to annotate a set of active enhancers in the brain. These analyses led to the identification of about 79,000 brain-specific active enhancers (*67*). From this set of 79,000 enhancers, regions of the Roadmap PFC enhancers that overlapped with PsychENCODE PFC enhancers were identified as a set of high-confidence PFC enhancers (18,212 regions). The high-confidence PFC enhancers were regions with strong ATAC-seq and DNase signals, as well as strong H3K27ac signals from both the Roadmap PFC and PsychENCODE PFC ChIP-seq experiments. This set of high-confidence enhancers was included as targets in the first capture panel. In addition to this set of high-confidence enhancers, 165 regions from the initial set of 79,000 brain-specific enhancers were included that overlapped bipolar or schizophrenia GWAS variants from the GWAS catalog (*71*). The final group of regions added to the first panel included a set of 4,427 predicted enhancers from Kozlenkov et al. (*72*). This resulted in a final panel size of 22,804 regions spanning about 14Mbp of the genome (**Figure 1A**).

#### Panel 2

To design the second panel of targets for STARR-seq, we combined data from various data resources as well as leveraged the deep learning model DECODE (*73*) to identify targets with a high likelihood of being an active enhancer in the brain. We made use of existing bulk PFC ATAC-seq data from the HumanFC and BrainGVEX cohorts of PsychENCODE (*74, 75*). These peaks were processed in a way similar to cCREs as described in Moore et al. (*76*), which resulted in a total of ∼350,000 candidate enhancers derived from PFC. Next, we used DECODE (*73*) to analyze data from the ENCODE NPC cell line, including various histone modifications (H3K27ac, H3K4me1, H3K4me3, H3K9ac) and DNase data, and identified a total of ∼89,000 high resolution candidate enhancers. We intersected the results from the ENCODE DECODE analysis and the PFC ATAC-seq analysis, resulting in ∼72,000 candidate enhancers shared by both datasets. Next, we further validated this group of candidate enhancers using three independent data sources. Developmental enhancers from Trevino et al. (*77*) and de la Torre-Ubieta et al. (*65*), which represent *in vivo* data in developing brain, confirmed around ∼90% overlap with the designated panel (Fig. S4). GWAS and SNPs in LD (r^2^ > 0.4) were also intersected against the panel design, showing a total of 460 unique GWAS, and ∼30,000 total linked SNPs intersecting. This high level of intersection suggests the candidate enhancers identified potentially cover regions of the chromatin relevant to regulation of genes or loci associated with brain traits and diseases, bolstering their selection for validation. Finally, eQTL (*10, 67*) and transcriptome-wide association study hits from fetal human brain (*10, 78*) were intersected with ∼48% of the candidate panel suggesting high functional significance of the panel targets. Of the resulting panel of ∼72,000 targets, 65,000 were new as compared to panel 1 and the overlapping 7,000 were removed. To compensate for this loss, we furthermore added around 3,600 additional targets as controls and 3,400 targets derived as top scoring cCREs from the Weng Lab at the University of Massachusetts Medical School (**Figure 1B**).

### Probe Design

Probes were designed to capture the target regions using HyperDesign software from Roche Sequencing Solutions (Pleasanton, CA). Our regions (human genome hg38) were uploaded into the software using the following settings: maximum close matches = 20, overhang = 30bp. The regions were consolidated, meaning any overlapping regions were collapsed into a single continuous candidate region. This consolidation resulted in a final panel size of 22,400 regions for Panel 1 and 56,215 regions for Panel 2. The software then designed KAPA Target Enrichment Probes covering the inputted regions. These probes are 120bp in length and, following hybridization with genomic DNA, can be captured through a bead-based capture method. For Panel 1, the software predicted 98.5% coverage of the candidate regions. For Panel 2, the software predicted 99.3% coverage of the candidate regions. Missing coverage was due to repetitive regions that are often present in noncoding regions of the genome. Following selection through HyperDesign, KAPA Target Enrichment Probes were ordered through Roche Diagnostics. The manufacturer probe design changed between Panels 1 and 2, which resulted in a slightly higher “off-target” rate in Panel 2 (Table S1).

### Input Library Generation

Human male genomic DNA obtained from Promega (Ref:G1471; Madison, WI) was used to generate the input library. Two lots of DNA were used: Lot #0000305466 (concentration = 173ng/µL) and Lot #0000461400 (concentration = 197ng/µL). DNA was sheared using a Covaris (Woburn, MA) LE220 ultrasonicator with the following settings: peak incident power = 450, duty factor = 5%, cycles per burst = 200, treatment time = 120 seconds. After shearing, DNA fragments were size selected (∼500bp) and isolated from an agarose gel with the Qiagen (Germantown, MD) MinElute Gel Extraction Kit. The NEBNext Ultra End Repair/dA-tailing Module (New England Biolabs; Ipswich, MA) was used for end repair and A-tailing of the isolated, size-selected fragments. Ligation of custom adaptors was performed using the NEBNext Ultra Ligation Module for DNA, and the ligation products were cleaned up using 0.8X AMPure XP beads (Beckman Coulter; Indianapolis, IN). The custom adaptor sequences can be found in Table S17. They were designed using Integrated DNA Technologies (IDT; Coralville, IA) and produced using ion-exchange high performance liquid chromatography (IE-HPLC) purification. The resulting fragments were amplified using ligation-mediated polymerase chain reaction (LM-PCR) with Q5 Hot Start High-Fidelity 2X Master Mix (NEB) to allow the addition of homology arms necessary for cloning. The LM-PCR primers can be found in Table S17 and the cycle conditions were as follows: 98°C for 30 seconds; 10 cycles: 98°C for 10 seconds, 65°C for 30 seconds, 72°C for 30 seconds; 72°C for 2 minutes; hold at 4°C.

The LM-PCR products were then hybridized to the KAPA Target Enrichment Probes following the KAPA HyperCap Workflow v3.0 (Roche Diagnostics). To adjust this protocol for our cloning purposes, the LM-PCR primers MPI_ORI_F/R (Table S17) were used in place of Universal Enhancing Oligos and the Post-Capture PCR Oligos. After hybridization, the captured genomic regions were cloned into the hSTARR-seq_ORI vector (Addgene #99296; (*79*)) following linearization of the vector through restriction enzyme digestion using AgeI-HF (NEB) and SalI-HF (NEB). Cloning was done using Gibson Assembly Master Mix (NEB) and the resulting products were cleaned up using SPRIselect beads (Beckman Coulter) and purified using a Slide-A-Lyzer MINI Dialysis Device (Thermo Scientific; Waltham, MA). The purified library was transformed into MegaX D10 electrocompetent cells (Invitrogen; Waltham, MA) which were cultured in 4L of LB broth with 1X ampicillin until the culture reached an optical density of about 1.0. The plasmid library was isolated using a Qiagen Plasmid Plus Giga kit. Following isolation, the library was concentrated with an Amicon Ultra-15 Centrifugal Filter Unit (MilliporeSigma) and purified with a Slide-A-Lyzer MINI Dialysis Device to produce the final input plasmid library. This library was then amplified using Illumina (San Diego, CA) sequencing primers (Table S17) and cleaned up using 1.8X AMPure XP beads (Beckman Coulter). The input library was sequenced on one lane of an Illumina MiSeq at the University of Chicago Genomics Facility using MiSeq Reagent Kit V3 and 75bp paired-end reads.

### Transfection of Library and Output Library Preparation

The input capture library was electroporated into the phNPC line using a BTX (Holliston, MA) AgilePulse MAX large volume transfection system. The passage number and cell counts used for each capture panel were as follows:

– Panel 1, replicate 1 = 69 million cells, passage 19
– Panel 1, replicate 2 = 58 million cells, passage 18
– Panel 2, replicate 1 = 92 million cells, passage 18
– Panel 2, replicate 2 = 103 million cells, passage 18

We transfected 10μg of the input plasmid library per million cells using BTXpress High Performance Electroporation Solution (100μL per 5 million cells). Two different sets of electroporation parameters were used. For the first replicate of Panel 1, we used the following parameters: first pulse = 200V amplitude, 1ms duration, 20ms interval and second pulse = 130V amplitude, 3ms duration, 10ms interval. We were able to modify the parameters to improve transfection efficiency and cell viability following the first replicate of Panel 1, and the new parameters were used for the remaining replicates and panels. The new parameters were as follows: first pulse = 220V amplitude, 1ms duration, 20ms interval and second pulse = 140V amplitude, 3ms duration, 10ms interval. Electroporation resulted in a transfection efficiency of ∼70% and a cell viability rate of ∼40-60% post-electroporation. Transfection efficiency was determined using a pmaxGFP plasmid (Lonza; Basel, Switzerland), as this plasmid is similar in size to the hSTARR-seq_ORI vector.

RNA was isolated from the phNPCs 24 hours after electroporation using the Qiagen RNeasy Mini Kit. The Dynabeads mRNA DIRECT Purification Kit (Invitrogen) was used to isolate mRNA from the total RNA. The mRNA was treated with TURBO DNase (Invitrogen), cleaned up using the Zymo (Irvine, CA) RNA Clean and Concentrator kit, converted into cDNA using the SuperScript III First-Strand Synthesis SuperMix kit (Invitrogen), and treated with RNase A/T1 Mix (Thermo Scientific). The final cDNA sample was then split into 16 separate reactions for final PCR amplification using 16 unique Illumina indexing primers and Q5 Hot Start High-Fidelity 2X Master Mix (NEB). After amplifications, the reactions were pooled into a single sample, cleaned up using 1.8X AMPure beads (Beckman Coulter), and quantified. This output library was sequenced on one lane of an Illumina MiSeq at the University of Chicago Genomics Facility. Sequencing was performed using MiSeq Reagent Kit V3 and 75bp paired-end reads.

### Enhancer Peak Calling

Sequenced CapSTARR-seq libraries were processed using STARRPeaker v1.2, which includes a new feature to restrict peak calling analysis to a supplied capture panel (*23*). Both input DNA and output RNA libraries were aligned to GRCh38 reference genome (https://www.encodeproject.org/files/GRCh38_no_alt_analysis_set_GCA_000001405.15/) using BWA-MEM v0.7.17 (*80*). For alignments within each sub-reaction, we removed duplicates and filtered for properly aligned paired-end reads. We merged the filtered alignments from 16 sub-reactions to create a single BAM output file for STARRPeaker peak calling analysis. Default parameters were used for STARRPeaker except for the step size used to bin genome. 500-bp window length with a 50-bp step size was used. Capture region was extended by 50bp in each direction before binning. In addition to genomic input, three covariate tracks were utilized, namely GC-content, mappability, and folding energy prediction, to model the null distribution. We removed ENCODE blacklist regions (ENCFF419RSJ) from the analysis. We identified putative enhancer regions for each capture panel and replicate.

### Transcription Factor Binding Site Analysis

In each panel, we intersected technical replicates and defined them as high-confident enhancers when there was at least 20bp overlap. We used HOMER (*81*) and MEME-Suite (*82*) to perform motif enrichment analysis in the putative enhancers for each panel separately. Only those motifs that were detected by both HOMER and MEME-Suite were considered as true signals and used for the downstream analysis.

We performed motif discovery in the 200bp region around the center of the enhancers. For HOMER, the masked version of the genomes was used. The control regions were defined from STARR-seq negative constructs, which are regions with no enhancer activity in our CapSTARR-seq assay. We used the default setting of HOMER (v4.11.1), which allows for zero or one occurrence per sequence. To match the enriched sequences to known motifs, several well-known motif databases were used by HOMER, including HOMER motif database and JASPAR database. The novel motif discovery in MEME-Suite was carried out by XSTREME (*83*). For XSTREME, we used synthetic sequences of the second Markov order as the control. We used the XSTREME web server with the default parameter settings of MEME-Suite web-server (v5.4.1). We allowed any number of occurrences per sequence. TOMTOM (*84*) embedded in MEME-Suite (v5.4.1) matched the motifs to the 1,956 known motifs in the JASPAR Core database (*85*). At the end, FIMO (*86*) in MEME-Suite (v5.4.1) scanned the enhancer regions and denoted the positions of the enriched sequences.

### Transcription Factor Expression in phNPCs

For motifs that were enriched in the putative enhancers, we examined the expression level of their corresponding TFs in day 0 (pre-differentiation) phNPCs (data accessible at NCBI GEO database (*15*) accession GSE28046). For each target motif, we first determined its Illumina ID in Illumina HumanRef-8 v3.0 expression beadchip data table (data accessible at NCBI GEO database accession GPL6883), and then computed the expression score (log2 transformed, quantile normalized expression levels) of the associated TF. The expression score was averaged over the four replicates. Because motif matching is a noisy process, we did not limit our analysis to the best match provided by HOMER. For a given enriched sequence, we considered all the similar motifs with a HOMER score of at least 0.85. We then averaged the expression scores of similar motifs. For the background dataset, we chose 100 randomly selected TFs from Lambert et al. (*29*). We used t-tests to calculate p-values comparing expression levels of associated TFs from Panels 1 and 2 with the background dataset of random TFs.

### Predicted Target Genes and Pathway Analysis

For each of our putative enhancer regions, predicted target gene(s) were identified by integrating open chromatin peaks from ATAC-seq and expression profiles from RNA-seq (*30*). Specifically, ArchR was used to predict the most likely gene target (*87*). After identifying the most likely chromatin peak-to-gene linkages, open chromatin peaks were intersected with putative enhancers to establish high confidence enhancer-gene linkages. Of the total 8,148 enhancers, we identified a total of 2,288 unique linked genes (Table S6). We utilized a number of publicly available pathway analysis tools to examine the list of predicted target genes associated with our enhancer regions: DAVID ((*31*); https://david.ncifcrf.gov/), Enrichr ((*32*); https://maayanlab.cloud/Enrichr/), and STRING database ((*33*); https://string-db.org/). For each database, we inputted our list of predicted target genes to identify biological pathways enriched within that gene list. As a comparison, we also generated 10 random subsets of 2,288 genes from our entire candidate region list (Table S7). We inputted these random gene lists into each database to determine background p-values to which our CapSTARR-seq gene set could be compared using a one-sample t-test.

### MutSTARR-seq

To further validate the enhancers, we identified a subset of eQTLs identified from around 150 individuals with genotype and snRNA-Seq data intersecting the putative enhancer regions (*30*). eQTLs were identified by using standard linear model approaches with consideration of various covariates (i.e., age, disorder, batch, etc.).

Ranking of enhancers to be tested by MutSTARR-seq were prioritized based on the intersecting eQTL’s statistical significance, effect size, and cell type ubiquity to maximize functional effect on the enhancer region. A total of 54 enhancers were selected to be mutated according to the alternate allele present in the eQTL. For each enhancer, we created eBlock (IDT) gene fragments with (alternate) and without (reference) the eQTL. Of our candidate regions, 7 did not pass complexity and quality control tests by IDT due to repetitive elements. Those regions were excluded from the candidate list, so we tested a total of 47 regions. An additional 15 regions had to be trimmed from either the 5’ or 3’ ends to eliminate repetitive regions to allow the sequence to pass quality control tests. For the trimmed regions, we ensured that the eQTL variant was not affected by this trimming. The remaining regions passed complexity and quality control tests during oligo design with IDT software. The final list of regions is in Table S11.

We constructed the input library for MutSTARR-seq by cloning these eBlock fragments into the linearized hSTARR-seq ORI vector using Gibson Assembly Master Mix (NEB). We then introduced the transformed plasmids into MegaX DH10 electrocompetent cells (Invitrogen). Plasmids were subsequently extracted using the Qiagen Plasmid Plus Giga Kit. Dialysis was performed using a Slide-A-Lyzer mini dialysis device (Thermo Fisher), and the plasmids were concentrated using an Amicon spin centrifugal filter unit (Millipore Sigma). A small portion of these plasmids was also prepared with 32 Illumina library indexes and sent for sequencing along with the output library.

To generate the output library, a total of 25 million phNPCs (passage 17 and 18) were transfected with our input library using the same transfection protocol used for CapSTARR-seq. We performed four output replicates in total. Following transfection, cells were incubated for 24 hours and then RNA was isolated using the Zymo RNA extraction kit. Isolation of mRNA was done using the Dynabeads mRNA Direct Purification Kit (Invitrogen). The resulting mRNA was treated with TURBO DNase (Invitrogen) and subsequently purified using the Zymo RNA Clean and Concentrator kit. Similar to the library construction for the CapSTARR-seq experiment, mRNA transcripts were converted into cDNA using the SuperScript III First-Strand Synthesis SuperMix kit (Invitrogen) and treated with RNase A/T1 Mix (Thermo Scientific). The cDNA was then split into 32 portions and amplified using individual Illumina sequencing index primers. Libraries were prepared for sequencing using the Illumina MiSeq Reagent Kit V3-600bp to generate 300bp paired-end reads. They were sequenced with a 25% PhiX spike-in on two lanes (2 replicates per lane) of an Illumina MiSeq at the University of Chicago Genomics Facility.

Enhancer peaks were called as described above. A Chi-squared test was used to test for difference in enhancer activity between the alternate and reference alleles. The expected value was calculated by dividing the alternate input read count by the reference input read count. The observed values for each replicate were calculated by dividing the alternate output read count by the reference output read count. The Chi-squared statistic was calculated using these expected and observed values, and this statistic was used to determine the p-value for each tested eQTL.

### Candidate Selection for CRISPR/Cas9 Knockout (KO)

To prioritize candidate enhancers for further functional validation, we overlapped enriched enhancer regions from Panel 1 of CapSTARR-seq with 165 disease-associated GWAS regions and identified 29 psychiatric disease associated active enhancers. The target gene of the enhancer was defined as the gene with the shortest distance from transcription start site (TSS) to the enhancer. Then, we overlapped these enhancers with the gene regulatory network from the PsychENCODE integrative paper (*30*). Four enhancers (EH37E1198822, EH37E1000386, EH37E0114246 and EH37E0426064) that have the same predicted target gene were selected for the further functional validation (Table S18).

### KO of Top Candidate Enhancers in phNPCs through Ribonucleoprotein (RNP) - mediated CRISPR/Cas9 Genome Editing *Guide RNA (gRNA) Design*

For each enhancer tested, a pair of upstream and downstream gRNAs were designed in a 300 bp window of the 5’ and 3’ flanking regions of the enhancer with the IDT gRNA design algorithm (IDT). The gRNAs with on-target score > 50 and off-target score > 50 were chosen for custom synthesis from IDT (Table S15).

### RNP Complex Preparation

For each crRNA XT (or gRNA), we first prepared the crRNA XT plus shortened universal transactivatingRNA oligonucleotide (tracrRNA, IDT) duplex by incubating them at 95°C for 5 minutes in a thermocycler at equimolar concentrations (100 µM enhancer specific upstream or downstream crRNA XT 0.72 µl + 100 µM tracrRNA 0.72 µl), then keeping the duplex at room temperature for 15 minutes. 1.02 µl S.P. HiFi Cas9 nuclease V3 (IDT) and 0.54 µl PBS were added to the duplex on ice and then incubated at room temperature for 20 minutes to form the RNP complex. For each enhancer KO electroporation reaction, we combined upstream RNP with downstream RNP at equimolar quantity on ice to form enhancer specific RNPs pair.

### Electroporation of RNPs into phNPCs

The phNPC cell line was maintained as described above. Only cells with low passage number (P15 - P17) were used for the electroporation experiments with biological replicates (BR) design as BR1 and BR2. For the CRISPR-Cas9 protocol, we used the 4D-Nucleofector system and Amaxa P3 primary Cell 4D-Nucleofector X Kit S from Lonza. The electroporation buffer used was P3 primary cell Nucleofector Solution with Supplement 1 (Lonza). 2.5 × 10 ^5^ phNPCs cells were used per electroporation reaction in one cuvette of the 16-well Nucleocuvette Stripe (Lonza). After trypsinization, cells were washed with PBS and then resuspended in 20 µl Nucleofector solution with 1µl 100 µM Cas9 Electroporation enhancer (IDT) and 5 µl enhancer specific RNPs pair. The mixture of cells and RNPs was transferred to each cuvette of the 16 well Nucleocuvette strip and underwent the electroporation in 4D-Nucleofector X Unit with the program CL-133. The cells resuspended in the nucleofector solution without any RNPs pair added were transferred to the cuvette and underwent the electroporation simultaneously to serve as control. After electroporation, prewarmed recovery full media was added to each cuvette immediately. The cells of each electroporation reaction were then seeded to two corresponding wells of two Poly-ornithine/fibronectin coated 24 well plates. 1.25 × 10 ^5^ cells were seeded per well. The cells were incubated in the incubator for 24 hours followed by DNA extraction and RNA isolation.

### Enhancer KO Genotyping PCR

DNA was extracted from phNPCs with the QuickExtract DNA Extraction Solution (Lucigen). DNA lysate was used as genotyping PCR template with Q5 Hot Start High-Fidelity 2X Master Mix (NEB) and enhancer specific genotyping primer pair spanning the upstream and downstream Cas9-guide RNA cleavage sites (Table S19). The DNA input amount, the annealing temperature and PCR cycle numbers need to be optimized for each enhancer. In general, ∼10 ng DNA amount was used in PCR with annealing temperature ranging from 66.9 to 68 ℃ and 25 to 31 PCR cycles.

The uncleaved control wild type (WT) band and edited band after enhancer KO were separated by agarose gel electrophoresis prepared with SYBR Safe DNA gel stain (Invitrogen) in 1 X Tris/Acetic Acid/EDTA buffer (ThermoScientific). Gel images were obtained using a ChemiDoc MP Imaging System (Bio-Rad). The genome editing KO efficiency (percentage) was calculated through densitometric analysis. The DNA band intensities were analyzed using Image Labs software (Bio-Rad) by plotting the band intensities for each lane. The edited bands were cut from the gel and purified with QIAquick Gel Extraction Kit (Qiagen) for the Sanger sequencing (Azenta).

### Target Gene Expression Assay

RNA extraction from phNPCs and reverse transcription (RT) were performed with Power SYBR Green Cells-to-C_T_ kit (Invitrogen) according to the manufacturer’s instructions. Each predesigned PrimeTime qPCR Probe Assay (IDT) (Table S20) for the target of interest was first tested to confirm an amplification efficiency between 88%∼110%.

The cultured cells were washed with PBS, mixed with 65 µl lysis solution supplemented with DNase I, and incubated at room temperature for 5 min. 6.5 µl Stop Solution was mixed into the lysate and incubated at room temperature for 2 min. The concentration of the RNA lysate was measured with NanoDrop.

For each 50 µl RT reaction, 1 µg RNA lysate was mixed with 25 µl 2 X SYBR RT buffer, 2.5 µl 20 X RT Enzyme Mix and nuclease-free water. The RT reaction was incubated at 37 ℃ for 60 min, then at 95 ℃ for 5 min in a thermal cycler. The cDNA was amplified by real time qPCR using PrimeTime Gene Expression Master Mix (IDT) and the PrimeTime qPCR Probe Assay for the target of interest. Briefly, 2 µl cDNA lysate was mixed with 5 µl Master Mix, 0.5 µl (500 nM primers and 250 nM probes) PrimeTime Probe Assay, and 2.5 µl nuclease-free water. The primers and probe of reference gene beta-actin (ACTB) were used to normalize the cDNA loading for each reaction. For each sample, the 10 µl reaction was triplicated in 3 wells in a MicroAmp Optical 384-Well Reaction Plate (Applied Biosystems). Negative controls (no RNA or cDNA) were included to verify the absence of contamination. qPCR was performed using the QuantStudio 7 Flex Real-Time PCR System (Applied Biosystems). Amplification was performed using a two stages procedure (hold stage at 95 ℃, 3 min, then 40 cycles PCR stage (95 ℃, 15 s, 60 ℃, 60 s with a single fluorescence measurement), and resultant quantification threshold cycles (C_T_) were calculated using the default settings in the QuantStudio Real Time PCR Software v1.3 (Applied Biosystems) (Tables S21-S24). Results were analyzed using the Pfaffl mathematical model (*88*), with the control cells undergoing the electroporation simultaneously without any RNPs serving as calibrator.

## Supporting information

Supplementary Materials (includes text, figures, tables)

## Acknowledgments

We would like to acknowledge the members of the PsychENCODE consortium for their contributions to this work. The full author list for the PsychENCODE consortium can be found in the Supplementary Materials.

## Author contributions

Conceptualization: SCG, LC, MG, DG, KPW

Methodology: SCG, LC, MS, JL, GW, MS, MF, JRM, MG, DL, KPW

Investigation: SCG, LC, MS, MS, MF, JRM

Formal Analysis: JL, GW, MW, AH, MG, ZC, YC, JZ, DL, MG

Visualization: SCG, LC, JL, GW, DL Supervision: JRM, MG, DG, KPW

Writing – original draft: SCG, LC, JL, GW, DL

Writing – review & editing: MS, JRM, DL, MG, DG, KPW

## Data and materials availability

The source data described in this manuscript are available via the PsychENCODE Knowledge Portal (https://psychencode.synapse.org/). The PsychENCODE Knowledge Portal is a platform for accessing data, analyses, and tools generated through grants funded by the National Institute of Mental Health (NIMH) PsychENCODE Consortium. Data is available for general research use according to the following requirements for data access and data attribution: (https://psychencode.synapse.org/DataAccess). For access to content described in this manuscript see: https://doi.org/10.7303/syn50900302.1.

## References

1. Substance Abuse and Mental Health Services Administration. (HHS Publication No. PEP21-07-01-003, NSDUH Series H-56, 2021).

2. K. R. Merikangas, J. P. He, M. Burstein, S. A. Swanson, S. Avenevoli, L. Cui, C. Benjet, K. Georgiades, J. Swendsen, Lifetime prevalence of mental disorders in U.S. adolescents: results from the National Comorbidity Survey Replication--Adolescent Supplement (NCS-A). J Am Acad Child Adolesc Psychiatry 49, 980–989 (2010).

3. N. Robinson, S. E. Bergen, Environmental Risk Factors for Schizophrenia and Bipolar Disorder and Their Relationship to Genetic Risk: Current Knowledge and Future Directions. Front Genet 12, 686666 (2021).

4. T. Horwitz, K. Lam, Y. Chen, Y. Xia, C. Liu, A decade in psychiatric GWAS research. Mol Psychiatry 24, 378–389 (2019).

5. A. Barešić, A. J. Nash, T. Dahoun, O. Howes, B. Lenhard, Understanding the genetics of neuropsychiatric disorders: the potential role of genomic regulatory blocks. Mol Psychiatry 25, 6–18 (2020).

6. X. Chang, Y. Liu, C. G. Hahn, R. E. Gur, P. M. A. Sleiman, H. Hakonarson, RNA-seq analysis of amygdala tissue reveals characteristic expression profiles in schizophrenia. Transl Psychiatry 7, e1203 (2017).

7. R. C. Ramaker, K. M. Bowling, B. N. Lasseigne, M. H. Hagenauer, A. A. Hardigan, N. S. Davis, J. Gertz, P. M. Cartagena, D. M. Walsh, M. P. Vawter, E. G. Jones, A. F. Schatzberg, J. D. Barchas, S. J. Watson, B. G. Bunney, H. Akil, W. E. Bunney, J. Z. Li, S. J. Cooper, R. M. Myers, Post-mortem molecular profiling of three psychiatric disorders. Genome Med 9, 72 (2017).

8. S. P. Pantazatos, Y. Y. Huang, G. B. Rosoklija, A. J. Dwork, V. Arango, J. J. Mann, Whole-transcriptome brain expression and exon-usage profiling in major depression and suicide: evidence for altered glial, endothelial and ATPase activity. Mol Psychiatry 22, 760–773 (2017).

9. G. E. Hoffman, J. Bendl, G. Voloudakis, K. S. Montgomery, L. Sloofman, Y. C. Wang, H. R. Shah, M. E. Hauberg, J. S. Johnson, K. Girdhar, L. Song, J. F. Fullard, R. Kramer, C. G. Hahn, R. Gur, S. Marenco, B. K. Lipska, D. A. Lewis, V. Haroutunian, S. Hemby, P. Sullivan, S. Akbarian, A. Chess, J. D. Buxbaum, G. E. Crawford, E. Domenici, B. Devlin, S. K. Sieberts, M. A. Peters, P. Roussos, CommonMind Consortium provides transcriptomic and epigenomic data for Schizophrenia and Bipolar Disorder. Sci Data 6, 180 (2019).

10. R. L. Walker, G. Ramaswami, C. Hartl, N. Mancuso, M. J. Gandal, L. de la Torre-Ubieta, B. Pasaniuc, J. L. Stein, D. H. Geschwind, Genetic Control of Expression and Splicing in Developing Human Brain Informs Disease Mechanisms. Cell 179, 750–771.e722 (2019).

11. S. K. Sieberts, T. M. Perumal, M. M. Carrasquillo, M. Allen, J. S. Reddy, G. E. Hoffman, K. K. Dang, J. Calley, P. J. Ebert, J. Eddy, X. Wang, A. K. Greenwood, S. Mostafavi, L. Omberg, M. A. Peters, B. A. Logsdon, P. L. De Jager, N. Ertekin-Taner, L. M. Mangravite, C. C. (CMC), The A.-A. Consortium,Large eQTL meta-analysis reveals differing patterns between cerebral cortical and cerebellar brain regions. Sci Data 7, 340 (2020).

12. D. M. Werling, S. Pochareddy, J. Choi, J. Y. An, B. Sheppard, M. Peng, Z. Li, C. Dastmalchi, G. Santpere, A. M. M. Sousa, A. T. N. Tebbenkamp, N. Kaur, F. O. Gulden, M. S. Breen, L. Liang, M. C. Gilson, X. Zhao, S. Dong, L. Klei, A. E. Cicek, J. D. Buxbaum, H. Adle-Biassette, J. L. Thomas, K. A. Aldinger, D. R. O’Day, I. A. Glass, N. A. Zaitlen, M. E. Talkowski, K. Roeder, M. W. State, B. Devlin, S. J. Sanders, N. Sestan, Whole-Genome and RNA Sequencing Reveal Variation and Transcriptomic Coordination in the Developing Human Prefrontal Cortex. Cell Rep 31, 107489 (2020).

13. P. P. Zandi, A. E. Jaffe, F. S. Goes, E. E. Burke, L. Collado-Torres, L. Huuki-Myers, A. Seyedian, Y. Lin, F. Seifuddin, M. Pirooznia, C. A. Ross, J. E. Kleinman, D. R. Weinberger, T. M. Hyde, Amygdala and anterior cingulate transcriptomes from individuals with bipolar disorder reveal downregulated neuroimmune and synaptic pathways. Nat Neurosci 25, 381–389 (2022).

14. S. Akbarian, C. Liu, J. A. Knowles, F. M. Vaccarino, P. J. Farnham, G. E. Crawford, A. E. Jaffe, D. Pinto, S. Dracheva, D. H. Geschwind, J. Mill, A. C. Nairn, A. Abyzov, S. Pochareddy, S. Prabhakar, S. Weissman, P. F. Sullivan, M. W. State, Z. Weng, M. A. Peters, K. P. White, M. B. Gerstein, A. Amiri, C. Armoskus, A. E. Ashley-Koch, T. Bae, A. Beckel-Mitchener, B. P. Berman, G. A. Coetzee, G. Coppola, N. Francoeur, M. Fromer, R. Gao, K. Grennan, J. Herstein, D. H. Kavanagh, N. A. Ivanov, Y. Jiang, R. R. Kitchen, A. Kozlenkov, M. Kundakovic, M. Li, Z. Li, S. Liu, L. M. Mangravite, E. Mattei, E. Markenscoff-Papadimitriou, F. C. Navarro, N. North, L. Omberg, D. Panchision, N. Parikshak, J. Poschmann, A. J. Price, M. Purcaro, T. E. Reddy, P. Roussos, S. Schreiner, S. Scuderi, R. Sebra, M. Shibata, A. W. Shieh, M. Skarica, W. Sun, V. Swarup, A. Thomas, J. Tsuji, H. van Bakel, D. Wang, Y. Wang, K. Wang, D. M. Werling, A. J. Willsey, H. Witt, H. Won, C. C. Wong, G. A. Wray, E. Y. Wu, X. Xu, L. Yao, G. Senthil, T. Lehner, P. Sklar, N. Sestan, P. Consortium, The PsychENCODE project. Nat Neurosci 18, 1707–1712 (2015).

15. G. Konopka, E. Wexler, E. Rosen, Z. Mukamel, G. E. Osborn, L. Chen, D. Lu, F. Gao, K. Gao, J. K. Lowe, D. H. Geschwind, Modeling the functional genomics of autism using human neurons. Mol Psychiatry 17, 202–214 (2012).

16. J. L. Stein, L. de la Torre-Ubieta, Y. Tian, N. N. Parikshak, I. A. Hernández, M. C. Marchetto, D. K. Baker, D. Lu, C. R. Hinman, J. K. Lowe, E. M. Wexler, A. R. Muotri, F. H. Gage, K. S. Kosik, D. H. Geschwind, A quantitative framework to evaluate modeling of cortical development by neural stem cells. Neuron 83, 69–86 (2014).

17. O. Corradin, P. C. Scacheri, Enhancer variants: evaluating functions in common disease. Genome Med 6, 85 (2014).

18. A. C. Edwards, F. Aliev, L. J. Bierut, K. K. Bucholz, H. Edenberg, V. Hesselbrock, J. Kramer, S. Kuperman, J. I. Nurnberger, M. A. Schuckit, B. Porjesz, D. M. Dick, Genome-wide association study of comorbid depressive syndrome and alcohol dependence. Psychiatr Genet 22, 31–41 (2012).

19. S. Davidson, M. Lear, L. Shanley, B. Hing, A. Baizan-Edge, A. Herwig, J. P. Quinn, G. Breen, P. McGuffin, A. Starkey, P. Barrett, A. MacKenzie, Differential activity by polymorphic variants of a remote enhancer that supports galanin expression in the hypothalamus and amygdala: implications for obesity, depression and alcoholism. Neuropsychopharmacology 36, 2211–2221 (2011).

20. F. Muerdter, Ł. Boryń, C. D. Arnold, STARR-seq - principles and applications. Genomics 106, 145–150 (2015).

21. Y. Liu, S. Yu, V. K. Dhiman, T. Brunetti, H. Eckart, K. P. White, Functional assessment of human enhancer activities using whole-genome STARR-sequencing. Genome Biol 18, 219 (2017).

22. L. Vanhille, A. Griffon, M. A. Maqbool, J. Zacarias-Cabeza, L. T. Dao, N. Fernandez, B. Ballester, J. C. Andrau, S. Spicuglia, High-throughput and quantitative assessment of enhancer activity in mammals by CapStarr-seq. Nat Commun 6, 6905 (2015).

23. D. Lee, M. Shi, J. Moran, M. Wall, J. Zhang, J. Liu, D. Fitzgerald, Y. Kyono, L. Ma, K. P. White, M. Gerstein, STARRPeaker: Uniform processing and accurate identification of STARR-seq active regions. bioRxiv, (2020).

24. S. Britsch, D. E. Goerich, D. Riethmacher, R. I. Peirano, M. Rossner, K. A. Nave, C. Birchmeier, M. Wegner, The transcription factor Sox10 is a key regulator of peripheral glial development. Genes Dev 15, 66–78 (2001).

25. X. Lai, J. Liu, Z. Zou, Y. Wang, X. Liu, W. Huang, Y. Ma, Q. Chen, F. Li, G. Wu, W. Li, W. Wang, Y. Yuan, B. Jiang, SOX10 ablation severely impairs the generation of postmigratory neural crest from human pluripotent stem cells. Cell Death Dis 12, 814 (2021).

26. S. B. Huggett, M. C. Stallings, Genetic Architecture and Molecular Neuropathology of Human Cocaine Addiction. J Neurosci 40, 5300–5313 (2020).

27. L. F. Wockner, E. P. Noble, B. R. Lawford, R. M. Young, C. P. Morris, V. L. Whitehall, J. Voisey, Genome-wide DNA methylation analysis of human brain tissue from schizophrenia patients. Transl Psychiatry 4, e339 (2014).

28. Schizophrenia Working Group of the Psychiatric Genomics Consortium, Biological insights from 108 schizophrenia-associated genetic loci. Nature 511, 421-427 (2014).

29. S. A. Lambert, A. Jolma, L. F. Campitelli, P. K. Das, Y. Yin, M. Albu, X. Chen, J. Taipale, T. R. Hughes, M. T. Weirauch, The Human Transcription Factors. Cell 175, 598–599 (2018).

30. PsychENCODE Consortium, Single-cell genomics & regulatory networks for 388 human brains. Under revision in Science, (2024). 10.7303/syn51111084.1.

31. B. T. Sherman, M. Hao, J. Qiu, X. Jiao, M. W. Baseler, H. C. Lane, T. Imamichi, W. Chang, DAVID: a web server for functional enrichment analysis and functional annotation of gene lists (2021 update). Nucleic Acids Res 50, W216-W221 (2022).

32. Z. Xie, A. Bailey, M. V. Kuleshov, D. J. B. Clarke, J. E. Evangelista, S. L. Jenkins, A. Lachmann, M. L. Wojciechowicz, E. Kropiwnicki, K. M. Jagodnik, M. Jeon, A. Ma’ayan, Gene Set Knowledge Discovery with Enrichr. Curr Protoc 1, e90 (2021).

33. D. Szklarczyk, R. Kirsch, M. Koutrouli, K. Nastou, F. Mehryary, R. Hachilif, A. L. Gable, T. Fang, N. T. Doncheva, S. Pyysalo, P. Bork, L. J. Jensen, C. von Mering, The STRING database in 2023: protein-protein association networks and functional enrichment analyses for any sequenced genome of interest. Nucleic Acids Res 51, D638–D646 (2023).

34. C. Wen, M. Margolis, R. Dai, P. Zhang, P. F. Przytycki, D. D. Vo, A. Bhattacharya, M. Kim, N. Matoba, E. Tsai, C. Hoh, C. Jiao, N. Aygun, R. L. Walker, C. Chatzinakos, D. Clarke, H. Pratt, M. A. Peters, M. Gerstein, N. P. Daskalakis, Z. Weng, A. E. Jaffe, J. E. Kleinman, T. M. Hyde, D. R. Weinberger, N. J. Bray, N. Sestan, D. H. Geschwind, K. Roeder, A. Gusev, B. Pasaniuc, J. L. Stein, M. I. Love, K. S. Pollard, C. Liu, M. J. Gandal, P. Consortium, Cross-ancestry, cell-type-informed atlas of gene, isoform, and splicing regulation in the developing human brain. medRxiv, (2023).

35. K. R. Bowles, D. A. Pugh, Y. Liu, T. Patel, A. E. Renton, S. Bandres-Ciga, Z. Gan-Or, P. Heutink, A. Siitonen, S. Bertelsen, J. D. Cherry, C. M. Karch, S. J. Frucht, B. H. Kopell, I. Peter, Y. J. Park, A. Charney, T. Raj, J. F. Crary, A. M. Goate, I. P. s. D. G. C. (IPDGC), 17q21.31 sub-haplotypes underlying H1-associated risk for Parkinson’s disease are associated with LRRC37A/2 expression in astrocytes. Mol Neurodegener 17, 48 (2022).

36. G. Jun, C. A. Ibrahim-Verbaas, M. Vronskaya, J. C. Lambert, J. Chung, A. C. Naj, B. W. Kunkle, L. S. Wang, J. C. Bis, C. Bellenguez, D. Harold, K. L. Lunetta, A. L. Destefano, B. Grenier-Boley, R. Sims, G. W. Beecham, A. V. Smith, V. Chouraki, K. L. Hamilton-Nelson, M. A. Ikram, N. Fievet, N. Denning, E. R. Martin, H. Schmidt, Y. Kamatani, M. L. Dunstan, O. Valladares, A. R. Laza, D. Zelenika, A. Ramirez, T. M. Foroud, S. H. Choi, A. Boland, T. Becker, W. A. Kukull, S. J. van der Lee, F. Pasquier, C. Cruchaga, D. Beekly, A. L. Fitzpatrick, O. Hanon, M. Gill, R. Barber, V. Gudnason, D. Campion, S. Love, D. A. Bennett, N. Amin, C. Berr, M. Tsolaki, J. D. Buxbaum, O. L. Lopez, V. Deramecourt, N. C. Fox, L. B. Cantwell, L. Tárraga, C. Dufouil, J. Hardy, P. K. Crane, G. Eiriksdottir, D. Hannequin, R. Clarke, D. Evans, T. H. Mosley, L. Letenneur, C. Brayne, W. Maier, P. De Jager, V. Emilsson, J. F. Dartigues, H. Hampel, M. I. Kamboh, R. F. de Bruijn, C. Tzourio, P. Pastor, E. B. Larson, J. I. Rotter, M. C. O’Donovan, T. J. Montine, M. A. Nalls, S. Mead, E. M. Reiman, P. V. Jonsson, C. Holmes, P. H. St George-Hyslop, M. Boada, P. Passmore, J. R. Wendland, R. Schmidt, K. Morgan, A. R. Winslow, J. F. Powell, M. Carasquillo, S. G. Younkin, J. Jakobsdóttir, J. S. Kauwe, K. C. Wilhelmsen, D. Rujescu, M. M. Nöthen, A. Hofman, L. Jones, J. L. Haines, B. M. Psaty, C. Van Broeckhoven, P. Holmans, L. J. Launer, R. Mayeux, M. Lathrop, A. M. Goate, V. Escott-Price, S. Seshadri, M. A. Pericak-Vance, P. Amouyel, J. Williams, C. M. van Duijn, G. D. Schellenberg, L. A. Farrer, I. Consortium, A novel Alzheimer disease locus located near the gene encoding tau protein. Mol Psychiatry 21, 108–117 (2016).

37. M. A. Nalls, N. Pankratz, C. M. Lill, C. B. Do, D. G. Hernandez, M. Saad, A. L. DeStefano, E. Kara, J. Bras, M. Sharma, C. Schulte, M. F. Keller, S. Arepalli, C. Letson, C. Edsall, H. Stefansson, X. Liu, H. Pliner, J. H. Lee, R. Cheng, M. A. Ikram, J. P. Ioannidis, G. M. Hadjigeorgiou, J. C. Bis, M. Martinez, J. S. Perlmutter, A. Goate, K. Marder, B. Fiske, M. Sutherland, G. Xiromerisiou, R. H. Myers, L. N. Clark, K. Stefansson, J. A. Hardy, P. Heutink, H. Chen, N. W. Wood, H. Houlden, H. Payami, A. Brice, W. K. Scott, T. Gasser, L. Bertram, N. Eriksson, T. Foroud, A. B. Singleton, I. P. s. D. G. C. (IPDGC), P. s. S. G. P. P. s. R. T. O. G. I. (PROGENI), 23andMe, GenePD, N. R. C. (NGRC), H. I. o. H. G. (HIHG), A. J. D. Investigator, C. f. H. a. A. R. i. G. E. (CHARGE), N. A. B. E. C. (NABEC), U. K. B. E. C. (UKBEC), G. P. s. D. Consortium, A. G. A. Group, Large-scale meta-analysis of genome-wide association data identifies six new risk loci for Parkinson’s disease. Nat Genet 46, 989–993 (2014).

38. M. A. Nalls, C. Blauwendraat, C. L. Vallerga, K. Heilbron, S. Bandres-Ciga, D. Chang, M. Tan, D. A. Kia, A. J. Noyce, A. Xue, J. Bras, E. Young, R. von Coelln, J. Simón-Sánchez, C. Schulte, M. Sharma, L. Krohn, L. Pihlstrøm, A. Siitonen, H. Iwaki, H. Leonard, F. Faghri, J. R. Gibbs, D. G. Hernandez, S. W. Scholz, J. A. Botia, M. Martinez, J. C. Corvol, S. Lesage, J. Jankovic, L. M. Shulman, M. Sutherland, P. Tienari, K. Majamaa, M. Toft, O. A. Andreassen, T. Bangale, A. Brice, J. Yang, Z. Gan-Or, T. Gasser, P. Heutink, J. M. Shulman, N. W. Wood, D. A. Hinds, J. A. Hardy, H. R. Morris, J. Gratten, P. M. Visscher, R. R. Graham, A. B. Singleton, a. R. Team, S. G. o. P. s. D. Consortium, I. P. s. D. G. Consortium, Identification of novel risk loci, causal insights, and heritable risk for Parkinson’s disease: a meta-analysis of genome-wide association studies. Lancet Neurol 18, 1091–1102 (2019).

39. L. M. Reus, B. Pasaniuc, D. Posthuma, T. Boltz, Y. A. L. Pijnenburg, R. A. Ophoff, I. F.-G. Consortium, Gene Expression Imputation Across Multiple Tissue Types Provides Insight Into the Genetic Architecture of Frontotemporal Dementia and Its Clinical Subtypes. Biol Psychiatry 89, 825–835 (2021).

40. G. U. Höglinger, N. M. Melhem, D. W. Dickson, P. M. Sleiman, L. S. Wang, L. Klei, R. Rademakers, R. de Silva, I. Litvan, D. E. Riley, J. C. van Swieten, P. Heutink, Z. K. Wszolek, R. J. Uitti, J. Vandrovcova, H. I. Hurtig, R. G. Gross, W. Maetzler, S. Goldwurm, E. Tolosa, B. Borroni, P. Pastor, L. B. Cantwell, M. R. Han, A. Dillman, M. P. van der Brug, J. R. Gibbs, M. R. Cookson, D. G. Hernandez, A. B. Singleton, M. J. Farrer, C. E. Yu, L. I. Golbe, T. Revesz, J. Hardy, A. J. Lees, B. Devlin, H. Hakonarson, U. Müller, G. D. Schellenberg, P. G. S. Group, Identification of common variants influencing risk of the tauopathy progressive supranuclear palsy. Nat Genet 43, 699–705 (2011).

41. Y. A. Cooper, N. Teyssier, N. M. Dräger, Q. Guo, J. E. Davis, S. M. Sattler, Z. Yang, A. Patel, S. Wu, S. Kosuri, G. Coppola, M. Kampmann, D. H. Geschwind, Functional regulatory variants implicate distinct transcriptional networks in dementia. Science 377, eabi8654 (2022).

42. F. Spitz, E. E. Furlong, Transcription factors: from enhancer binding to developmental control. Nat Rev Genet 13, 613–626 (2012).

43. A. Panigrahi, B. W. O’Malley, Mechanisms of enhancer action: the known and the unknown. Genome Biol 22, 108 (2021).

44. J. A. Beagan, M. T. Duong, K. R. Titus, L. Zhou, Z. Cao, J. Ma, C. V. Lachanski, D. R. Gillis, J. E. Phillips-Cremins, YY1 and CTCF orchestrate a 3D chromatin looping switch during early neural lineage commitment. Genome Res 27, 1139–1152 (2017).

45. J. Yang, X. Wu, J. Huang, Z. Chen, G. Huang, X. Guo, L. Zhu, L. Su, TP53 Polymorphism Contributes to the Susceptibility to Bipolar Disorder but Not to Schizophrenia in the Chinese Han Population. J Mol Neurosci 68, 679–687 (2019).

46. Y. Li, C. Ma, W. Li, Y. Yang, X. Li, J. Liu, J. Wang, S. Li, Y. Liu, K. Li, J. Li, D. Huang, R. Chen, L. Lv, M. Li, X. J. Luo, A missense variant in NDUFA6 confers schizophrenia risk by affecting YY1 binding and NAGA expression. Mol Psychiatry 26, 6896–6911 (2021).

47. E. G. Contreras, J. Sierralta, A. Glavic, p53 is required for brain growth but is dispensable for resistance to nutrient restriction during Drosophila larval development. PLoS One 13, e0194344 (2018).

48. H. Yamagata, S. Uchida, K. Matsuo, K. Harada, A. Kobayashi, M. Nakashima, M. Nakano, K. Otsuki, N. Abe-Higuchi, F. Higuchi, T. Watanuki, T. Matsubara, S. Miyata, M. Fukuda, M. Mikuni, Y. Watanabe, Identification of commonly altered genes between in major depressive disorder and a mouse model of depression. Sci Rep 7, 3044 (2017).

49. C. V. Weiss, L. Harshman, F. Inoue, H. B. Fraser, D. A. Petrov, N. Ahituv, D. Gokhman, The cis-regulatory effects of modern human-specific variants. Elife 10, (2021).

50. C. Deng, S. Whalen, M. Steyert, R. Ziffra, P. F. Przytycki, F. Inoue, D. A. Pereira, D. Capauto, S. Norton, F. M. Vaccarino, A. Pollen, T. J. Nowakowski, N. Ahituv, K. S. Pollard, Massively parallel characterization of psychiatric disorder-associated and cell-type-specific regulatory elements in the developing human cortex. bioRxiv, (2023).

51. G. T. Consortium, D. A. Laboratory, G. Coordinating Center-Analysis Working, G. Statistical Methods groups-Analysis Working, G. g. Enhancing, N. I. H. C. Fund, Nih/Nci, Nih/Nhgri, Nih/Nimh, Nih/Nida, N. Biospecimen Collection Source Site, R. Biospecimen Collection Source Site, V. Biospecimen Core Resource, B. Brain Bank Repository-University of Miami Brain Endowment, M. Leidos Biomedical-Project, E. Study, I. Genome Browser Data, E. B. I. Visualization, I. Genome Browser Data, U. o. C. S. C. Visualization-Ucsc Genomics Institute, a. Lead, D. A. Laboratory, C. Coordinating, N. I. H. p. management, c. Biospecimen, Pathology, Q. T. L. m. w. g. e, A. Battle, C. D. Brown, B. E. Engelhardt, S. B. Montgomery, Genetic effects on gene expression across human tissues. Nature 550, 204–213 (2017).

52. J. M. Reul, F. Holsboer, On the role of corticotropin-releasing hormone receptors in anxiety and depression. Dialogues Clin Neurosci 4, 31–46 (2002).

53. J. Sun, L. Qiu, H. Zhang, Z. Zhou, L. Ju, J. Yang, CRHR1 antagonist alleviates LPS-induced depression-like behaviour in mice. BMC Psychiatry 23, 17 (2023).

54. F. Zhou, D. Wang, The associations between the MAPT polymorphisms and Alzheimer’s disease risk: a meta-analysis. Oncotarget 8, 43506–43520 (2017).

55. R. Simone, F. Javad, W. Emmett, O. G. Wilkins, F. L. Almeida, N. Barahona-Torres, J. Zareba-Paslawska, M. Ehteramyan, P. Zuccotti, A. Modelska, K. Siva, G. S. Virdi, J. S. Mitchell, J. Harley, V. A. Kay, G. Hondhamuni, D. Trabzuni, M. Ryten, S. Wray, E. Preza, D. A. Kia, A. Pittman, R. Ferrari, C. Manzoni, A. Lees, J. A. Hardy, M. A. Denti, A. Quattrone, R. Patani, P. Svenningsson, T. T. Warner, V. Plagnol, J. Ule, R. de Silva, MIR-NATs repress MAPT translation and aid proteostasis in neurodegeneration. Nature 594, 117–123 (2021).

56. N. R. Rodrigues, A. M. Theodosiou, M. A. Nesbit, L. Campbell, A. T. Tandle, D. Saranath, K. E. Davies, Characterization of Ngef, a novel member of the Dbl family of genes expressed predominantly in the caudate nucleus. Genomics 65, 53–61 (2000).

57. S. M. Shamah, M. Z. Lin, J. L. Goldberg, S. Estrach, M. Sahin, L. Hu, M. Bazalakova, R. L. Neve, G. Corfas, A. Debant, M. E. Greenberg, EphA receptors regulate growth cone dynamics through the novel guanine nucleotide exchange factor ephexin. Cell 105, 233–244 (2001).

58. Y. Wu, H. Cao, A. Baranova, H. Huang, S. Li, L. Cai, S. Rao, M. Dai, M. Xie, Y. Dou, Q. Hao, L. Zhu, X. Zhang, Y. Yao, F. Zhang, M. Xu, Q. Wang, Multi-trait analysis for genome-wide association study of five psychiatric disorders. Transl Psychiatry 10, 209 (2020).

59. X. Yao, J. T. Glessner, J. Li, X. Qi, X. Hou, C. Zhu, X. Li, M. E. March, L. Yang, F. D. Mentch, H. S. Hain, X. Meng, Q. Xia, H. Hakonarson, Integrative analysis of genome-wide association studies identifies novel loci associated with neuropsychiatric disorders. Transl Psychiatry 11, 69 (2021).

60. C. L. McGrath, S. J. Glatt, P. Sklar, H. Le-Niculescu, R. Kuczenski, A. E. Doyle, J. Biederman, E. Mick, S. V. Faraone, A. B. Niculescu, M. T. Tsuang, Evidence for genetic association of RORB with bipolar disorder. BMC Psychiatry 9, 70 (2009).

61. E. A. Stahl, G. Breen, A. J. Forstner, A. McQuillin, S. Ripke, V. Trubetskoy, M. Mattheisen, Y. Wang, J. R. I. Coleman, H. A. Gaspar, C. A. de Leeuw, S. Steinberg, J. M. W. Pavlides, M. Trzaskowski, E. M. Byrne, T. H. Pers, P. A. Holmans, A. L. Richards, L. Abbott, E. Agerbo, H. Akil, D. Albani, N. Alliey-Rodriguez, T. D. Als, A. Anjorin, V. Antilla, S. Awasthi, J. A. Badner, M. Bækvad-Hansen, J. D. Barchas, N. Bass, M. Bauer, R. Belliveau, S. E. Bergen, C. B. Pedersen, E. Bøen, M. P. Boks, J. Boocock, M. Budde, W. Bunney, M. Burmeister, J. Bybjerg-Grauholm, W. Byerley, M. Casas, F. Cerrato, P. Cervantes, K. Chambert, A. W. Charney, D. Chen, C. Churchhouse, T. K. Clarke, W. Coryell, D. W. Craig, C. Cruceanu, D. Curtis, P. M. Czerski, A. M. Dale, S. de Jong, F. Degenhardt, J. Del-Favero, J. R. DePaulo, S. Djurovic, A. L. Dobbyn, A. Dumont, T. Elvsåshagen, V. Escott-Price, C. C. Fan, S. B. Fischer, M. Flickinger, T. M. Foroud, L. Forty, J. Frank, C. Fraser, N. B. Freimer, L. Frisén, K. Gade, D. Gage, J. Garnham, C. Giambartolomei, M. G. Pedersen, J. Goldstein, S. D. Gordon, K. Gordon-Smith, E. K. Green, M. J. Green, T. A. Greenwood, J. Grove, W. Guan, J. Guzman-Parra, M. L. Hamshere, M. Hautzinger, U. Heilbronner, S. Herms, M. Hipolito, P. Hoffmann, D. Holland, L. Huckins, S. Jamain, J. S. Johnson, A. Juréus, R. Kandaswamy, R. Karlsson, J. L. Kennedy, S. Kittel-Schneider, J. A. Knowles, M. Kogevinas, A. C. Koller, R. Kupka, C. Lavebratt, J. Lawrence, W. B. Lawson, M. Leber, P. H. Lee, S. E. Levy, J. Z. Li, C. Liu, S. Lucae, A. Maaser, D. J. MacIntyre, P. B. Mahon, W. Maier, L. Martinsson, S. McCarroll, P. McGuffin, M. G. McInnis, J. D. McKay, H. Medeiros, S. E. Medland, F. Meng, L. Milani, G. W. Montgomery, D. W. Morris, T. W. Mühleisen, N. Mullins, H. Nguyen, C. M. Nievergelt, A. N. Adolfsson, E. A. Nwulia, C. O’Donovan, L. M. O. Loohuis, A. P. S. Ori, L. Oruc, U. Ösby, R. H. Perlis, A. Perry, A. Pfennig, J. B. Potash, S. M. Purcell, E. J. Regeer, A. Reif, C. S. Reinbold, J. P. Rice, F. Rivas, M. Rivera, P. Roussos, D. M. Ruderfer, E. Ryu, C. Sánchez-Mora, A. F. Schatzberg, W. A. Scheftner, N. J. Schork, C. Shannon Weickert, T. Shehktman, P. D. Shilling, E. Sigurdsson, C. Slaney, O. B. Smeland, J. L. Sobell, C. Søholm Hansen, A. T. Spijker, D. St Clair, M. Steffens, J. S. Strauss, F. Streit, J. Strohmaier, S. Szelinger, R. C. Thompson, T. E. Thorgeirsson, J. Treutlein, H. Vedder, W. Wang, S. J. Watson, T. W. Weickert, S. H. Witt, S. Xi, W. Xu, A. H. Young, P. Zandi, P. Zhang, S. Zöllner, R. Adolfsson, I. Agartz, M. Alda, L. Backlund, B. T. Baune, F. Bellivier, W. H. Berrettini, J. M. Biernacka, D. H. R. Blackwood, M. Boehnke, A. D. Børglum, A. Corvin, N. Craddock, M. J. Daly, U. Dannlowski, T. Esko, B. Etain, M. Frye, J. M. Fullerton, E. S. Gershon, M. Gill, F. Goes, M. Grigoroiu-Serbanescu, J. Hauser, D. M. Hougaard, C. M. Hultman, I. Jones, L. A. Jones, R. S. Kahn, G. Kirov, M. Landén, M. Leboyer, C. M. Lewis, Q. S. Li, J. Lissowska, N. G. Martin, F. Mayoral, S. L. McElroy, A. M. McIntosh, F. J. McMahon, I. Melle, A. Metspalu, P. B. Mitchell, G. Morken, O. Mors, P. B. Mortensen, B. Müller-Myhsok, R. M. Myers, B. M. Neale, V. Nimgaonkar, M. Nordentoft, M. M. Nöthen, M. C. O’Donovan, K. J. Oedegaard, M. J. Owen, S. A. Paciga, C. Pato, M. T. Pato, D. Posthuma, J. A. Ramos-Quiroga, M. Ribasés, M. Rietschel, G. A. Rouleau, M. Schalling, P. R. Schofield, T. G. Schulze, A. Serretti, J. W. Smoller, H. Stefansson, K. Stefansson, E. Stordal, P. F. Sullivan, G. Turecki, A. E. Vaaler, E. Vieta, J. B. Vincent, T. Werge, J. I. Nurnberger, N. R. Wray, A. Di Florio, H. J. Edenberg, S. Cichon, R. A. Ophoff, L. J. Scott, O. A. Andreassen, J. Kelsoe, P. Sklar, e. Consortium, B. Consortium, B. D. W. G. o. t. P. G. Consortium, Genome-wide association study identifies 30 loci associated with bipolar disorder. Nat Genet 51, 793–803 (2019).

62. C. A. Reynolds, M. G. Hong, U. K. Eriksson, K. Blennow, F. Wiklund, B. Johansson, B. Malmberg, S. Berg, A. Alexeyenko, H. Grönberg, M. Gatz, N. L. Pedersen, J. A. Prince, Analysis of lipid pathway genes indicates association of sequence variation near SREBF1/TOM1L2/ATPAF2 with dementia risk. Hum Mol Genet 19, 2068–2078 (2010).

63. Y. N. Ou, Y. X. Yang, Y. T. Deng, C. Zhang, H. Hu, B. S. Wu, Y. Liu, Y. J. Wang, Y. Zhu, J. Suckling, L. Tan, J. T. Yu, Identification of novel drug targets for Alzheimer’s disease by integrating genetics and proteomes from brain and blood. Mol Psychiatry 26, 6065–6073 (2021).

64. Y. J. Ge, Y. N. Ou, Y. T. Deng, B. S. Wu, L. Yang, Y. R. Zhang, S. D. Chen, Y. Y. Huang, Q. Dong, L. Tan, J. T. Yu, I. F.-G. Consortium, Prioritization of Drug Targets for Neurodegenerative Diseases by Integrating Genetic and Proteomic Data From Brain and Blood. Biol Psychiatry 93, 770–779 (2023).

65. L. de la Torre-Ubieta, J. L. Stein, H. Won, C. K. Opland, D. Liang, D. Lu, D. H. Geschwind, The Dynamic Landscape of Open Chromatin during Human Cortical Neurogenesis. Cell 172, 289–304.e218 (2018).

66. A. Gordon, S. J. Yoon, S. S. Tran, C. D. Makinson, J. Y. Park, J. Andersen, A. M. Valencia, S. Horvath, X. Xiao, J. R. Huguenard, S. P. Pașca, D. H. Geschwind, Long-term maturation of human cortical organoids matches key early postnatal transitions. Nat Neurosci 24, 331–342 (2021).

67. D. Wang, S. Liu, J. Warrell, H. Won, X. Shi, F. C. P. Navarro, D. Clarke, M. Gu, P. Emani, Y. T. Yang, M. Xu, M. J. Gandal, S. Lou, J. Zhang, J. J. Park, C. Yan, S. K. Rhie, K. Manakongtreecheep, H. Zhou, A. Nathan, M. Peters, E. Mattei, D. Fitzgerald, T. Brunetti, J. Moore, Y. Jiang, K. Girdhar, G. E. Hoffman, S. Kalayci, Z. H. Gümüş, G. E. Crawford, P. Roussos, S. Akbarian, A. E. Jaffe, K. P. White, Z. Weng, N. Sestan, D. H. Geschwind, J. A. Knowles, M. B. Gerstein, P. Consortium, Comprehensive functional genomic resource and integrative model for the human brain. Science 362, (2018).

68. A. Sethi, M. Gu, E. Gumusgoz, L. Chan, K. K. Yan, J. Rozowsky, I. Barozzi, V. Afzal, J. A. Akiyama, I. Plajzer-Frick, C. Yan, C. S. Novak, M. Kato, T. H. Garvin, Q. Pham, A. Harrington, B. J. Mannion, E. A. Lee, Y. Fukuda-Yuzawa, A. Visel, D. E. Dickel, K. Y. Yip, R. Sutton, L. A. Pennacchio, M. Gerstein, Supervised enhancer prediction with epigenetic pattern recognition and targeted validation. Nat Methods 17, 807–814 (2020).

69. ENCODE Project Consortium, An integrated encyclopedia of DNA elements in the human genome. Nature 489, 57–74 (2012).

70. A. Kundaje, W. Meuleman, J. Ernst, M. Bilenky, A. Yen, A. Heravi-Moussavi, P. Kheradpour, Z. Zhang, J. Wang, M. J. Ziller, V. Amin, J. W. Whitaker, M. D. Schultz, L. D. Ward, A. Sarkar, G. Quon, R. S. Sandstrom, M. L. Eaton, Y. C. Wu, A. R. Pfenning, X. Wang, M. Claussnitzer, Y. Liu, C. Coarfa, R. A. Harris, N. Shoresh, C. B. Epstein, E. Gjoneska, D. Leung, W. Xie, R. D. Hawkins, R. Lister, C. Hong, P. Gascard, A. J. Mungall, R. Moore, E. Chuah, A. Tam, T. K. Canfield, R. S. Hansen, R. Kaul, P. J. Sabo, M. S. Bansal, A. Carles, J. R. Dixon, K. H. Farh, S. Feizi, R. Karlic, A. R. Kim, A. Kulkarni, D. Li, R. Lowdon, G. Elliott, T. R. Mercer, S. J. Neph, V. Onuchic, P. Polak, N. Rajagopal, P. Ray, R. C. Sallari, K. T. Siebenthall, N. A. Sinnott-Armstrong, M. Stevens, R. E. Thurman, J. Wu, B. Zhang, X. Zhou, A. E. Beaudet, L. A. Boyer, P. L. De Jager, P. J. Farnham, S. J. Fisher, D. Haussler, S. J. Jones, W. Li, M. A. Marra, M. T. McManus, S. Sunyaev, J. A. Thomson, T. D. Tlsty, L. H. Tsai, W. Wang, R. A. Waterland, M. Q. Zhang, L. H. Chadwick, B. E. Bernstein, J. F. Costello, J. R. Ecker, M. Hirst, A. Meissner, A. Milosavljevic, B. Ren, J. A. Stamatoyannopoulos, T. Wang, M. Kellis, R. E. Consortium, Integrative analysis of 111 reference human epigenomes. Nature 518, 317–330 (2015).

71. A. Buniello, J. A. L. MacArthur, M. Cerezo, L. W. Harris, J. Hayhurst, C. Malangone, A. McMahon, J. Morales, E. Mountjoy, E. Sollis, D. Suveges, O. Vrousgou, P. L. Whetzel, R. Amode, J. A. Guillen, H. S. Riat, S. J. Trevanion, P. Hall, H. Junkins, P. Flicek, T. Burdett, L. A. Hindorff, F. Cunningham, H. Parkinson, The NHGRI-EBI GWAS Catalog of published genome-wide association studies, targeted arrays and summary statistics 2019. Nucleic Acids Res 47, D1005–D1012 (2019).

72. A. Kozlenkov, M. W. Vermunt, P. Apontes, J. Li, K. Hao, C. C. Sherwood, P. R. Hof, J. J. Ely, M. Wegner, E. A. Mukamel, M. P. Creyghton, E. V. Koonin, S. Dracheva, Evolution of regulatory signatures in primate cortical neurons at cell-type resolution. Proc Natl Acad Sci U S A 117, 28422–28432 (2020).

73. Z. Chen, J. Zhang, J. Liu, Y. Dai, D. Lee, M. R. Min, M. Xu, M. Gerstein, DECODE: a Deep-learning framework for Condensing enhancers and refining boundaries with large-scale functional assays. Bioinformatics 37, i280–i288 (2021).

74. J. F. Fullard, M. E. Hauberg, J. Bendl, G. Egervari, M. D. Cirnaru, S. M. Reach, J. Motl, M. E. Ehrlich, Y. L. Hurd, P. Roussos, An atlas of chromatin accessibility in the adult human brain. Genome Res 28, 1243–1252 (2018).

75. J. Bryois, M. E. Garrett, L. Song, A. Safi, P. Giusti-Rodriguez, G. D. Johnson, A. W. Shieh, A. Buil, J. F. Fullard, P. Roussos, P. Sklar, S. Akbarian, V. Haroutunian, C. A. Stockmeier, G. A. Wray, K. P. White, C. Liu, T. E. Reddy, A. Ashley-Koch, P. F. Sullivan, G. E. Crawford, Evaluation of chromatin accessibility in prefrontal cortex of individuals with schizophrenia. Nat Commun 9, 3121 (2018).

76. J. E. Moore, M. J. Purcaro, H. E. Pratt, C. B. Epstein, N. Shoresh, J. Adrian, T. Kawli, C. A. Davis, A. Dobin, R. Kaul, J. Halow, E. L. Van Nostrand, P. Freese, D. U. Gorkin, Y. Shen, Y. He, M. Mackiewicz, F. Pauli-Behn, B. A. Williams, A. Mortazavi, C. A. Keller, X. O. Zhang, S. I. Elhajjajy, J. Huey, D. E. Dickel, V. Snetkova, X. Wei, X. Wang, J. C. Rivera-Mulia, J. Rozowsky, J. Zhang, S. B. Chhetri, A. Victorsen, K. P. White, A. Visel, G. W. Yeo, C. B. Burge, E. Lécuyer, D. M. Gilbert, J. Dekker, J. Rinn, E. M. Mendenhall, J. R. Ecker, M. Kellis, R. J. Klein, W. S. Noble, A. Kundaje, R. Guigó, P. J. Farnham, J. M. Cherry, R. M. Myers, B. Ren, B. R. Graveley, M. B. Gerstein, L. A. Pennacchio, M. P. Snyder, B. E. Bernstein, B. Wold, R. C. Hardison, T. R. Gingeras, J. A. Stamatoyannopoulos, Z. Weng, E. P. Consortium, Expanded encyclopaedias of DNA elements in the human and mouse genomes. Nature 583, 699–710 (2020).

77. A. E. Trevino, N. Sinnott-Armstrong, J. Andersen, S. J. Yoon, N. Huber, J. K. Pritchard, H. Y. Chang, W. J. Greenleaf, S. P. Pașca, Chromatin accessibility dynamics in a model of human forebrain development. Science 367, (2020).

78. M. J. Gandal, P. Zhang, E. Hadjimichael, R. L. Walker, C. Chen, S. Liu, H. Won, H. van Bakel, M. Varghese, Y. Wang, A. W. Shieh, J. Haney, S. Parhami, J. Belmont, M. Kim, P. Moran Losada, Z. Khan, J. Mleczko, Y. Xia, R. Dai, D. Wang, Y. T. Yang, M. Xu, K. Fish, P. R. Hof, J. Warrell, D. Fitzgerald, K. White, A. E. Jaffe, M. A. Peters, M. Gerstein, C. Liu, L. M. Iakoucheva, D. Pinto, D. H. Geschwind, P. Consortium, Transcriptome-wide isoform-level dysregulation in ASD, schizophrenia, and bipolar disorder. Science 362, (2018).

79. F. Muerdter, Ł. Boryń, A. R. Woodfin, C. Neumayr, M. Rath, M. A. Zabidi, M. Pagani, V. Haberle, T. Kazmar, R. R. Catarino, K. Schernhuber, C. D. Arnold, A. Stark, Resolving systematic errors in widely used enhancer activity assays in human cells. Nat Methods 15, 141–149 (2018).

80. H. Li, Aligning sequence reads, clone sequences and assembly contigs with BWA-MEM. arXiv 1303.3997v1, (2013).

81. S. Heinz, C. Benner, N. Spann, E. Bertolino, Y. C. Lin, P. Laslo, J. X. Cheng, C. Murre, H. Singh, C. K. Glass, Simple combinations of lineage-determining transcription factors prime cis-regulatory elements required for macrophage and B cell identities. Mol Cell 38, 576–589 (2010).

82. T. L. Bailey, J. Johnson, C. E. Grant, W. S. Noble, The MEME Suite. Nucleic Acids Res 43, W39–49 (2015).

83. C. Grant, T. Bailey, XSTREME: Comprehensive motif analysis of biological sequence datasets. bioRxiv, (2021).

84. S. Gupta, J. A. Stamatoyannopoulos, T. L. Bailey, W. S. Noble, Quantifying similarity between motifs. Genome Biol 8, R24 (2007).

85. J. A. Castro-Mondragon, R. Riudavets-Puig, I. Rauluseviciute, R. B. Lemma, L. Turchi, R. Blanc-Mathieu, J. Lucas, P. Boddie, A. Khan, N. Manosalva Pérez, O. Fornes, T. Y. Leung, A. Aguirre, F. Hammal, D. Schmelter, D. Baranasic, B. Ballester, A. Sandelin, B. Lenhard, K. Vandepoele, W. W. Wasserman, F. Parcy, A. Mathelier, JASPAR 2022: the 9th release of the open-access database of transcription factor binding profiles. Nucleic Acids Res 50, D165–D173 (2022).

86. C. E. Grant, T. L. Bailey, W. S. Noble, FIMO: scanning for occurrences of a given motif. Bioinformatics 27, 1017–1018 (2011).

87. J. M. Granja, M. R. Corces, S. E. Pierce, S. T. Bagdatli, H. Choudhry, H. Y. Chang, W. J. Greenleaf, ArchR is a scalable software package for integrative single-cell chromatin accessibility analysis. Nat Genet 53, 403–411 (2021).

88. M. W. Pfaffl, A new mathematical model for relative quantification in real-time RT-PCR. Nucleic Acids Res 29, e45 (2001).

