## Supplementary Materials (includes text, figures, tables) for "Validation of Enhancer Regions in Primary Human Neural Progenitor Cells using Capture STARR-seq"

### Supplementary Text

PsychENCODE Authorship List

Schahram Akbarian[1], Alexej Abyzov[2], Nadav Ahituv[3], Dhivya Arasappan[4], Jose Juan Almagro Armenteros[5], Brian J. Beliveau[6], Jaroslav Bendl[1], Sabina Berretta[7], Rahul A. Bharadwaj[8], Arjun Bhattacharya[9], Lucy Bicks[9], Kristen Brennand[10], Davide Capauto[10], Frances A. Champagne[4], Tanima Chatterjee[10], Chris Chatzinakos[7], Yuhang Chen[10], H. Isaac Chen[11], Yuyan Cheng[9], Lijun Cheng[12], Andrew Chess[1], Jo-fan Chien[13], Zhiyuan Chu[10], Declan Clarke[10], Ashley Clement[3], Leonardo Collado-Torres[8], Gregory M. Cooper[14], Gregory E. Crawford[15], Rujia Dai[16], Nikolaos P. Daskalakis[7], Jose Davila-Velderrain[17], Amy Deep-Soboslay[8], Chengyu Deng[3], Christopher P. DiPietro[7], Stella Dracheva[1], Shiron Drusinsky[18], Ziheng Duan[19], Duc Duong[21], Cagatay Dursun[10], Nicholas J. Eagles[8], Jonathan Edelstein[1], Prashant S. Emani[10], John F. Fullard[1], Kiki Galani[22], Timur Galeev[10], Michael J. Gandal[11], Sophia Gaynor[12], Mark Gerstein[10], Daniel H. Geschwind[9], Kiran Girdhar[1], Fernando S. Goes[23], William Greenleaf[5], Jennifer Grundman[9], Hanmin Guo[5], Qiuyu Guo[9], Chirag Gupta[24], Yoav Hadas[1], Joachim Hallmayer[5], Xikun Han[22], Vahram Haroutunian[1], Natalie Hawken[9], Chuan He[25], Ella Henry[10], Stephanie C. Hicks[37], Marcus Ho[5], Li-Lun Ho[22], Gabriel E. Hoffman[1], Yiling Huang[5], Louise A. Huuki-Myers[8], Ahyeon Hwang[19], Thomas M. Hyde[8], Artemis Iatrou[7], Fumitaka Inoue[3], Aarti Jajoo[7], Matthew Jensen[10], Lihua Jiang[5], Peng Jin[21], Ting Jin[23], Connor Jops[11], Alexandre Jourdon[10], Riki Kawaguchi[9], Manolis Kellis[21], Saniya Khullar[24], Joel E. Kleinman[8], Steven P. Kleopoulos[1], Alex Kozlenkov[1], Arnold Kriegstein[3], Anshul Kundaje[5], Soumya Kundu[5], Cheyu Lee, University California Irvine[19], Donghoon Lee[1], Junhao Li[13], Mingfeng Li[10], Xiao Lin[1], Shuang Liu[10], Jason Liu[10], Jianyin Liu[9], Chunyu Liu[16], Shuang Liu[24], Shaoke Lou[10], Jacob M. Loupe[14], Dan Lu[26], Shaojie Ma[10], Liang Ma[27], Michael Margolis[9], Jessica Mariani[10], Keri Martinowich[8], Kristen R. Maynard[8], Samantha Mazariegos[9], Ran Meng[10], Richard M. Myers[14], Courtney Micallef[1], Tatiana Mikhailova[16], Guo-li Ming[11], Shahin Mohammadi[28], Emma Monte[5], Kelsey S. Montgomery[26], Jill E. Moore[29], Jennifer R. Moran[12], Eran A. Mukamel[13], Angus C. Nairn[10], Charles B. Nemeroff[30], Pengyu Ni[10], Scott Norton[10], Tomasz Nowakowski[3], Larsson Omberg[26], Stephanie C. Page[8], Saejeong Park[10], Ashok Patowary[9], Reenal Pattni[5], Geo Pertea[8], Mette A. Peters[26], Nishigandha Phalke[29], Dalila Pinto[1], Milos Pjanic[1], Sirisha Pochareddy[10], Katherine S. Pollard[3,18,19], Alex Pollen[3], Henry Pratt[29], Pawel F. Przytycki[18], Carolin Purmann[5], Zhaohui S. Qin[21], Ping-Ping Qu[5], Diana Quintero[9], Towfique Raj[1], Ananya S. Rajagopalan[10], Sarah Reach[1] , Thomas Reimonn[29], Kerry J. Ressler[7], Deanna Ross[4], Panos Roussos[1], Joel Rozowsky[10], Misir Ruth[1], W. Brad Ruzicka[7], Stephan J. Sanders[3,31], Juliane M. Schneider[26], Soraya Scuderi[10], Robert Sebra[1], Nenad Sestan[10], Nicholas Seyfried[21], Zhiping Shao[1], Nicole Shedd[29], Annie W. Shieh[32], Joo Heon Shin[8], Mario Skarica[10], Clara Snijders[7], Hongjun Song[11], Matthew W. State[3], Jason Stein[33], Marilyn Steyert[3], Sivan Subburaju[7], Thomas Sudhof[5], Michael Snyder[5], Ran Tao[8], Karen Therrien[1], Li-Huei Tsai[22], Alexander E. Urban[5], Flora M. Vaccarino[10], Harm van Bakel[1], Daniel Vo[11], Georgios Voloudakis[1], Brie Wamsley[9], Tao Wang[5], Sidney H. Wang[32], Daifeng Wang[24], Yifan Wang[2], Jonathan Warrell[10], Yu Wei[16], Annika K. Weimer[5], Daniel R. Weinberger[8], Cindy Wen[9], Zhiping Weng[29], Sean Whalen[18], Kevin P. White[34], A. Jeremy Willsey[3],

Hyejung Won[33], Wing Wong[5], Hao Wu[21], Feinan Wu[10], Stefan Wuchty[35], Dennis Wylie[4], Siwei Xu[20], Chloe X. Yap[36], Biao Zeng[1], Pan Zhang[9], Chunling Zhang[16], Bin Zhang[1], Jing Zhang[20], Yanqiong Zhang[33], Xiao Zhou[10], Ryan Ziffra[3], Zane R. Zeier[35], Trisha M. Zintel[26]

Affiliations
[1]Icahn School of Medicine at Mount Sinai, New York, NY, USA. [2]Mayo Clinic Rochester, Rochester, MN, USA. [3]University of California, San Francisco, San Francisco, CA, USA. [4]The University of Texas at Austin, Austin, TX, USA. [5]Stanford University, Stanford, CA, USA. [6]University of Washington, Seattle, WA, USA. [7]McLean Hospital, Harvard Medical School, Belmont, MA, USA. [8]Lieber Institute for Brain Development, Baltimore, MD, USA. [9]University of California, Los Angeles, Los Angeles, CA, USA. [10]Yale University, New Haven, CT, USA. [11]University of Pennsylvania, Philadelphia, PA, USA. [12]Tempus Labs, Inc., Chicago, IL, USA. [13]University of California, San Diego, San Diego, CA, USA. [14]HudsonAlpha Institute for Biotechnology, Huntsville, AL, USA. [15]Duke University, Durham, NC, USA. [16]SUNY Upstate Medical University, Syracuse, NY, USA. [17]Human Technopole, Milan, Italy. [18]Gladstone Institutes, San Francisco, CA, USA. [19]Chan Zuckerburg Biohub San Francisco, San Francisco, CA, USA. [20]University of California, Irvine, Irvine, CA, USA. [21]Emory University, Atlanta, GA, USA. [22]Massachusetts Institute of Technology, Cambridge, MA, USA. [23]Johns Hopkins University, Baltimore, MD, USA. [24]University of Wisconsin-Madison, Madison, WI, USA. [25]The University of Chicago, Chicago, IL, USA. [26]Sage Bionetworks, Seattle, WA, USA. [27]The University of Texas Health Science Center at San Antonio, San Antonio, TX, USA. [28]Broad Institute of MIT and Harvard, Cambridge, MA, USA. [29]University of Massachusetts Chan Medical School, Worcester, MA, USA. [30]The University of Texas at Austin Dell Medical School, Austin, MA, USA. [31]University of Oxford, Oxford, England, UK. [32]The University of Texas Health Science Center at Houston, Houston, TX, USA. [33]University of North Carolina at Chapel Hill, Chapel Hill, USA. [34]National University of Singapore, Singapore, Singapore. [35]University of Miami, Miami, FL, USA. [36]University of Queensland, Queensland, NZ. [37]Johns Hopkins Bloomberg School of Public Health, Baltimore, MD, USA.

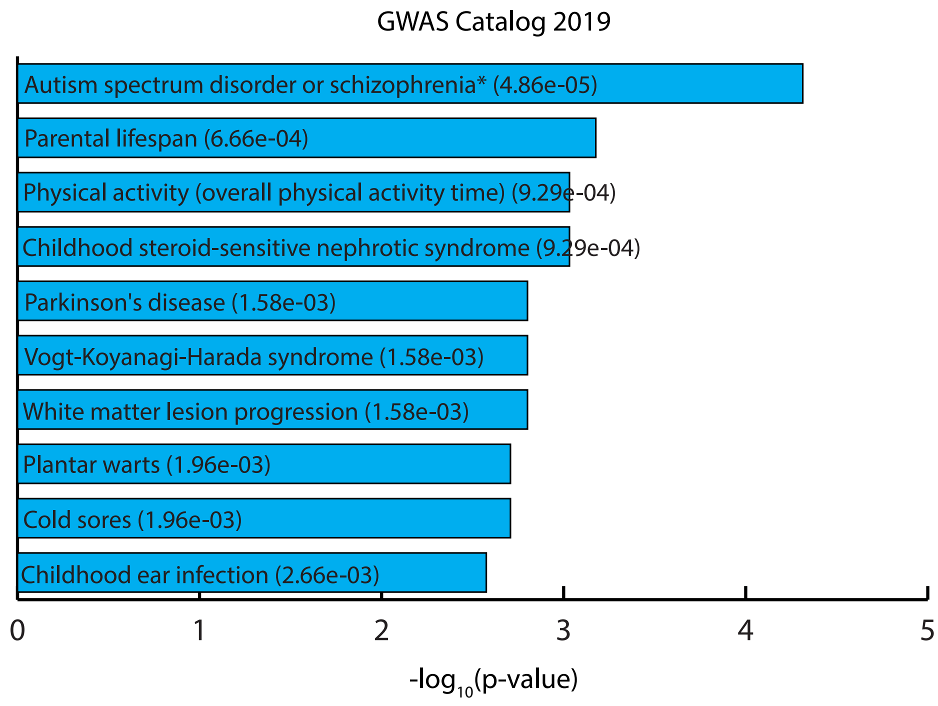

**Fig. S1.**
**Genome-wide association study (GWAS) pathway analysis results from Enrichr for target genes of enhancers overlapping adult and fetal expression quantitative trait loci (eQTLs).** The p-values for each category are included on the bars for each category. The asterisks (*) indicate that the adjusted p-value for that category is also significant (<0.05). Enrichr calculates p-values using the Fisher exact test and adjusted p-values using the Benjamini-Hochberg method.

A.

B.

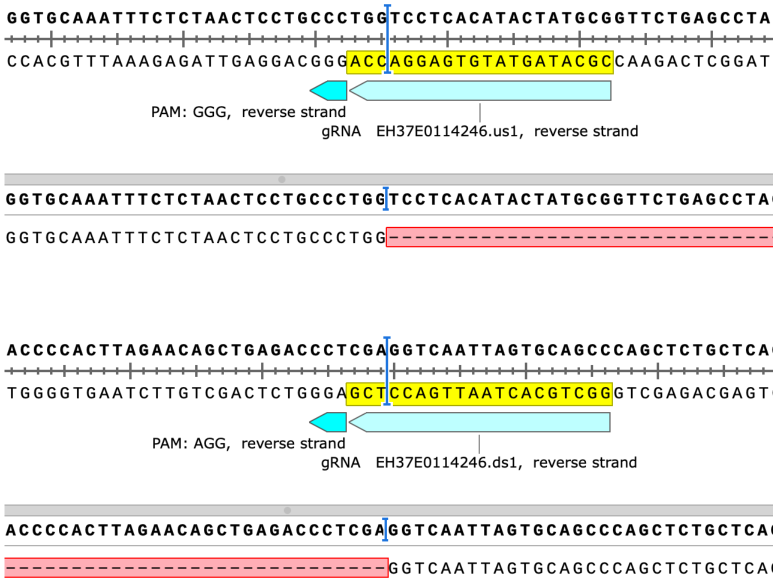

**Fig. S2.**

**Sanger sequencing results for CRISPR/Cas9 knockout (KO).** The Sanger sequencing results of the genome edited band after KO candidate enhancer showed that Cas9 nuclease always cuts 3 nucleotides precisely upstream of the PAM sites. Enhancer EH37E0114246 was used as an example here. The Sanger read was aligned to reference DNA sequence in SnapGene software. (**A**): Top sequence: reference DNA sequence. The upstream gRNA sequence was labeled in yellow color, followed by the PAM sequence GGG. Bottom sequence: The aligned Sanger read showed a cutting between "G" and "T" and just 3-nt upstream of the PAM site. (**B**): Top sequence: The downstream gRNA sequence was labeled in yellow color, followed by the PAM sequence AGG. Bottom sequence: The aligned Sanger read showed a cutting between "A" and "G" and 3-nt upstream of the PAM site. The dash line labeled with red color showed KO deletion between upstream and downstream cutting sites.

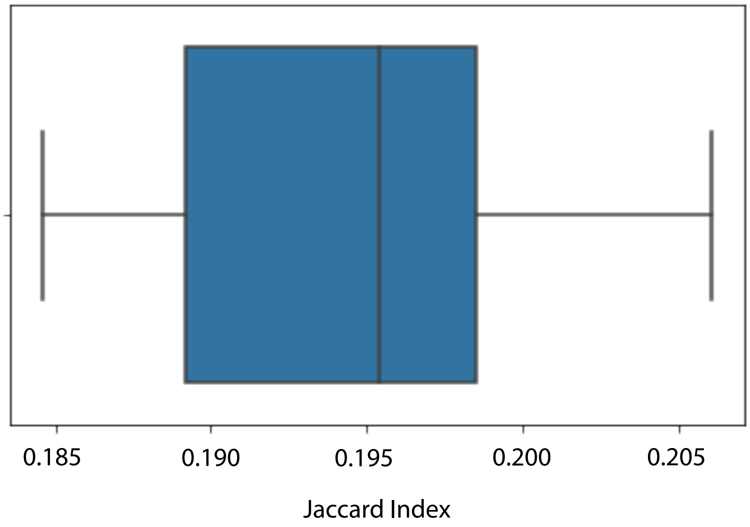

**Figure S3**.
**Comparison of open chromatin regions between primary human neural progenitor cells (phNPCs) and human embryonic stem cell (hESC)-derived NPCs**. Jaccard indices were calculated to determine the similarity between ATAC-seq data from phNPCs (*65*) and DNase-seq data from an hESC-derived NPC line (https://www.encodeproject.org/experiments/ENCSR278FVO/). ATAC-seq data was available for 10 different phNPC samples from the NPC-rich GZ zone. DNase-seq data was only available for a single hESC-derived NPC line. The individual Jaccard indices and the exact data files used are listed in Table S16.

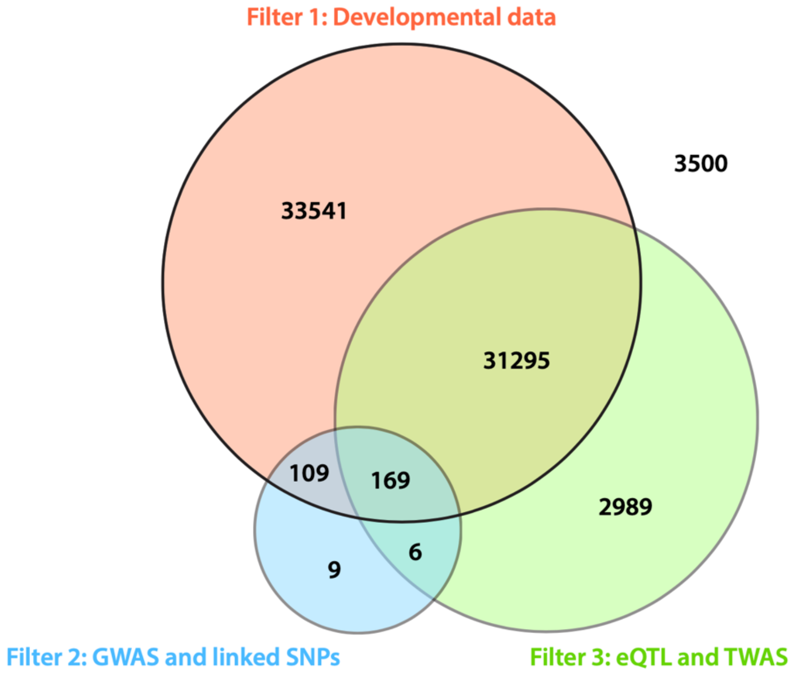

**Figure S4**.
**Intersection of candidate enhancers for Panel 2 and existing developmental, eQTL, TWAS and GWAS datasets**. We overlapped ENCODE data from neuronal progenitor cells and PsychENCODE ATAC-seq data from prefrontal cortex to identify an initial set of ~72,000 candidate enhancer regions for Panel 2 of CapSTARR-seq. This diagram represents the intersection between this set of candidates and three separate filters. Filter 1 (developmental) is shown is red. We examined overlap with *in vivo* enhancers from developing brain using data from (*77*) and (*65*). Filter 2 (GWAS and linked SNPs) is shown in blue. We identified 460 unique GWAS variants and ~30,000 total linked SNPs overlapping our candidate regions. Filter 3 (eQTL and TWAS) is shown in green. We examined overlap with eQTL and TWAS hits from fetal human brain using data from (*67*), (*78*), and (*10*). The remaining 3,500 regions (no color) represent putative enhancers from our initial list of ~72,000 candidate regions that did not overlap any of these datasets. Overall, we found that ~95% of our candidate regions overlapped at least one of these additional datasets. Abbreviations: eQTL = expression quantitative trait loci, TWAS = transcriptome-wide association study, GWAS = genome-wide association study, SNP = single nucleotide polymorphism.

| | Panel 1 | | | Panel 2 | | |
|---|---|---|---|---|---|---|
| | **Input** | **Output-R1** | **Output-R2** | **Input** | **Output-R1** | **Output-R2** |
| **Primary alignment** | 18,218,282 | 38,617,348 | 44,520,086 | 45,352,130 | 48,320,288 | 54,647,238 |
| **Properly paired** | 18,000,464 | 36,958,904 | 42,253,962 | 45,078,556 | 47,427,054 | 53,626,310 |
| | 98.80% | 95.71% | 94.91% | 99.40% | 98.15% | 98.13% |
| **PCR duplicates** | 2,155,985 | 10,037,373 | 13,877,561 | 16,571,564 | 22,608,412 | 26,941,994 |
| | 11.83% | 25.99% | 31.17% | 36.54% | 46.79% | 49.30% |
| **Barcoded duplicates** | - | 6,909,412 | 9,053,292 | - | 15,969,148 | 22,608,412 |
| **Library size** | 18,000,464 | 30,049,492 | 33,200,670 | 45,078,556 | 31,457,906 | 31,017,898 |
| **On-target** | 95.31% | 95.39% | 95.72% | 89.36% | 89.61% | 89.61% |
| **Off-target** | 4.69% | 4.61% | 4.28% | 10.64% | 10.39% | 10.39% |

**Table S1.**

**Quality control metrics for Panel 1 and Panel 2 input and output libraries**. Alignment and duplicate values represent read counts. Library size is measured in base pairs. On- and off-target percentages are a measure of the percentage of reads that fall within (on-target) or outside (off-target) our initial set of candidate regions for each panel. Abbreviations: R1 = replicate 1, R2 = replicate 2, PCR = polymerase chain reaction.

Too large – provided as a separate file.

**Table S2.**

**Active enhancer regions for each panel and each replicate as determined by STARRPeaker**. Panel replicates are on separate tabs of the spreadsheet. The “Name” column contains the peak rank based on score with 1 being the highest rank (most active enhancer). The “Score” column is an integer value calculated by multiplying the fold change by 100. The “Strand” column indicates on which strand of the DNA (+ or -) the enhancer is found. The “Log2 Fold Change” column is a normalized output to input ratio. The “Input” and “Output” fragment coverage values represent the total fragments across/within replicates.

| | Gene | Expression Score |
|---|---|---|
| **Motif 1** | JUNB | 9.73 |
| | FOSL2 | 8.77 |
| | ATF3 | 10.81 |
| | BATF | 7.41 |
| **Motif 1 Average** | | **9.18** |
| **Motif 2** | **TP53** | **10.40** |
| **Motif 3** | MITF | 8.30 |
| | TFE3 | 10.39 |
| | USF1 | 10.26 |
| | USF2(BHLH) | 9.04 |
| | ARNTL | 8.60 |
| | NPAS2(BHLH) | 10.09 |
| | TFEB | 7.34 |
| **Motif 3 Average** | | **9.15** |
| **Motif 4** | SOX10 | 7.48 |
| | SOX4 | 11.78 |
| | SOX11 | 13.51 |
| | SOX3 | 12.00 |
| | SOX2 | 13.25 |
| **Motif 4 Average** | | **11.60** |

**Table S3.**

**Expression scores for Panel 1 enriched transcription factors (TFs).** Expression scores were obtained from phNPC microarray data (*15*). The motifs represent the 4 enriched TF binding site motifs from Panel 1 CapSTARR-seq active enhancers. For some motifs, multiple TFs were included to account for similar TF binding site motifs from HOMER (see Materials and Methods; (*81*)). The values highlighted in red represent the motif values used for final expression comparisons.

| | Gene | Expression Score |
|---|---|---|
| **Motif 1** | **YY1** | **10.14** |
| Motif 2 | ELK1 | 12.22 |
| | ELK4 | 7.59 |
| | ETV4 | 10.45 |
| | GABPA | 6.98 |
| | ETV1 | 9.34 |
| | ELK3 | 8.93 |
| | ETV5 | 14.40 |
| | ETV4 | 10.45 |
| | ELF1 | 11.24 |
| **Motif 2 Average** | | **10.18** |
| **Motif 3** | **THAP11** | **13.00** |
| Motif 4 | SREBF2 | 8.88 |
| | SREBF1 | 7.86 |
| | BHLHE41 | 11.60 |
| | TFE3 | 10.39 |
| | MLX | 7.29 |
| **Motif 4 Average** | | **9.20** |
| **Motif 5** | **ZNF143** | **10.84** |
| **Motif 6** | **NRF1** | **8.21** |
| Motif 7 | FOSL2 | 8.77 |
| | FOSL1 | 11.43 |
| | JUND | 14.75 |
| | JUNB | 9.73 |
| **Motif 7 Average** | | **11.17** |
| **Motif 8** | **ZBTB33** | **12.77** |
| **Motif 9** | **TP53** | **10.40** |

**Table S4.**

**Expression scores for Panel 2 enriched TFs**. Expression scores were obtained from phNPC microarray data (*15*). The motifs represent the 9 enriched TF binding site motifs from Panel 2 CapSTARR-seq active enhancers. For some motifs, multiple TFs were included to account for similar TF binding site motifs from HOMER (see Materials and Methods; (*81*)). The values highlighted in red represent the motif values used for final expression comparisons.

| Gene | Expression Score |
|---|---|
| ATOH7 | 7.65 |
| ZNF649 | 10.79 |
| ZNF354A | 8.67 |
| ST18 | 7.38 |
| ZNF189 | 11.16 |
| ZNF80 | 7.53 |
| ANKZF1 | 9.98 |
| MEIS3 | 9.51 |
| LHX5 | 8.33 |
| ESRRB | 8.52 |
| PBX3 | 12.86 |
| NFIX | 13.75 |
| ZNF213 | 10.68 |
| ZSCAN1 | 8.08 |
| SCML4 | 7.08 |
| SOX30 | 7.76 |
| PITX1 | 7.64 |
| AHRR | 7.80 |
| PRDM10 | 10.24 |
| FOXD4 | 9.60 |
| NKX6-3 | 8.52 |
| TBX4 | 7.19 |
| ETV5 | 11.28 |
| FOXA2 | 7.12 |
| ZNF77 | 10.05 |
| FOXD2 | 7.69 |
| BCL6B | 8.62 |
| MESP1 | 9.73 |
| MYT1L | 7.28 |
| MAFG | 7.84 |
| ZNF74 | 9.20 |
| SOX17 | 7.07 |
| ZNF43 | 7.27 |
| UBP1 | 12.53 |
| ZNF587 | 7.75 |
| ZNF208 | 7.18 |
| ATF6 | 11.51 |

| | |
|---|---|
| TBX15 | 7.77 |
| TFAP2A | 7.70 |
| AHDC1 | 8.86 |
| ZNF18 | 10.57 |
| HES1 | 10.25 |
| SMYD3 | 12.18 |
| DMBX1 | 7.39 |
| ZNF670 | 8.97 |
| HOXD3 | 8.51 |
| ZNF254 | 9.90 |
| TFAP2E | 7.38 |
| MYCN | 10.02 |
| RELA | 10.51 |
| SP2 | 10.73 |
| HMG20B | 12.68 |
| CRX | 8.39 |
| ZNF7 | 10.55 |
| YY1 | 10.14 |
| SETBP1 | 11.20 |
| TFEC | 7.31 |
| ZBED5 | 12.04 |
| ZNF398 | 9.49 |
| ZNF341 | 9.59 |
| ZNF540 | 7.72 |
| EMX1 | 7.25 |
| ZNF19 | 7.96 |
| NFIL3 | 9.62 |
| CREBL2 | 8.16 |
| THAP2 | 7.07 |
| ZNF26 | 10.49 |
| BARHL1 | 7.46 |
| ZNF667 | 8.22 |
| ZNF423 | 12.15 |
| PPARD | 9.40 |
| MTF2 | 9.29 |
| ZNF557 | 9.79 |
| ZNF577 | 9.80 |
| ZFP2 | 9.71 |
| POU2AF1 | 7.84 |

| | |
|---|---|
| ZNF146 | 11.01 |
| ZNF366 | 8.24 |
| ZNF311 | 7.82 |
| HOXC8 | 7.35 |
| ZNF599 | 7.93 |
| TSC22D1 | 10.12 |
| PHF21A | 12.61 |
| SCMH1 | 9.06 |
| FOXR1 | 7.04 |
| NEUROG2 | 7.59 |
| BACH1 | 7.57 |
| PHF1 | 8.58 |
| ZNF41 | 7.87 |
| ELK4 | 7.59 |
| ZNF441 | 7.78 |
| ZNF615 | 10.84 |
| BATF2 | 7.90 |
| KLF17 | 7.90 |
| ZHX1 | 9.73 |
| ZNF214 | 7.18 |
| HOXB2 | 6.98 |
| ZNF8 | 8.27 |
| ZNF266 | 8.89 |
| AKAP8L | 10.69 |

**Table S5.**

**Expression scores for randomly selected TFs**. Expression scores were obtained from phNPC microarray data (*15*). These TFs were randomly selected from (*29*) to use as a background comparison set for the expression scores for the Panel 1 and Panel 2 enriched TFs.

Too large – provided as a separate file.

**Table S6.**

**Predicted target genes for active enhancer regions from CapSTARR-seq**. The chromosomal regions represent the location of the active enhancer regions. The “Transcription Factor” column indicates the transcription factor(s) predicted to bind those enhancer regions. The “Target Gene” column indicates the predicted target gene(s) for the enhancer regions.

Too large – provided as a separate file.

**Table S7.**

**Pathway analyses comparing CapSTARR-seq genes with background gene sets.** The "CapSTARRseq Genes" tab contains the list of 2,288 predicted target genes for the active enhancer regions identified in our CapSTARR-seq experiment. The "Entire Gene List" tab contains the list of putative target genes for our full candidate list of enhancer regions from Panels 1 and 2 regardless of STARR-seq activity. This list was used to generate the background sets of randomly selected genes found in the "Random Subsets" tab. We generated 10 random lists of 2,288 genes to use as a comparison for our CapSTARR-seq gene list. The "Comparison" tab contains the pathway analyses for these random subsets compared with our CapSTARR-seq gene list. We determined p-values for the random gene lists for each neuronal-associated pathway that was enriched in our CapSTARR-seq gene list. If a pathway was not identified for a given gene list, the p-value was designated as "n.s." for "not significant." The average, standard deviation, and standard error of the mean was calculated for the set of random gene lists, and these values were used to run a one-sample t-test comparing the p-value of the random gene lists to the CapSTARR-seq p-value. Pathways that were significantly more enriched in the CapSTARR-seq gene set are highlighted in yellow. Pathways containing an "N/A" entry were either not significant across any random gene lists or were significant in only a single gene list, making it impossible to perform a t-test for those pathways.

Too large – provided as a separate file.

**Table S8.**

**Enhancer regions overlapping adult and/or fetal expression quantitative trait loci (eQTLs)**. The enhancer regions that overlap eQTLs from “Adult” (*30*), “Fetal” (*34*), or “Both” are on separate tabs of the spreadsheet. Chromosomal locations for each enhancer, including starting and ending base pair, are indicated.

| Ensembl ID | Gene Name |
|---|---|
| ENSG00000165275 | TRMT10B |
| ENSG00000168803 | ADAL |
| ENSG00000106733 | NMRK1 |
| ENSG00000118197 | DDX59 |
| ENSG00000136235 | GPNMB |
| ENSG00000178878 | APOLD1 |
| ENSG00000120451 | SNX19 |
| ENSG00000089486 | CDIP1 |
| ENSG00000124587 | PEX6 |
| ENSG00000239335 | LLPH-AS1 |
| ENSG00000169752 | NRG4 |
| ENSG00000241316 | SUCLG2-AS1 |
| ENSG00000175643 | RMI2 |
| ENSG00000224043 | CCNT2-AS1 |
| ENSG00000257335 | MGAM |
| ENSG00000177112 | MRVI1-AS1 |
| ENSG00000262246 | CORO7 |
| ENSG00000234719 | NPIPB2 |
| ENSG00000197020 | ZNF100 |
| ENSG00000198502 | HLA-DRB5 |
| ENSG00000117335 | CD46 |
| ENSG00000237037 | NDUFA6-AS1 |
| ENSG00000157322 | CLEC18A |
| ENSG00000166435 | XRRA1 |
| ENSG00000254860 | TMEM9B-AS1 |
| ENSG00000162782 | TDRD5 |
| ENSG00000214401 | KANSL1-AS1 |
| ENSG00000180776 | ZDHHC20 |
| ENSG00000249846 | LINC02021 |
| ENSG00000183066 | WBP2NL |
| ENSG00000238083 | LRRC37A2 |
| ENSG00000233885 | YEATS2-AS1 |
| ENSG00000285053 | TBCE |
| ENSG00000213523 | SRA1 |
| ENSG00000154710 | RABGEF1 |
| ENSG00000151689 | INPP1 |
| ENSG00000228789 | HCG22 |

| | |
|---|---|
| ENSG00000112619 | PRPH2 |
| ENSG00000033327 | GAB2 |
| ENSG00000125388 | GRK4 |
| ENSG00000197062 | ZSCAN26 |
| ENSG00000118514 | ALDH8A1 |
| ENSG00000269293 | ZSCAN16-AS1 |
| ENSG00000136243 | NUPL2 |
| ENSG00000228775 | WEE2-AS1 |
| ENSG00000260088 | AL445483.1 |
| ENSG00000264589 | MAPT-AS1 |
| ENSG00000115084 | SLC35F5 |
| ENSG00000103550 | KNOP1 |
| ENSG00000185627 | PSMD13 |
| ENSG00000152219 | ARL14EP |
| ENSG00000115109 | EPB41L5 |
| ENSG00000204257 | HLA-DMA |
| ENSG00000228696 | ARL17B |
| ENSG00000185418 | TARSL2 |
| ENSG00000214087 | ARL16 |
| ENSG00000223745 | CCDC18-AS1 |
| ENSG00000135828 | RNASEL |
| ENSG00000280670 | CCDC163 |
| ENSG00000267871 | ZNF460-AS1 |
| ENSG00000152348 | ATG10 |
| ENSG00000180481 | GLIPR1L2 |
| ENSG00000134265 | NAPG |
| ENSG00000196118 | CCDC189 |
| ENSG00000136982 | DSCC1 |
| ENSG00000157578 | LCA5L |
| ENSG00000247572 | CKMT2-AS1 |
| ENSG00000188825 | LINC00910 |
| ENSG00000197124 | ZNF682 |
| ENSG00000161692 | DBF4B |
| ENSG00000225138 | SLC9A3-AS1 |
| ENSG00000176222 | ZNF404 |
| ENSG00000138760 | SCARB2 |
| ENSG00000272274 | LINC00551 |
| ENSG00000102858 | MGRN1 |

| | |
|---|---|
| ENSG00000225914 | HCG23 |
| ENSG00000103168 | TAF1C |
| ENSG00000263715 | LINC02210-CRHR1 |
| ENSG00000197013 | ZNF429 |
| ENSG00000204740 | MALRD1 |
| ENSG00000130653 | PNPLA7 |
| ENSG00000204138 | PHACTR4 |
| ENSG00000226686 | LINC01535 |
| ENSG00000176681 | LRRC37A |
| ENSG00000196126 | HLA-DRB1 |
| ENSG00000124713 | GNMT |
| ENSG00000066056 | TIE1 |
| ENSG00000205464 | ATP6AP1L |
| ENSG00000186994 | KANK3 |
| ENSG00000169684 | CHRNA5 |
| ENSG00000105248 | CCDC94 |
| ENSG00000204909 | SPINK9 |
| ENSG00000164048 | ZNF589 |
| ENSG00000256771 | ZNF253 |
| ENSG00000182093 | WRB |
| ENSG00000141576 | RNF157 |
| ENSG00000198185 | ZNF334 |
| ENSG00000178935 | ZNF552 |
| ENSG00000149781 | FERMT3 |
| ENSG00000196993 | NPIPB9 |
| ENSG00000232229 | LINC00865 |
| ENSG00000106261 | ZKSCAN1 |
| ENSG00000164167 | LSM6 |
| ENSG00000085982 | USP40 |
| ENSG00000149089 | APIP |
| ENSG00000076685 | NT5C2 |
| ENSG00000188312 | CENPP |
| ENSG00000123427 | EEF1AKMT3 |
| ENSG00000260230 | FRRS1L |
| ENSG00000104613 | INTS10 |
| ENSG00000075131 | TIPIN |
| ENSG00000174652 | ZNF266 |

| | |
|---|---|
| ENSG00000101751 | POLI |
| ENSG00000242574 | HLA-DMB |
| ENSG00000120071 | KANSL1 |
| ENSG00000054983 | GALC |
| ENSG00000162753 | SLC9C2 |
| ENSG00000223496 | EXOSC6 |
| ENSG00000100197 | CYP2D6 |
| ENSG00000227676 | LINC01068 |
| ENSG00000214435 | AS3MT |
| ENSG00000185324 | CDK10 |
| ENSG00000123415 | SMUG1 |
| ENSG00000231365 | AL359915.2 |
| ENSG00000163576 | EFHB |
| ENSG00000225231 | LINC02470 |
| ENSG00000026103 | FAS |
| ENSG00000260630 | SNAI3-AS1 |
| ENSG00000142675 | CNKSR1 |
| ENSG00000206503 | HLA-A |
| ENSG00000087365 | SF3B2 |
| ENSG00000141519 | CCDC40 |
| ENSG00000103479 | RBL2 |
| ENSG00000139624 | CERS5 |
| ENSG00000143891 | GALM |

**Table S9.**
**Predicted target genes for enhancers overlapping adult and fetal eQTLs.** Table includes the predicted target genes (Ensembl identifier and associated gene name) for enhancer regions that overlapped both an adult and fetal eQTL and had the same target gene with the same direction of effect.

| Category | Description | # Genes | FDR |
|---|---|---|---|
| KEGG_PATHWAY | Herpes simplex virus 1 infection | 15 | 8.20E-05 |
| GOTERM_BP_DIRECT | antigen processing and presentation of peptide or polysaccharide antigen via MHC class II | 5 | 1.20E-03 |
| KEGG_PATHWAY | Allograft rejection | 6 | 1.20E-04 |
| KEGG_PATHWAY | Graft-versus-host disease | 6 | 1.20E-04 |
| KEGG_PATHWAY | Type I diabetes mellitus | 6 | 1.20E-04 |
| KEGG_PATHWAY | Autoimmune thyroid disease | 6 | 2.80E-04 |
| GOTERM_BP_DIRECT | antigen processing and presentation | 5 | 9.00E-03 |
| GOTERM_BP_DIRECT | immunoglobulin production involved in immunoglobulin mediated immune response | 4 | 9.00E-03 |
| GOTERM_BP_DIRECT | peptide antigen assembly with MHC class II protein complex | 4 | 9.00E-03 |
| KEGG_PATHWAY | Viral myocarditis | 5 | 7.10E-03 |
| GOTERM_BP_DIRECT | antigen processing and presentation of exogenous peptide antigen via MHC class II | 4 | 5.00E-02 |
| GOTERM_BP_DIRECT | positive regulation of T cell activation | 4 | 5.00E-02 |
| KEGG_PATHWAY | Asthma | 4 | 1.20E-02 |
| KEGG_PATHWAY | Antigen processing and presentation | 5 | 1.30E-02 |
| KEGG_PATHWAY | Epstein-Barr virus infection | 7 | 1.30E-02 |
| KEGG_PATHWAY | Intestinal immune network for IgA production | 4 | 3.00E-02 |
| KEGG_PATHWAY | Influenza A | 6 | 3.00E-02 |

**Table S10.**

**DAVID analysis of predicted target genes for adult and fetal eQTL overlap**. Table includes all results for the GOTERM_BP_DIRECT, KEGG_PATHWAY, and GAD_DISEASE_CLASS categories with a false discovery rate (FDR) < 0.05. The "# Genes" column indicates the number of genes from the overlap set present in that specific pathway. DAVID calculates p-values using the Fisher exact test and false discovery rates (FDR) using the Benjamini-Hochberg method.

| Chromosome Location | eQTL Location | Reference Allele | Alternate Allele | eQTL P-value | Region trimmed? | Trimmed Region |
|---|---|---|---|---|---|---|
| chr1: 111755425-111756148 | chr1:111755544 | G | C | 1.83E-07 | N | - |
| chr1: 111755425-111756148 | chr1:111755960 | A | G | 5.00E-07 | N | - |
| chr1: 23959693-23960354 | chr1:23959818 | A | T | 5.94E-07 | N | - |
| chr1: 45687055-45687555 | chr1:45687531 | G | A | 3.31E-06 | N | - |
| chr11: 36289099-36289750 | chr11:36289709 | G | T | 4.03E-10 | N | - |
| chr11: 45056395-45057068 | chr11:45056899 | C | G | 2.23E-08 | N | - |
| chr11: 6234461-6234961 | chr11:6234708 | G | T | 3.78E-08 | Y | chr11: 6234471-6234951 |
| chr14: 55107549-55108173 | chr14:55108075 | T | C | 2.17E-06 | N | - |
| chr15: 39920700-39921303 | chr15:39920886 | G | T | 5.63E-09 | N | - |
| chr16: 3106107-3106660 | chr16:3106484 | A | C | 1.30E-08 | N | - |
| chr17: 20868246-20868746 | chr17:20868673 | G | A | 1.56E-06 | N | - |
| chr17: 43984025-43984644 | chr17:43984155 | TA | T | 8.40E-10 | N | - |

| | | | | | | |
|---|---|---|---|---|---|---|
| chr17: 45490502-45491036 | chr17:45490914 | G | A | 4.50E-07 | N | - |
| chr17: 45740659-45741399 | chr17:45740794 | A | C | 1.73E-06 | N | - |
| chr17: 45740659-45741399 | chr17:45740856 | C | A | 1.73E-06 | N | - |
| chr17: 45740659-45741399 | chr17:45741245 | G | A | 1.73E-06 | N | - |
| chr17: 45740659-45741399 | chr17:45741291 | T | C | 1.73E-06 | N | - |
| chr17: 45740659-45741399 | chr17:45741324 | G | T | 1.73E-06 | N | - |
| chr17: 45866949-45867697 | chr17:45867153 | C | T | 1.73E-06 | N | - |
| chr17: 45894107-45894607 | chr17:45894115 | T | C | 1.73E-06 | N | - |
| chr17: 45894107-45894607 | chr17:45894238 | C | G | 1.73E-06 | N | - |
| chr17: 45894107-45894607 | chr17:45894419 | A | G | 1.73E-06 | N | - |
| chr17: 45894107-45894607 | chr17:45894571 | C | A | 1.73E-06 | N | - |
| chr17: 46192430-46192951 | chr17:46192598 | T | A | 1.73E-06 | N | - |
| chr17: 46192430-46192951 | chr17:46192927 | G | T | 1.73E-06 | N | - |

| | | | | | | |
|---|---|---|---|---|---|---|
| chr17: 46192430-46192951 | chr17:46192693 | A | T | 1.73E-06 | Y | chr17: 46192430-46192921 |
| chr17: 46193103-46194451 | chr17:46193786 | G | A | 1.73E-06 | Y | chr17: 46193679-46194177 |
| chr17: 46193103-46194451 | chr17:46194064 | A | G | 1.73E-06 | Y | chr17: 46193841-46194375 |
| chr17: 4795850-4796452 | chr17:4795953 | G | T | 6.46E-07 | N | - |
| chr17: 4857952-4858551 | chr17:4858045 | G | T | 8.04E-08 | N | - |
| chr19: 1400528-1401067 | chr19:1400767 | G | A | 4.51E-07 | N | - |
| chr19: 2270358-2270901 | chr19:2270535 | T | C | 1.50E-07 | N | - |
| chr19: 44500276-44500951 | chr19:44500644 | G | C | 2.19E-07 | Y | chr19: 44500276-44500812 |
| chr19: 44500276-44500951 | chr19:44500647 | C | G | 2.19E-07 | Y | chr19: 44500276-44500812 |
| chr19: 44500276-44500951 | chr19:44500683 | G | C | 2.19E-07 | Y | chr19: 44500276-44500812 |
| chr19: 506946-507851 | chr19:507089 | C | A | 4.38E-07 | Y | chr19: 506946-507480 |
| chr19: 506946-507851 | chr19:507444 | T | TGG | 5.57E-08 | Y | chr19: 507100-507583 |
| chr19: 55117597-55118210 | chr19:55117865 | C | T | 2.82E-07 | Y | chr19: 55117734-55118210 |

| | | | | | | |
|---|---|---|---|---|---|---|
| chr2: 677149-677751 | chr2:677591 | C | G | 2.83E-06 | Y | chr2: 677230-677751 |
| chr22: 40856647-40857358 | chr22:40856958 | C | G | 7.29E-08 | Y | chr22: 40856647-40857078 |
| chr5: 892544-893044 | chr5:892731 | C | T | 5.84E-09 | N | - |
| chr5: 892544-893044 | chr5:892840 | G | A | 5.84E-09 | N | - |
| chr6: 149963675-149964175 | chr6:149964166 | G | A | 4.05E-08 | N | - |
| chr6: 18122440-18123016 | chr6:18122711 | G | C | 3.38E-08 | Y | chr6: 18122507-18123016 |
| chr7: 105531799-105532438 | chr7:105532059 | T | C | 8.98E-09 | Y | chr7: 105531891-105532438 |
| chr8: 142225856-142226399 | chr8:142226252 | G | T | 1.33E-08 | N | - |
| chr9: 97921849-97922683 | chr9:97922252 | A | G | 9.95E-08 | Y | chr9: 97921926-97922420 |

**Table S11.**

**Regions selected for MutSTARR-seq experiments**. Chromosome location lists the putative enhancer region, and eQTL location lists the chromosomal coordinates for the eQTL. The reference and alternate alleles refer to the wild-type and mutant alleles respectively. Alleles that include multiple base pairs represent insertions or deletions. Regions that had to be trimmed for quality control purposes due to repetitive regions are denoted with a “Y” in the “Region trimmed?” column, and the final regions used to generate the eBlocks can be found in the final column (“Trimmed Region”). Abbreviations: eQTL = quantitative trait locus.

Too large – provided as a separate file.

**Table S12.**

**Results from MutSTARR-seq experiment**. Regions overlapping eQTLs are indicated by their ID and chromosomal location. The “Expectation” column contains the input for the alternative allele divided by the input for the reference allele. The “Observation” columns contain the output for the alternative allele divided by the output for the reference allele across each technical replicate (1-4). P-values were determined using a Chi-square test and adjusted p-values were calculated using Bonferroni correction and Benjamini-Hochberg false discovery weight approaches.

Too large – provided as a separate file.

**Table S13.**
**Predicted target genes for significant region from MutSTARR-seq experiment.**

Genes predicted to be regulated by the significant region from the MutSTARR-seq experiment (chr17: 45,894,107-45,894,607; see Table S12). These genes were identified based on QTL analyses from (*30*). The SNP associated with the QTL is identified in the "QTL SNP ID" column. The chromosomal location of each SNP is provided as is the p-value for the QTL from the original paper (*30*). When performing eQTL analysis for a specific SNP, to assess the effect of different genotypes on expression, a linear-based model is run. The "QTL r2" represents the coefficient of determination of the best fit model. The "QTL beta slope" column contains the effect size of the genotype on gene expression. A larger (absolute) value represents a greater change in expression between reference and alternate genotype. A positive value indicates increased gene expression with the alternate allele, and a negative value indicates decreased expression. The "Cell Type" column states in which cell type that specific result was seen. Abbreviations: TSS = transcription start site, eQTL = expression quantitative trait locus, SNP = single nucleotide polymorphism.

| Enhancer ID | Band Label | Base Pairs (bp) | Adjusted Volume (band intensity) | Editing Efficiency |
|---|---|---|---|---|
| EH37E1198822 | WT band | 3375 | 135182913 | 23.19% |
| | genome edited band | 850 | 10279032 | |
| EH37E1000386 | WT band | 1938 | 231086083 | 25.12% |
| | genome edited band | 809 | 32352910 | |
| EH37E0114246 | WT band | 1849 | 22865778 | 19.69% |
| | genome edited band | 1205 | 3652503 | |
| EH37E0426064 | WT band | 3069 | 41081182 | 44.97% |
| | genome edited band | 2203 | 24099382 | |

**Table S14**.
**Genome editing knockout (KO) efficiency of CRISPR/Cas9 enhancer deletion.** The genome editing KO efficiency was calculated through densitometric analysis of DNA genotyping PCR gel image. The band intensities were analyzed using Image Labs software (Bio-Rad) by plotting the band intensities for each lane.

| Enhancer ID/guide name | Guide sequence | PAM | Location | Cutting site | KO deletion size (bp) |
|---|---|---|---|---|---|
| EH37E1198822. us2 | CGAGGAGCACCGG AGACTAT | TGG | chr2:232871313-232871332 | chr2:232871329 | 2525 bp |
| EH37E1198822. ds1 | GGGTGGTCTTCA AGGTTTGC | GGG | chr2:232873838-232873857 | chr2:232873854 | |
| EH37E1000386. us1 | TCCGGTGTGAC CAAAAGTAG | TGG | chr9:74727563-74727582 | chr9:74727579 | 1129 bp |
| EH37E1000386. ds1 | GTTCTACTAGAAT ACCCCAA | AGG | chr9:74728692-74728711 | chr9:74728708 | |
| EH37E0114246. us1 | CGCATAGTATGT GAGGACCA | GGG | chr1:150163018-150163037 | chr1:150163021 | 644 bp |
| EH37E0114246. ds1 | GGCTGCACTAAT TGACCTCG | AGG | chr1:150163662-150163681 | chr1:150163665 | |
| EH37E0426064. us1 | ATGTGAACTGCC TGCAACGG | TGG | chr17:17920923-17920942 | chr17:17920926 | 866 bp |
| EH37E0426064. ds1 | GACTGCACGAAGA GGCCGAG | AGG | chr17:17921789-17921808 | chr17:17921792 | |

**Table S15**.
**crRNA XT (gRNA) sequences, PAM sequences, Cas9 nuclease cutting sites and knockout (KO) deletion sizes**. gRNA sequence was designed in a 300 bp window of the 5’ or 3’ flanking regions of the enhancer with the IDT gRNA design algorithm. The PAM sequence is required for the Cas9 nuclease to cut and is generally found 3 nucleotides downstream from the Cas9 cut site. The distance between upstream and downstream Cas9 cutting sites is calculated as KO deletion size. Abbreviations: gRNA = guide RNA, PAM = protospacer adjacent motif.

| phNPC ATAC-seq Sample | hESC-derived NPC DNase-seq Sample | Jaccard Index |
|---|---|---|
| GSM2494716_ATAC-SeqID13 | ENCFF687UQV.bed | 0.18460003 |
| GSM2494717_ATAC-SeqID14 | ENCFF687UQV.bed | 0.18879659 |
| GSM2494718_ATAC-SeqID15 | ENCFF687UQV.bed | 0.18652245 |
| GSM2494719_ATAC-SeqID16 | ENCFF687UQV.bed | 0.19026071 |
| GSM2494723_ATAC-SeqID123 | ENCFF687UQV.bed | 0.19773698 |
| GSM2494724_ATAC-SeqID124 | ENCFF687UQV.bed | 0.1940287 |
| GSM2494725_ATAC-SeqID125 | ENCFF687UQV.bed | 0.19673102 |
| GSM2494729_ATAC-SeqID135 | ENCFF687UQV.bed | 0.20607529 |
| GSM2494730_ATAC-SeqID136 | ENCFF687UQV.bed | 0.19879302 |
| GSM2494731_ATAC-SeqID137 | ENCFF687UQV.bed | 0.20542265 |

**Table S16**.
**Comparison of open chromatin regions between phNPCs and hESC-derived NPCs**. Jaccard indices were calculated to determine the similarity between ATAC-seq data from phNPCs (*65*) and DNase-seq data from an hESC-derived NPC line (https://www.encodeproject.org/experiments/ENCSR278FVO/). DNase-seq data was only available for a single hESC-derived NPC sample, as shown in the "hESC-derived NPC DNase-seq Sample" column. ATAC-seq data was available for 10 different phNPC samples from the NPC-rich GZ zone. The specific sample names used are shown in the "phNPC ATAC-seq Sample" column. Jaccard indices were calculated comparing the hESC-derived NPC sample to each phNPC ATAC-seq sample. These Jaccard indices are visualized in the box plot in Fig. S3.

| Oligo | Purpose | Sequence |
|---|---|---|
| Adaptor I | Adaptor ligation | 5’ - /5Phos/GATCGGAAGAGCACACGTCT - 3’ |
| Adaptor II | Adaptor ligation | 5’ - ACACTCTTTCCCTACACGACGCTCTTCCGATCT - 3’ |
| MPI_ORI_F | LM-PCR forward primer | 5’ - TGATCTAGAGCATGCACCGGACACTCTTTC CCTACACGACGCTCTTCCGATCT - 3’ |
| MPI_ORI_R | LM-PCR reverse primer | 5’ - TCTAGCCTTCTCGTGTGCAGACTTGAGGTCA GTGAGCTGCTTAAGCCGGCCGGCG - 3’ |
| MPI_universa l | Illumina sequencing primer | 5’ - AATGATACGGCGACCACCGAGATCTAC ACTCTTTCCCTACACGACGCTCTTCCGATCT - 3’ |
| MPII_index_0 2 | Illumina sequencing primer | 5’ - CAAGCAGAAGACGGCATACGAGATACATC GGTGACTGGAGTTCAGACGTG - 3’ |

**Table S17**.
**Oligonucleotides for the input library generation**. Adaptors were used for the ligation step and primers were used for the LM-PCR. The /5Phos/ indicates a 5’ phosphorylation modification of Adaptor I. All oligos were ordered from Integrated DNA Technologies (IDT; Coralville, IA) and were ion-exchange-high-performance liquid chromatography (IE-HPLC) purified.

Too large – provided as a separate file.

**Table S18.**

**Candidate enhancers for CRISPR/Cas9 knockout (KO)**. The four candidate enhancer regions selected for CRISPR/Cas9 validation are displayed. CapSTARR-seq enrichment ratios are shown for replicate 1 (R1) and replicate 2 (R2) of Panel 1. The nearest genes for each enhancer region are provided, included the location of the transcription start site (TSS) for each gene and its transcript ID.

| Primer name | Primer sequence | chr | strand | start | end | WT amplicon size | genome editing amplicon size |
|---|---|---|---|---|---|---|---|
| EH37E1198822.gen.F | CGACGTCCTTAGAGAATGAGGACC | chr2 | + | 232870838 | 232870861 | 3375 bp | 850 bp |
| EH37E1198822.gen.R | ATGTTGGCAGTTGGCAAGACTAG | chr2 | - | 232874190 | 232874212 | | |
| EH37E1000386.gen.F | ACAAGTCCGGTGTGACCAAA | chr9 | + | 74727558 | 74727577 | 1938 bp | 809 bp |
| EH37E1000386.gen.R | CTACTCGCCTCGGAGCAAAG | chr9 | - | 74729476 | 74729495 | | |
| EH37E0114246.gen.F | AGAGGAGAGAGCCAGAAGGG | chr1 | + | 150162246 | 150162265 | 1849 bp | 1205 bp |
| EH37E0114246.gen.R | ACCCCAACTTGTTCACAGCA | chr1 | - | 150164075 | 150164094 | | |
| EH37E0426064.gen.F | CGGTGTTGTTGCAAGTTCCC | chr17 | + | 17919662 | 17919681 | 3069 bp | 2203 bp |
| EH37E0426064.gen.R | CCAGGTTCCCTGTTTCGTGT | chr17 | - | 17922711 | 17922730 | | |

**Table S19**.

**DNA genotyping PCR primer sequences and amplicon sizes**. The primer pairs were designed in the flanking sequence of the 5'-cutting site or 3'-cutting site. The calculation formula: the genome editing amplicon size (bp) = WT amplicon size (bp) - KO deletion size (bp, see Table S15).

| IDT Assay ID | Gene | Species | Ref Seq # | Detects All Variants | Exon Location | Forward primer | Reverse primer | Probe | PCR amplification efficiency |
|---|---|---|---|---|---|---|---|---|---|
| Hs.PT.58.20686686 | NGEF | Homo_sapiens | NM_001114090 | Yes | 15 - 16 | TCA AGA TCT CCT CAG TCA TGG A | GCC GAC ATC CTC AAC ATC C | /56-FAM/AG ACG CTC G/ZEN/C CAA AGA TCC ACC /3IABkFQ | 106.28 % |
| Hs.PT.58.15573952 | RORB | Homo_sapiens | NM_006914 | Yes | 5 - 6 | TGC AAA CTC CAC CAC GTA T | GCA GAC CCA CAC CTA TGA AG | /56-FAM/AG CAT ATC A/ZEN/A AGC AAG TCC AGG GAA GC/3IABkFQ/ | 108.68 % |
| Hs.PT.58.40931620 | PLEKHO1 | Homo_sapiens | NM_016274 | Yes | 5 - 6 | TCA AGT CCA AGG TCA GCA TC | ACA GCT ATC TTG CCC ATC C | /56-FAM/CA AAA ATC C/ZEN/A GCA CTC CCG CCG /3IABkFQ/ | 99.66 % |
| Hs.PT.58.2267363 | TOM1L2 | Homo_sapiens | NM_001033551 | Yes | 8 - 10 | CAG GTC TAT TAA GTT GTC TTC GGT | ACC TCA ACA ACG TCT TCC TTC | /56-FAM/AT TAC TGG C/ZEN/A TTT TGA ACG GAT CGG C/3IABkFQ/ | 99.66 % |

| | | | | | | | | | |
|---|---|---|---|---|---|---|---|---|---|
| Hs.PT.39a.22214847 | ACTB | Homo_sapiens | NM_001101 | Yes | 1 - 2 | CCT TGC ACA TGC CGG AG | ACA GAG CCT CGC CTT TG | /56-FAM/TC ATC CAT G/ZEN/G TGA GCT GGC GG/3IABkFQ/ | 88.57 % |

**Table S20**.

**Sequences of TaqMan qPCR primers and probes**. The IDT Assay ID designates the predesigned PrimeTime qPCR Probe Assay available from IDT. The double quenched qPCR probe is labeled with 5’ FAM fluorophore dye, a ZEN internal quencher and the 3’ Iowa Black FQ quencher. Each Probe Assay was first tested with qPCR to confirm an amplification efficiency between 88%~110%.

| Well Position | Sample Name | Target Name | CT | Ct Mean | Ct SD |
|---|---|---|---|---|---|
| A1 | EH37E1198822 KO BR1 | NGEF | 31.289 | 31.377 | 0.151 |
| A2 | EH37E1198822 KO BR1 | NGEF | 31.552 | 31.377 | 0.151 |
| A3 | EH37E1198822 KO BR1 | NGEF | 31.291 | 31.377 | 0.151 |
| A7 | EH37E1198822 KO BR1 | ACTB | 22.908 | 23.056 | 0.134 |
| A8 | EH37E1198822 KO BR1 | ACTB | 23.169 | 23.056 | 0.134 |
| A9 | EH37E1198822 KO BR1 | ACTB | 23.09 | 23.056 | 0.134 |
| B1 | EH37E1198822 KO BR2 | NGEF | 30.745 | 30.915 | 0.148 |
| B2 | EH37E1198822 KO BR2 | NGEF | 31.02 | 30.915 | 0.148 |
| B3 | EH37E1198822 KO BR2 | NGEF | 30.979 | 30.915 | 0.148 |
| B7 | EH37E1198822 KO BR2 | ACTB | 22.868 | 22.664 | 0.189 |
| B8 | EH37E1198822 KO BR2 | ACTB | 22.495 | 22.664 | 0.189 |
| B9 | EH37E1198822 KO BR2 | ACTB | 22.629 | 22.664 | 0.189 |
| A13 | Control: cells, +e BR1 | NGEF | 30.207 | 30.112 | 0.106 |
| A14 | Control: cells, +e BR1 | NGEF | 29.998 | 30.112 | 0.106 |
| A15 | Control: cells, +e BR1 | NGEF | 30.13 | 30.112 | 0.106 |
| A22 | Control: cells, +e BR1 | ACTB | 22.993 | 22.976 | 0.025 |
| C22 | Control: cells, +e BR1 | ACTB | 22.973 | 22.976 | 0.025 |
| E22 | Control: cells, +e BR1 | ACTB | 22.934 | 22.976 | 0.025 |
| G22 | Control: cells, +e BR1 | ACTB | 22.991 | 22.976 | 0.025 |
| I22 | Control: cells, +e BR1 | ACTB | 22.99 | 22.976 | 0.025 |
| B13 | Control: cells, +e BR2 | NGEF | 29.875 | 29.681 | 0.277 |
| B14 | Control: cells, +e BR2 | NGEF | 29.804 | 29.681 | 0.277 |
| B15 | Control: cells, +e BR2 | NGEF | 29.364 | 29.681 | 0.277 |
| B22 | Control: cells, +e BR2 | ACTB | 22.508 | 22.657 | 0.139 |
| D22 | Control: cells, +e BR2 | ACTB | 22.68 | 22.657 | 0.139 |
| J22 | Control: cells, +e BR2 | ACTB | 22.783 | 22.657 | 0.139 |

**Table S21.**
**Ct value raw data of TaqMan qPCR assay for *NGEF***. qPCR was performed using the QuantStudio 7 Flex Real-Time PCR System (Applied Biosystems). The quantification threshold cycles (CT) were calculated using the default settings in the QuantStudio Real Time PCR Software v1.3 (Applied Biosystems).

| Well Position | Sample Name | Target Name | CT | Ct Mean | Ct SD |
|---|---|---|---|---|---|
| E1 | EH37E100386 KO BR1 | RORB | 30.579 | 30.564 | 0.078 |
| E2 | EH37E100386 KO BR1 | RORB | 30.479 | 30.564 | 0.078 |
| E3 | EH37E100386 KO BR1 | RORB | 30.633 | 30.564 | 0.078 |
| E7 | EH37E100386 KO BR1 | ACTB | 21.766 | 21.8 | 0.072 |
| E8 | EH37E100386 KO BR1 | ACTB | 21.751 | 21.8 | 0.072 |
| E9 | EH37E100386 KO BR1 | ACTB | 21.882 | 21.8 | 0.072 |
| F1 | EH37E1000386 KO BR2 | RORB | 31.349 | 31.318 | 0.03 |
| F2 | EH37E1000386 KO BR2 | RORB | 31.288 | 31.318 | 0.03 |
| F3 | EH37E1000386 KO BR2 | RORB | 31.316 | 31.318 | 0.03 |
| F7 | EH37E1000386 KO BR2 | ACTB | 21.801 | 21.839 | 0.049 |
| F8 | EH37E1000386 KO BR2 | ACTB | 21.824 | 21.839 | 0.049 |
| F9 | EH37E1000386 KO BR2 | ACTB | 21.894 | 21.839 | 0.049 |
| E13 | Control: cells, +e BR1 | RORB | 30.232 | 30.4 | 0.236 |
| E14 | Control: cells, +e BR1 | RORB | 30.298 | 30.4 | 0.236 |
| E15 | Control: cells, +e BR1 | RORB | 30.669 | 30.4 | 0.236 |
| F13 | Control: cells, +e BR2 | RORB | 30.123 | 30.351 | 0.222 |
| F14 | Control: cells, +e BR2 | RORB | 30.366 | 30.351 | 0.222 |
| F15 | Control: cells, +e BR2 | RORB | 30.566 | 30.351 | 0.222 |
| A22 | Control: cells, +e BR1 | ACTB | 22.003 | 22.146 | 0.086 |
| C22 | Control: cells, +e BR1 | ACTB | 22.128 | 22.146 | 0.086 |
| E22 | Control: cells, +e BR1 | ACTB | 22.214 | 22.146 | 0.086 |
| G22 | Control: cells, +e BR1 | ACTB | 22.18 | 22.146 | 0.086 |
| I22 | Control: cells, +e BR1 | ACTB | 22.203 | 22.146 | 0.086 |
| B22 | Control: cells, +e BR2 | ACTB | 22.277 | 22.139 | 0.138 |
| D22 | Control: cells, +e BR2 | ACTB | 22.193 | 22.139 | 0.138 |
| F22 | Control: cells, +e BR2 | ACTB | 22.066 | 22.139 | 0.138 |
| H22 | Control: cells, +e BR2 | ACTB | 21.935 | 22.139 | 0.138 |
| J22 | Control: cells, +e BR2 | ACTB | 22.222 | 22.139 | 0.138 |

**Table S22.**
**Ct value raw data of TaqMan qPCR assay for *RORB***. qPCR was performed using the QuantStudio 7 Flex Real-Time PCR System (Applied Biosystems). The quantification threshold cycles (CT) were calculated using the default settings in the QuantStudio Real Time PCR Software v1.3 (Applied Biosystems).

| Well Position | Sample Name | Target Name | CT | Ct Mean | Ct SD |
|---|---|---|---|---|---|
| C1 | EH37E0114246 KO BR1 | PLEKHO1 | 29.176 | 29.302 | 0.111 |
| C2 | EH37E0114246 KO BR1 | PLEKHO1 | 29.343 | 29.302 | 0.111 |
| C3 | EH37E0114246 KO BR1 | PLEKHO1 | 29.387 | 29.302 | 0.111 |
| C7 | EH37E0114246 KO BR1 | ACTB | 22.114 | 22.152 | 0.032 |
| C8 | EH37E0114246 KO BR1 | ACTB | 22.172 | 22.152 | 0.032 |
| C9 | EH37E0114246 KO BR1 | ACTB | 22.169 | 22.152 | 0.032 |
| D1 | EH37E0114246 KO BR2 | PLEKHO1 | 28.357 | 28.578 | 0.274 |
| D2 | EH37E0114246 KO BR2 | PLEKHO1 | 28.884 | 28.578 | 0.274 |
| D3 | EH37E0114246 KO BR2 | PLEKHO1 | 28.492 | 28.578 | 0.274 |
| D7 | EH37E0114246 KO BR2 | ACTB | 21.026 | 21.118 | 0.08 |
| D8 | EH37E0114246 KO BR2 | ACTB | 21.156 | 21.118 | 0.08 |
| D9 | EH37E0114246 KO BR2 | ACTB | 21.173 | 21.118 | 0.08 |
| C13 | Control: cells, +e BR1 | PLEKHO1 | 26.646 | 26.816 | 0.219 |
| C14 | Control: cells, +e BR1 | PLEKHO1 | 26.738 | 26.816 | 0.219 |
| C15 | Control: cells, +e BR1 | PLEKHO1 | 27.063 | 26.816 | 0.219 |
| D13 | Control: cells, +e BR2 | PLEKHO1 | 26.471 | 26.737 | 0.239 |
| D14 | Control: cells, +e BR2 | PLEKHO1 | 26.804 | 26.737 | 0.239 |
| D15 | Control: cells, +e BR2 | PLEKHO1 | 26.934 | 26.737 | 0.239 |
| A22 | Control: cells, +e BR1 | ACTB | 22.003 | 22.146 | 0.086 |
| C22 | Control: cells, +e BR1 | ACTB | 22.128 | 22.146 | 0.086 |
| E22 | Control: cells, +e BR1 | ACTB | 22.214 | 22.146 | 0.086 |
| G22 | Control: cells, +e BR1 | ACTB | 22.18 | 22.146 | 0.086 |
| I22 | Control: cells, +e BR1 | ACTB | 22.203 | 22.146 | 0.086 |
| B22 | Control: cells, +e BR2 | ACTB | 22.277 | 22.139 | 0.138 |
| D22 | Control: cells, +e BR2 | ACTB | 22.193 | 22.139 | 0.138 |
| F22 | Control: cells, +e BR2 | ACTB | 22.066 | 22.139 | 0.138 |
| H22 | Control: cells, +e BR2 | ACTB | 21.935 | 22.139 | 0.138 |
| J22 | Control: cells, +e BR2 | ACTB | 22.222 | 22.139 | 0.138 |

**Table S23.**
**Ct value raw data of TaqMan qPCR assay for *PLEKHO1***. qPCR was performed using the QuantStudio 7 Flex Real-Time PCR System (Applied Biosystems). The quantification threshold cycles (CT) were calculated using the default settings in the QuantStudio Real Time PCR Software v1.3 (Applied Biosystems).

| Well Position | Sample Name | Target Name | CT | Ct Mean | Ct SD | Note |
|---|---|---|---|---|---|---|
| E1 | EH37E0426064 KO BR1 | TOM1L2 | 28.472 | 28.553 | 0.115 | |
| E2 | EH37E0426064 KO BR1 | TOM1L2 | 28.634 | 28.553 | 0.115 | |
| E3 | EH37E0426064 KO BR1 | TOM1L2 | 29.253 | | | outlier, removed from Ct Mean calculation |
| E7 | EH37E0426064 KO BR1 | ACTB | 23.742 | | | outlier, removed from Ct Mean calculation |
| E8 | EH37E0426064 KO BR1 | ACTB | 22.452 | 22.08 | 0.526 | |
| E9 | EH37E0426064 KO BR1 | ACTB | 21.708 | 22.08 | 0.526 | |
| F1 | EH37E0426064 KO BR2 | TOM1L2 | 28.436 | 28.3 | 0.171 | |
| F2 | EH37E0426064 KO BR2 | TOM1L2 | 28.355 | 28.3 | 0.171 | |
| F3 | EH37E0426064 KO BR2 | TOM1L2 | 28.108 | 28.3 | 0.171 | |
| F7 | EH37E0426064 KO BR2 | ACTB | 22.033 | 21.857 | 0.25 | |
| F8 | EH37E0426064 KO BR2 | ACTB | 21.681 | 21.857 | 0.25 | |
| F9 | EH37E0426064 KO BR2 | ACTB | 21.067 | | | outlier, removed from Ct Mean calculation |
| E13 | Control: cells, +e BR1 | TOM1L2 | 28.021 | 27.81 | 0.196 | |
| E14 | Control: cells, +e BR1 | TOM1L2 | 27.775 | 27.81 | 0.196 | |
| E15 | Control: cells, +e BR1 | TOM1L2 | 27.634 | 27.81 | 0.196 | |
| F13 | Control: cells, +e BR2 | TOM1L2 | 27.519 | 27.458 | 0.083 | |
| F14 | Control: cells, +e BR2 | TOM1L2 | 27.364 | 27.458 | 0.083 | |
| F15 | Control: cells, +e BR2 | TOM1L2 | 27.492 | 27.458 | 0.083 | |

| | | | | | |
|---|---|---|---|---|---|
| A22 | Control: cells, +e BR1 | ACTB | 22.567 | 22.595 | 0.094 |
| C22 | Control: cells, +e BR1 | ACTB | 22.72 | 22.595 | 0.094 |
| E22 | Control: cells, +e BR1 | ACTB | 22.461 | 22.595 | 0.094 |
| G22 | Control: cells, +e BR1 | ACTB | 22.628 | 22.595 | 0.094 |
| I22 | Control: cells, +e BR1 | ACTB | 22.598 | 22.595 | 0.094 |
| B22 | Control: cells, +e BR2 | ACTB | 22.45 | 22.404 | 0.151 |
| D22 | Control: cells, +e BR2 | ACTB | 22.65 | 22.404 | 0.151 |
| F22 | Control: cells, +e BR2 | ACTB | 22.328 | 22.404 | 0.151 |
| H22 | Control: cells, +e BR2 | ACTB | 22.295 | 22.404 | 0.151 |
| J22 | Control: cells, +e BR2 | ACTB | 22.298 | 22.404 | 0.151 |

**Table S24.**
**Ct value raw data of TaqMan qPCR assay for *TOM1L2***. qPCR was performed using the QuantStudio 7 Flex Real-Time PCR System (Applied Biosystems). The quantification threshold cycles (CT) were calculated using the default settings in the QuantStudio Real Time PCR Software v1.3 (Applied Biosystems).
